# Large-scale all-optical electrophysiology links excitability and connectivity to circuit dynamics in vivo

**DOI:** 10.64898/2026.10.05.756595

**Authors:** Yitong Qi, Vicente Parot, Xiang Wu, Sarah E. Plutkis, Luke D. Lavis, Adam E. Cohen

**Affiliations:** Department of Chemistry and Chemical Biology, Harvard University, Cambridge, MA 02138; Universidad Catolica, Santiago, Chile; Janelia Farm Research Campus, Howard Hughes Medical Institute, Ashburn, VA; Department of Physics, Harvard University, Cambridge, MA 02138

## Abstract

To understand neural computations, one would like to relate circuit dynamics to neuronal intrinsic excitability and connectivity. We developed tools for large-scale all-optical electrophysiology and applied them to approximately 300 layer 1 interneurons per mouse *in vivo*. Across two mice, targeted stimulation and voltage imaging identified 1,469 inhibitory and 5,752 directed electrical connections among 14,312 tested pairs. Presynaptic identity was more strongly associated with inhibitory post-synaptic potential (IPSP) amplitude, whereas postsynaptic identity was more strongly associated with IPSP decay time. Gap junctional and synaptic interactions had distinct spatiotemporal kernels, with corresponding signatures in spontaneous activity. Excitability measured under optogenetic stimulus predicted spiking responses to subthreshold voltage, depolarization rate, and spike history during spontaneous firing. Genetic subtypes also showed different behavioral-state-dependent voltage-to-spike relationships despite similar population-average subthreshold voltages. These measurements link cellular properties and pairwise interactions to spontaneous circuit dynamics, establishing an approach for functional circuit mapping *in vivo*.

**Highlights:**

- All-optical mapping links excitability, connectivity, and spontaneous dynamics in vivo
- Presynaptic and postsynaptic identity relate differently to IPSP strength and decay
- Gap junctional and synaptic couplings reveal distinct spatiotemporal interactions
- Measured intrinsic excitability predicts features of spontaneous spiking in vivo

## Introduction

A fundamental aim of neuroscience is to determine how brain circuits transform their inputs to their outputs: how intrinsic excitability, local synaptic and gap junctional connections, and genetic cell-type identity interact to convert patterns of subthreshold inputs to spiking outputs. To answer this question, one must measure all these properties within the same circuit in a behaving animal, a combination that has been difficult to achieve at population scale. Anatomical connectomics resolves wiring but reports neither excitability nor circuit dynamics. Correlational analysis of spike trains reports circuit output, but confounds monosynaptic coupling with shared input or recurrent interactions^1^. Patch clamp measurements resolve subthreshold voltage and intrinsic properties directly, but do so in only one or two cells at a time and are difficult to combine with behavior. One would like to independently perturb and measure the membrane voltages of hundreds of genetically identified neurons in behaving animals.

Advances in genetically encoded voltage indicators (GEVIs)^2–5^ and microscopes^6^ have enabled voltage imaging of up to ∼300 neurons *in vivo*^7–9^. However, combined targeted optogenetic stimulation and voltage imaging have only been demonstrated for small ensembles, due to unavailability of spectrally orthogonal and bright actuator-indicator pairs^10–14^. To determine how network function emerges from single-neuron properties and pairwise interactions requires perturbation and measurement at large scale.

Cortical Layer 1 (L1) integrates top-down cortico-cortical inputs, bottom-up sensory inputs, and neuromodulatory inputs^15,16^. Its sparse interneuron population inhibits presynaptic axons^17^, apical dendrites^18^ and other deeper layer interneurons^19,20^, and forms a mutually inhibitory network^21^ that is also coupled by gap junctions^22^. This circuit gates activity in the underlying cortex^23^ and plays a critical role in modulating learning^24–26^ and perception^27^, but the collective dynamics, the computational function, and the roles of the genetically distinct sub-populations of the L1 microcircuit remain little understood. Most L1 interneurons express NDNF, and this population divides into two genetically defined types: NPY-expressing neurogliaform cells, and NPY-negative canopy cells^21^. L1 is thin, sparse, and is genetically and optically accessible, making it a favorable place to attempt a functional map of a microcircuit.

Here, we developed chemigenetic tools, optical instrumentation, and software for highly parallel all-optical electrophysiology in behaving mice. The platform combined voltage imaging at ∼800 Hz with simultaneous targeted optogenetic stimulation in ensembles of ∼300 L1 interneurons in the barrel cortex. We linked measurements of intrinsic excitability, pairwise inhibitory and electrical coupling, spontaneous subthreshold and spiking activity, and whisker motion, across two mice.

These joint measurements revealed relationships between cellular properties, pairwise interactions, and spontaneous circuit dynamics. Presynaptic identity contributed more to the amplitudes of inhibitory postsynaptic potentials (IPSPs) and postsynaptic identity contributed more to the time constants. Gap junctional and synaptic coupling showed distinct spatiotemporal kernels and were associated with rapid synchrony and delayed spike suppression, respectively, during spontaneous activity. Independently measured optical rheobase, spike-rate adaptation, and after-depolarization predicted sensitivities of the spontaneous spiking to subthreshold voltage, depolarization rate, and spike history, respectively.

Population voltage recordings also revealed broadly correlated subthreshold activity and traveling waves, alongside more localized and more transient spike synchrony. NPY-positive and NPY-negative populations had similar average subthreshold voltages but dramatically different spiking responses to whisking. These differences were traced to opposing behavioral state-dependent changes in the voltage-to-spiking relationships. Together, these findings illustrate how all-optical measurements of excitability, connectivity, and voltage dynamics can connect multiple levels of circuit function *in vivo*.

### Molecular and optical tools for wide-area all-optical electrophysiology

In previous Optopatch constructs, the voltage-sensitive microbial rhodopsin was very dim, requiring high-intensity laser illumination (> 10 W/mm^2^) and a high numerical aperture (NA > 0.7) objective^12–14^, which together constrained measurements to < 10 neurons simultaneously. The chemigenetic GEVIs Voltron1 and Voltron2 are ∼500-fold brighter than the pure opsin-based GEVIs^28^, leveraging fluorescence from a palette of bright and stable organic HaloTag-Ligand dyes^2,29^. We sought a bioavailable, red-shifted dye to combine with a blue-shifted channelrhodopsin, CheRiff^30^. While the commonly used Janelia Fluor (JF) 585 HaloTag ligand (HTL) is viable *in vivo*, we found the redder JF608-HTL to be 3.6 times brighter, with similar voltage sensitivity, leading to a 50% increase in signal-to-noise ratio (SNR) *in vivo* (Supplementary Fig. 1A-E). We developed an Optopatch construct for Cre-dependent co-expression of soma-targeted channelrhodopsin, CheRiff, and soma-targeted Voltron1 (Fig. 1A). An analogous construct made with Voltron2 performed similarly in our experiments.

**Figure 1.**
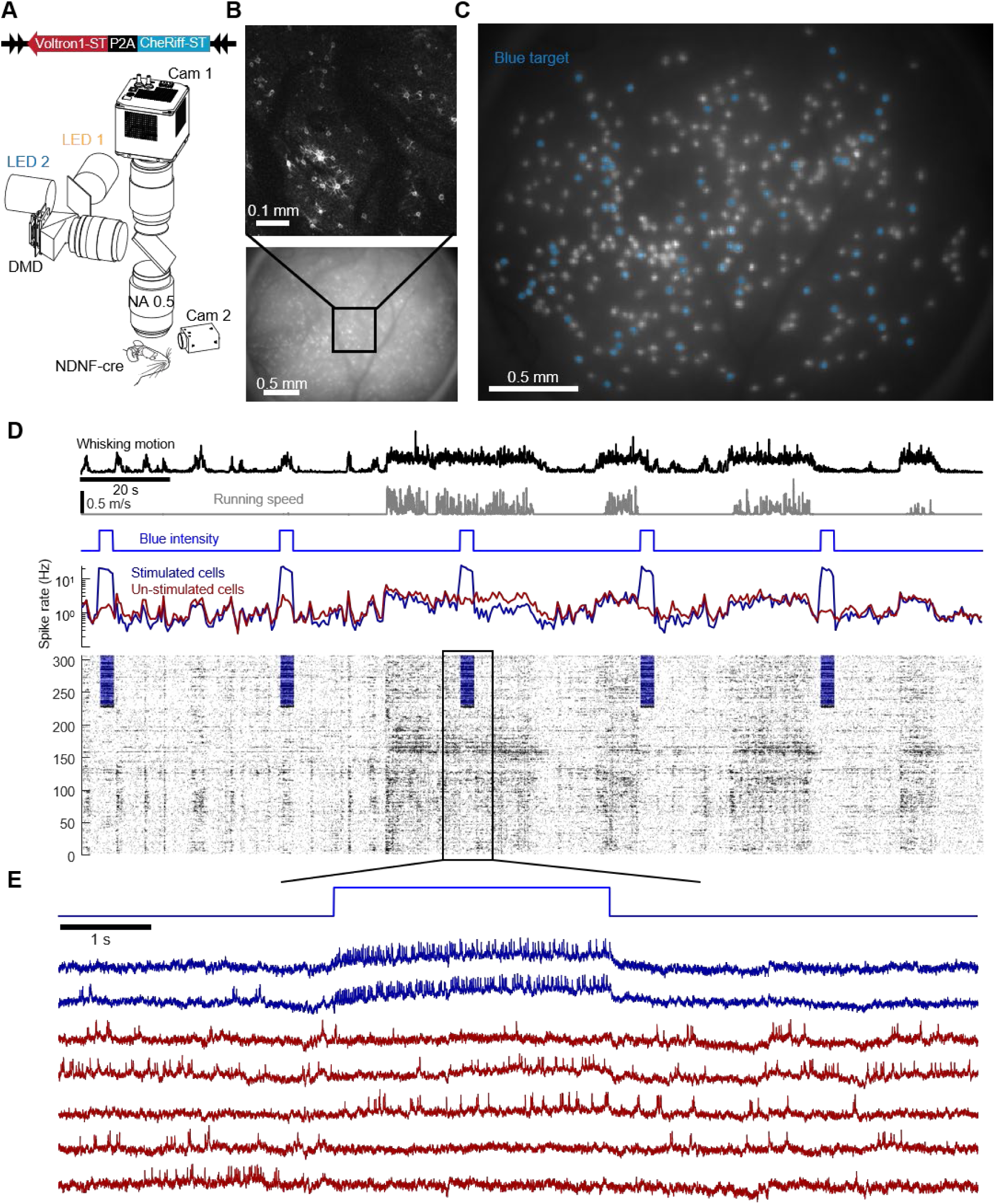
Molecular and optical systems for wide-area all-optical electrophysiology. (A) Top: molecular construct for CRE-dependent simultaneous expression of voltage indicator, Voltron1, and channelrhodopsin, CheRiff, where -ST denotes a soma-targeting motif. Bottom: optical system for simultaneous wide-area high-speed voltage imaging and optogenetic stimulation. (B) L1 interneurons expressing Optopatch in barrel cortex of an NDNF-Cre^+/-^ mouse acquired using (bottom) 1x wide-field imaging or (top) 7.5x Hadamard microscopy. (C) Same FOV as in (B) with soma-targeted illumination. 76 cells randomly selected for optogenetic stimulation in (D) are marked in blue. (D) Simultaneous behavior monitoring, voltage imaging, and optogenetic stimulation. Top: facial videography and locomotion were used to measure whisking and running speed. Middle: 76 cells were randomly selected and stimulated for 3-s epochs, repeated every 40 s. Stimulation increased the average firing rate of the stimulated cells (blue) 16-fold, and of the un-stimulated cells (red) 1.4-fold. Bottom: spike raster of all simultaneously measured cells. Stimulation marked by blue shading. (E) Example voltage traces. Blue: stimulated cells; red: un-stimulated cells.

We adapted an ultrawide-field imaging system^31^, to achieve an imaging field of view of 2 x 3 mm^2^ at 787 Hz (Fig. 1A; Supplementary Fig. 2; Methods). Blue and red high-power LEDs provided optogenetic stimulation and voltage imaging, respectively (Fig. 1C, Supplementary Fig. 1G; Methods). We combined both light sources onto a single digital micromirror device (DMD) (Fig. 1A; Supplementary Fig. 2), and synchronized the DMD patterns, the LED drive currents, and the camera frame trigger so each camera exposure (1.27 ms) was divided into an epoch (250 μs) for targeted optogenetic stimulation and an epoch (1000 μs) for structured illumination voltage imaging, each with a distinct DMD pattern (Supplementary Fig. 3; Methods). CheRiff has negligible photocurrents at wavelengths ≥ 594 nm^32^, and the small blue-light crosstalk into the voltage recordings was corrected during analysis (Methods).

We expressed the Optopatch construct in somatosensory cortex of NDNF-Cre mice, which drove expression in L1 interneurons, and imaged neural activity through a 3 mm-diameter cortical window. The mice were head-fixed on a linear treadmill and allowed to run freely. In addition to neural activity, we recorded synchronous locomotion and face videos.

Under wide-field red illumination, the individual neurons were difficult to resolve against the high fluorescence background (Fig. 1B). We used digital confocal Hadamard microscopy^33^ to make an optically sectioned image of the cortex, and then targeted the red light onto 320 neuronal somas found in the digital confocal image (Supplementary Fig. 4). This targeting increased the signal-to-background ratio ∼7.5-fold, from 0.15 ± 0.05 to 1.12 ± 0.32 (mean ± s.d., *n* = 320 cells, Fig. 1C, Supplementary Fig. 5). Movie 1 shows the voltage in *n* = 320 cells and the simultaneously recorded mouse face motion. High-resolution confocal z-stacks (Supplementary Fig. 6; Methods) showed that we recorded from ∼20% of the neurons in a focal plane of depth 50 μm and thickness ∼40 μm, and ∼12% (320/2,702) of all the NDNF-positive L1 interneurons within the cylinder bounded by the field of view (2.6 mm diameter and 170 μm thick). The density of NDNF-positive L1 interneurons was 2,860/mm^3^, consistent with previous estimates^34^.

The single-spike SNR (spike-height:baseline shot noise) was 4.5 ± 0.7 (mean ± s.d.). Using a fitted mixture model, we estimated the spike-detection false-discovery proportion to be 17 ± 13% across all neurons, and false-negative rate to be 28 ± 19% (Supplementary Fig. 7). After 1 h of continuous voltage imaging at 16 mW/mm^2^, the spike SNR decreased to 76 ± 13% of its initial value (Supplementary Fig. 8). To facilitate comparisons between cells, we normalized the signal for each cell by its mean initial spike height (Methods). Thus, all fluorescence signals are reported in units of Spike Height (SH). This measure does not readily convert to mV on account of the slightly nonlinear and non-instantaneous response of the voltage indicator and finite exposure time of the camera, which together blunted spike heights relative to slower subthreshold voltages.

To demonstrate targeted optogenetic perturbations, we periodically stimulated a randomly selected subset of 76 cells with blue light, while recording the voltage activity of all cells (Fig. 1C-E, Movie 2). All cells showed whisking and running modulated activity, while the stimulated subset showed on average a 16-fold increase in firing rate upon optogenetic stimulation. Non-stimulated cells showed only a slight (1.4-fold) increase in spike rate during stimulus epochs. To characterize this off-target stimulation, we laterally displaced the stimulation spots from the somas and found the excitatory response decayed with length constants of 47 and 25 μm for subthreshold voltage and firing rate, respectively (Supplementary Fig. 9). Since we continuously recorded voltages from stimulated and non-stimulated cells, we could incorporate the effects of this off-target stimulation during analysis, as described below.

### Circuit dissection by all-optical electrophysiology

We next characterized the intrinsic excitability, synaptic connectivity and gap junction connectivity of all recorded cells and eligible nearby cell pairs, using stimulus protocols designed to highlight each of these features of the microcircuit.

#### Intrinsic excitability

To measure intrinsic excitability, we stimulated sparse subsets of cells in awake mice with four 500-ms steps of blue light of increasing intensity, followed by a 5 s linear ramp, analogous to a stimulation sequence commonly used in patch clamp measurements^35–37^ (Fig. 2A). To minimize perturbations to network dynamics, each trial comprised ≤14 stimulated cells and each cell was > 800 μm from the nearest simultaneously stimulated neighbor (Supplementary Fig. 10A). L1 interneurons have an axonal spread of ∼300 μm and a dendritic spread of ∼150 μm^20,21^, so cells separated by > 800 μm were not likely to have direct connections. Over a total of 300 trials, each of the 320 cells received the stimulus protocol 7-10 times.

**Figure 2.**
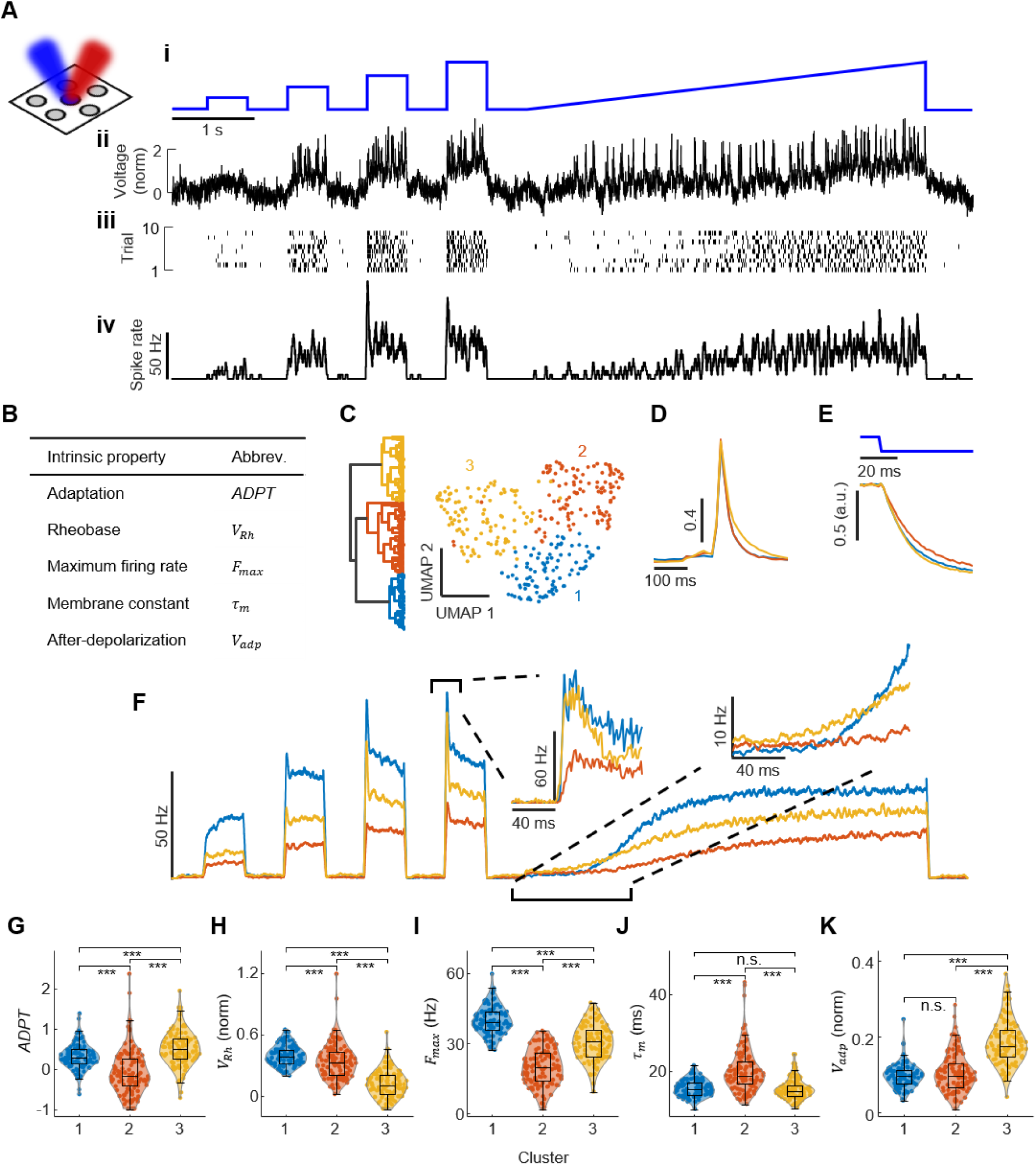
L1 interneurons show diverse intrinsic excitability *in vivo*. (A) Measurement protocol. **i**, stimulation waveform used to measure intrinsic excitability. **ii**, single trial response. **iii**, spike raster of all trials of the same cell in **ii**. **iv** trial-average firing rate of the same cell. (B) Intrinsic properties extracted from each neuron, and the abbreviations used throughout. (C) 2-dimensional UMAP embedding of five intrinsic properties of 320 cells from one mouse (right panel). We divided these cells into 3 bins via hierarchical clustering (left panel), n = 94, 116, and 110 for clusters 1, 2, and 3 respectively. (D) Cluster-average spike waveform using only blue-light-evoked spikes. (E) Cluster-average voltage recovery upon blue light turn-off. (F) Cluster-average mean firing rate. (G-K) Distributions of the five intrinsic properties for the three functionally defined clusters: spike-rate adaptation (G), optical rheobase (H), maximum firing rate (I), membrane time constant (J), and spike after-depolarization (K). The clusters were defined by hierarchical clustering on these same five properties, so these comparisons are descriptive rather than inferential. Double-sided Wilcoxon rank sum test, * 0.01 < *p* ≤ 0.05, ** 0.001 < *p* ≤ 0.01, *** *p* ≤ 0.001.

We extracted the subthreshold and spiking dynamics and calculated five excitability-related properties for each cell (Fig. 2B; Supplementary Fig. 10B-D). From the step-stimulus responses, we measured spike-rate adaptation. From the slow ramp, we determined the subthreshold depolarization at the tenth spike during the ramp (our operational measure of optical rheobase), and the maximum firing rate under strong stimulus. From the subthreshold return to baseline at the end of the blue stimuli, we extracted a membrane time constant. Finally, from the average spike waveform, we extracted a measure of spike after-depolarization, V_adp._ These parameters were designed to be relatively insensitive to the CheRiff expression level, which could vary between cells.

We repeated this protocol on two mice and saw similar broad distributions of intrinsic properties in both animals (Supplementary Fig. 10E,H). Some properties were weakly correlated with others. For instance, firing rate adaptation was positively correlated with spike after-depolarization (V_adp;_ Supplementary Fig. 10E, R = 0.23, p = 1.9E-8, Pearson correlation, *n* = 564 neurons, 2 mice), possibly due to bursting induced by high V_adp_ at the onset of a step-stimulus.

The broad distribution of intrinsic properties is consistent with prior measurements in acute slices^38^, and suggests that the intrinsic properties vary along a continuum within the NDNF^+^ population. The 2D UMAP^39^ embedding of intrinsic properties (Fig. 2C) also showed a continuous distribution. To visualize the variations in intrinsic properties, we nonetheless divided the population into three bins via hierarchical clustering, then calculated the per-bin average stimulation-evoked responses (Fig. 2C-F). The three bins illustrate the range of variability in NDNF^+^ interneuron properties (Fig. 2G-K). Bin 1 had both the highest firing rate and the highest optical rheobase (i.e. it required the greatest depolarization to begin spiking); bin 2 had the lowest firing rate and lowest spike-rate adaptation, and the highest membrane time constant; bin 3 showed the highest after-depolarization and spike-rate adaptation and the lowest optical rheobase.

#### Synaptic inhibition

We then studied the interactions between cells. The most common way to infer synaptic connectivity *in vivo* is through the cross-correlogram (CCG) of spontaneous spike trains^40–42:^ a transient change in the spike rate of a putative post-synaptic neuron upon arrival of a spike in a putative presynaptic neuron is interpreted as a functional connection^43,44^. However, firing rate covariations due to shared inputs can also produce such features in the CCG^1,44^. Thus, this method is typically used with stringent thresholds for calling putative connections, which unavoidably misses many connections^42^.

In principle, spike-triggered cross averages of subthreshold voltages could provide a more sensitive measure of connectivity: the subthreshold voltage can report a response for every presynaptic spike, while spike CCGs rely on the (comparatively rare) co-occurrence of spikes in both cells. We calculated the spike-triggered cross averages of spontaneous voltage (Supplementary Fig. 11,12), i.e. the mean voltage waveform of cell *i*, triggered off spike times in cell *j*. These cross averages were also dominated by shared subthreshold fluctuations (i.e. the fluctuations which caused cell *i* to spike also depolarized cell *j*), which obscured the influence of direct *i* → *j* synaptic connections.

To measure synaptic connectivity directly, we thus turned to targeted optogenetic stimulation. We again sequentially stimulated sparse subsets of cells, following the same rules as above (Supplementary Fig. 13A). Each trial comprised a single 20-ms blue pulse, which elicited 1 – 5 (mean: 2) spikes from the targeted cells. Each cell was stimulated 243 – 480 (mean: 326) times over a total of 50 min of recording. We recorded the responses from the targeted cells and all other cells. Motivated by the previously characterized neuroanatomy^20,21^, we tested for synaptic connections between cells separated by ≤ 400 μm.

Spontaneous membrane voltage fluctuations were much larger than a monosynaptic postsynaptic potential (PSP), so PSPs were not detectable in single trials. Hence, for each cell pair, we calculated the stimulus-triggered average subthreshold waveform, divided by the mean number of optically evoked presynaptic spikes (Fig. 3A, Methods). Our analysis corrected for off-target blue light effects (Supplementary Fig. 14; Methods). These protocols revealed clear inhibitory postsynaptic potentials (IPSPs; Fig. 3A). Median unitary IPSP amplitude among connected cells was -2.8% of a spike height.

**Figure 3.**
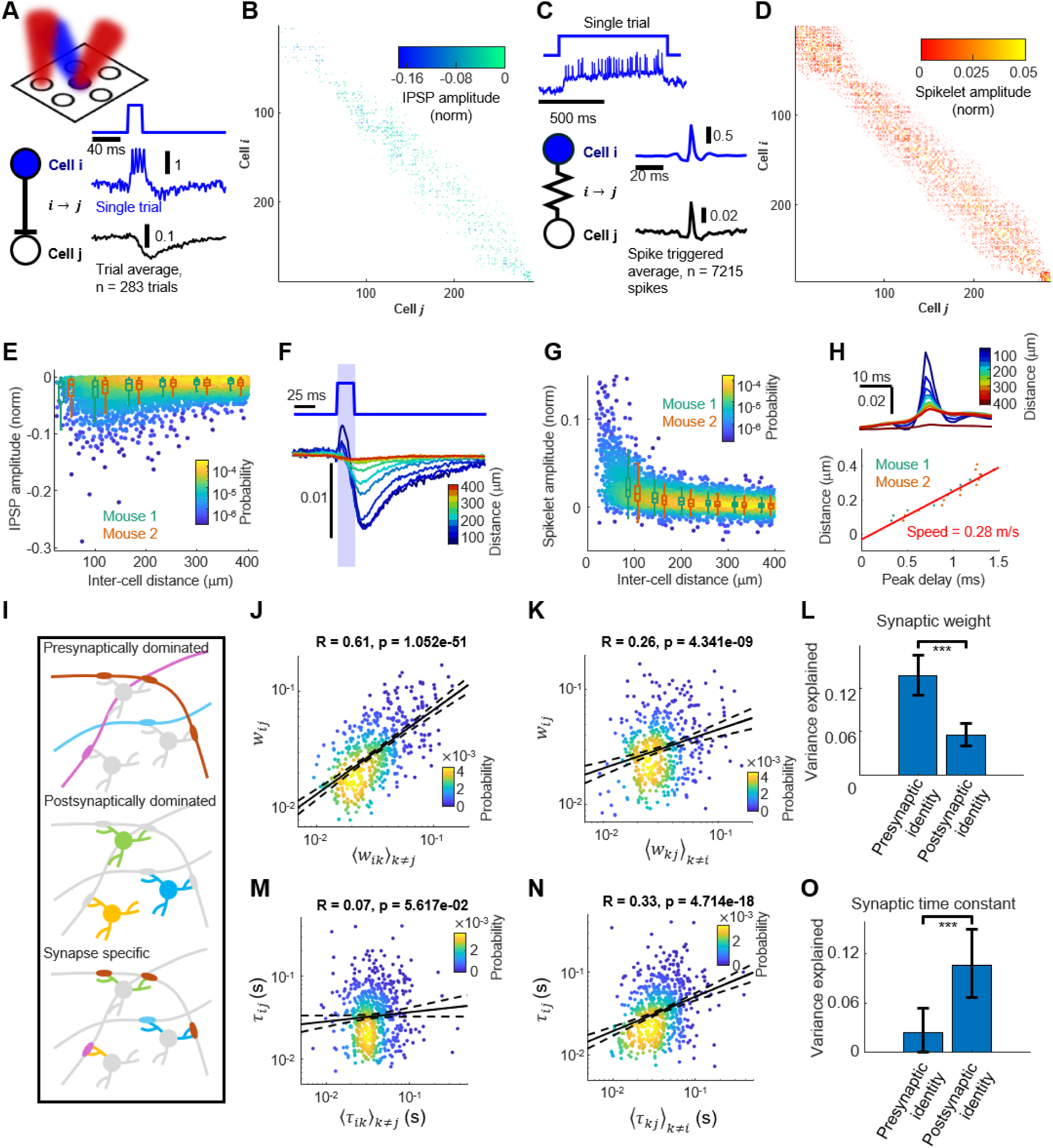
Mapping synaptic and gap junctional connectivity by all-optical electrophysiology *in vivo*. (A) Synaptic connectivity mapping. Top: targeted stimulation evoked spikes in presynaptic cell *i*. Bottom: trial-averaged response (n = 283 trials) in postsynaptic cell *j*. (B) Synaptic connectivity matrix for one mouse. The contrast of each matrix element is modulated by its p-value: 0.001 corresponds to maximum contrast, 0.5 corresponds to zero contrast. Inhibitory post-synaptic potential (IPSP) amplitude is measured in units of spike height per presynaptic spike. (C) Gap junction mapping. Cells were stimulated with 750 ms pulses in a pseudo-random sequence to evoke rapid spiking (blue). Spike-triggered average waveforms in a pre-junctional cell, *i*, and post-junctional cell, *j* (n = 7215 spikes of cell i). (D) Gap junctional connectivity matrix for one mouse. The contrast of each matrix element is modulated by its p-value: 0.001 corresponds to maximum contrast, 0.5 corresponds to zero contrast. Spikelet amplitude is measured in units of spike height. (E) Distance-dependent decay in IPSP amplitude. Box plots show median amplitude in 60-μm wide bins of cell-cell distance (box top and bottom: 25^th^ and 75^th^ percentile, whisker length: 1.5x inter-quartile range). Green: mouse 1, orange: mouse 2. (F) Distance-dependent population average IPSP waveform. Cell pairs were divided into 10 distance bins (bin size = 30 μm), and the average IPSP waveform was computed for each bin. Blue light scatter evoked a small stimulus artifact for closely spaced cell pairs (blue shading). (G) Distance-dependent decay in spikelet amplitude. Box plots show median amplitude in 60-μm wide bins of cell-cell distance, for two mice (box features same as in E). (H) Top: distance-dependent population average spikelet waveform. Cell pairs were divided into 10 distance bins (bin size = 30 μm), and the average spikelet waveform was computed for each bin. Bottom: spikelet peak, determined by sub-frame interpolation, shows distance-dependent time delay, consistent across two mice. Linear fit (red) yielded a propagation speed of 0.28 m/s. (I) Schematic illustration of synaptic properties determined by 1) presynaptic identity, 2) postsynaptic identity, or 3) synapse-specific features. (J) IPSP amplitude *w_ij_* for pair (*i*,*j*) is strongly correlated with mean output weight of cell *i* onto other cells 〈*w_ik_*〉*_k_*_≠*j*_. Solid line: log scale linear fit. Dotted line: 95% confidence interval. Log scale Pearson correlation, R = 0.61, p = 1.1E-51, n = 497 pairs. (K) IPSP amplitude *w_ij_* for pair (*i*,*j*) is weakly correlated with mean input weight of other synaptic inputs to cell *j*, 〈*w_ki_*〉*_k_*_≠*i*_. Log scale Pearson correlation, R = 0.26, p = 4.3E-9, n = 482 pairs. (L) More variance of IPSP amplitude is explained by presynaptic identity than by postsynaptic identity. We fitted the variance using a linear mixed-effects model, then ran bootstrap simulations to determine the consistency of the ordering of the pre- and postsynaptic variance components (see Methods). ***: < 0.001 of the simulations produced higher postsynaptic variance. (M) IPSP recovery time constant *τ_ij_* for pair (*i*,*j*) is uncorrelated with mean time constant of other cells postsynaptic to cell *i*, 〈*τ_ik_*〉*_k_*_≠*ij*_. Solid line: log scale linear fit. Dotted line: 95% confidence interval. Log scale Pearson correlation, R = 0.07, p = 0.06, n = 653 pairs. (N) IPSP time constant *τ_ij_* for pair (*i*,*j*) is correlated with mean time constant of other synaptic inputs onto cell *j*, 〈*τ_kj_*〉*_k_*_≠*i*_. Log scale Pearson correlation, R = 0.33, p = 4.7E-18, n = 651 pairs. (O) More variance of IPSP decay time constant is explained by postsynaptic identity than by presynaptic identity. We fitted the variance using a linear mixed-effects model, then ran bootstrap simulations to determine the consistency of the ordering of the pre- and postsynaptic variance components (see Methods). ***: < 0.001 of the simulations produced higher presynaptic variance.

In contrast, correlated fluctuations were ∼20% of a spike height, highlighting the importance of using trial-averaged evoked activity for synaptic connectivity mapping.

To determine statistical significance, we compared the measured IPSP for cell pair *i*-to-*j* to a null distribution calculated from the mean voltage waveform of cell *j* triggered on pulses targeting faraway cells (> 800 μm, Supplementary Fig. 15A,B; Methods). This procedure established a 50% detection probability for IPSP amplitude of -2 ± 1.1% of a spike (mean ± s.d., *n* = 100 neuron-pairs from each mouse, near detection cutoff).

We identified 802 statistically significant synaptic connections out of 7,280 pairs tested in one mouse, and 667 of 7,032 pairs tested in a second mouse. This corresponds to 3 ± 3 outgoing connections per cell and 3 ± 3 incoming connections per cell out of 25 ± 9 possible connections tested in each direction (mean ± s.d., *n* = 576 cells, 2 mice), i.e. for cells < 400 μm apart, the pair-wise probability of unidirectional connections above our detection threshold was 11 ± 9% (mean ± s.d.).

#### Gap junction coupling

L1 interneurons also have direct electrical coupling via gap junctions^22,45,46^, so a spike in one cell can drive a spikelet of a few millivolts in a connected cell. To identify these connections, we measured spike-triggered average waveforms of putative post-junctional cells, triggered off spikes of putative pre-junctional cells (cross-STA). To evoke temporally dense but pair-wise uncorrelated spike patterns, we stimulated the neurons with pseudo-random binary sequences derived from a Hadamard-Walsh code (binary 1: 750 ms blue on; binary 0: 750 ms blue off; Supplementary Fig. 13B). Thus, every cell was stimulated with a unique pattern, with 50% duty-cycle. Stimulated cells spiked rapidly (25.4 ± 16.6 Hz, mean ± s.d.) compared to non-stimulated cells (3.9 ± 4.5 Hz), and evoked spikes were largely uncorrelated between cells. Each cell received on average 640 stimuli, which elicited on average a cumulative 12,380 spikes per cell.

We then calculated cross-STA waveforms for all cell pairs separated by ≤ 400 μm. To minimize noise from spikes in post-junctional cells we (a) restricted the averages to blocks where the pre-junctional cell was stimulated and the putative post-junctional cell was not and (b) replaced residual post-junctional spikes with linearly interpolated subthreshold voltages. Spikelets were clearly visible for a subset of the cell pairs (Fig. 3C). Median spikelet amplitude among connected cells was 1.2% of a spike.

To determine the statistical significance, we compared the measured spikelet for cell pair *i*-to-*j* to a null distribution calculated from the mean voltage waveform of cell *j* triggered by spikes of faraway cells (> 800 μm, Supplementary Fig. 15C; Methods). This procedure established a 50% detection probability for spikelet amplitudes of 0.8 ± 0.48% of a spike (mean ± s.d., *n* = 100 neuron-pairs from each mouse, near detection cutoff). We identified 3,553 statistically significant directed functional electrical connections out of 7,280 pairs tested in one mouse, and 2,199 of 7,032 in a second mouse. This corresponds to 10 ± 6 outgoing connections per cell and 10 ± 5 incoming connections per cell out of 25 ± 9 possible connections tested in each direction (mean ± s.d., *n* = 576 cells, 2 mice), i.e. for cells < 400 μm apart, the pair-wise probability of unidirectional gap junction connections above our detection threshold was 41 ± 15%.

Since the spikelets were approximately synchronous with the pre-junctional spikes, we had to control for two possible sources of crosstalk. First, scattered fluorescence from spikes in the pre-junctional cell could contaminate the post-junctional measurement. We calculated the STA fluorescence of interstitial pixels (i.e. pixels not associated with a cell) as a function of distance from the triggering cell. This scattered fluorescence signal decayed over a substantially shorter distance than the spikelet amplitude (at 60 μm separation, spikelet:scattered fluorescence > 2.5). We excluded close (< 60 μm) neuron pairs from further analysis due to possible contamination from scattered fluorescence (Supplementary Fig. 17; Methods).

Second, fluorescence from *en passant* neurites of the pre-junctional cell could contaminate the post-junctional measurement. The Voltron1 reporter was nominally soma-localized, but this localization was not perfect. To check for this form of crosstalk, we repeated the gap junction mapping using a higher magnification (7.5x system magnification) objective (Supplementary Fig. 18A,B). We calculated the spatial STA footprints of putative post-junctional cells, triggered either off post-junctional spikes (self-STA) or off pre-junctional spikes (cross-STA; Supplementary Fig. 18C). For genuine gap junction connections, both STA footprints would show the shape of the post-junctional cell; for *en passant* neurites, the cross-STA would show the (possibly out of focus) neurite. Spikelet amplitude was strongly correlated with the coefficient of the self-STA footprint fitted to the cross-STA image (R = 0.94, p < 1E-15, *n* = 720 pairs from 48 cells; Supplementary Fig. 18D), supporting a post-junctional somatic origin for the detected signals.

#### The structure of connectivity in L1

We then examined the structures of the connectivity matrices (Fig. 3B,D). The synaptic connectivity matrix had a single-scale network structure^47^, with an exponential degree distribution. The probability of connection was 24% for closely separated cells (< 150 μm) and fell with a ∼105 μm length constant (Supplementary Fig. 19C,D). The gap junction matrix showed greater connectivity (connection probability 83% for cells separated by < 150 μm) and fell with a 159 μm length constant (Supplementary Fig. 19E,F).

IPSP and spikelet amplitudes both decayed with cell-cell distance (Fig. 3E,G), and the population-average IPSP and spikelet waveforms had similar time courses across distances (Fig. 3F,H). The spikelet peak times, determined by sub-frame interpolation, showed a distance-dependent time delay corresponding to a conduction speed of 0.28 m/s, consistent across *n* = 2 mice (Fig. 3H). This speed is comparable to prior reports of action potential propagation in unmyelinated neurites^48–51^, and further rules out optical crosstalk (which would be instantaneous) as a dominant contribution to the spikelets. IPSP and spikelet amplitudes both followed approximately log-normal distributions (Supplementary Fig. 19A,B). Prior anatomical, tracing, and patch clamp measurements had also found log-normal distributions of synaptic strength^52–55^, but there is little prior characterization of the distribution of gap junctional strengths^56,57^.

The population-average combined effect of gap junctional and synaptic connectivity is that a single L1 action potential evoked a brief excitatory pulse with a duration of ∼2 ms that rapidly propagated outward (0.28 m/s) with a ∼160 μm decay length constant, followed by a slower field of inhibition that peaked around 30 ms and decayed with a time constant of ∼35 ms (Movie 3).

Because our data paired many post-synaptic cells with each presynaptic cell, and *vice versa*, we could compare the contributions of presynaptic and postsynaptic identity to synaptic properties, i.e.: to what extent are synaptic properties (IPSP amplitude, decay time constant) influenced by the presynaptic cell vs. the postsynaptic cell (Fig. 3I)? For each synaptic connection from cell *i* to *j*, we plotted connection strength, *w_ij_*, vs. the mean strength of all connections of cell *i* to cells *k*, 〈*w_ik_*〉*_k_*_≠*i*_ (Fig. 3J). We then made a similar plot holding constant the postsynaptic identity *j*, and plotting the incoming *i* to *j* synaptic strength, *w_ij_*, vs. the mean strength of incoming connections from other cells *k*, 〈*w_kj_*〉*_k_*_≠*i*_ (Fig. 3K). The per-cell mean outgoing connection strength was a better predictor of single-synapse strength than was mean incoming connection strength (outgoing: R = 0.61, p = 1.1E-51, n = 497 pairs, vs. incoming: R = 0.26, p = 4.3E-9, n = 482 pairs).

Surprisingly, applying the same analysis to synaptic recovery time-constant, *τ_ij_* , yielded the opposite result (Fig. 3M,N). The per-cell mean inbound synaptic time constant, 〈*τ_kj_*〉*_k_*_≠*i*_, was a better predictor of single-synapse time constant *τ_ij_* than was mean outgoing time constant, 〈*τ_ik_*〉*_k_*_≠*i*_ (incoming: R = 0.33, p = 4.7E-18, n = 651 pairs, vs. outgoing: R = 0.07, p = 0.06 , n = 653 pairs). A linear mixed-effect model reproduced these findings (Fig. 3L,O; Methods). While the mechanistic basis for these observations is unknown, we speculate that the amplitude effect might reflect cell-to-cell differences in expression of presynaptic release machinery. The time-constant effect could reflect, e.g. cell-to-cell differences in the ratio of GABA_A_ to GABA_B_ receptor expression. The gap junction connection strength showed little association with pre- or post-junctional identity in this analysis.

### Spontaneous spikes and subthreshold voltages have distinct spatiotemporal dynamics

We next turned to the large-scale spontaneous dynamics of the L1 population, as the mouse performed self-directed locomotion on a treadmill. We used whisking as an indicator of arousal^58^, and compared L1 dynamics in whisking and non-whisking (“quiet”) states. We decomposed each cell’s voltage signal into subthreshold and spiking components (Methods) to assess input/output properties of the circuit (Fig. 4A). To avoid locomotion-induced motion artifacts, we excluded running epochs from these analyses.

**Figure 4.**
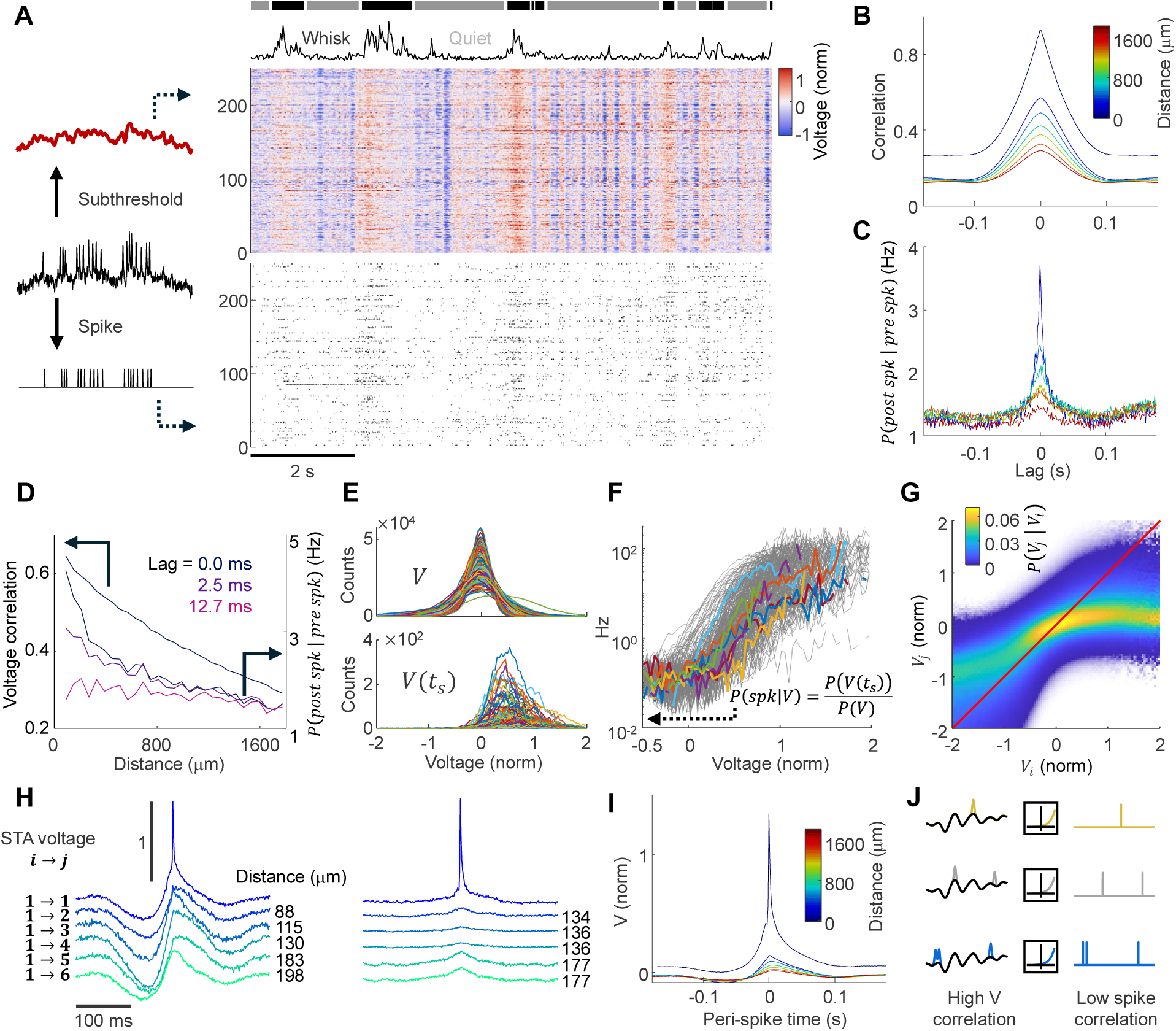
Threshold nonlinearities and private excursions block correlation transfer from subthreshold to spikes. (A) Fluorescence recordings were split into subthreshold and spiking traces. Subthreshold voltages (top) showed population-wide correlations, whereas spiking (bottom) was less correlated. (B) Mean subthreshold cross-correlation functions resolved by pairwise distance (quiet state). Each plot corresponds to a 60 μm-wide bin. Cell pairs in the 0 – 60 μm bin were omitted due to fluorescence scatter. (C) Mean spike-rate cross-correlograms, resolved by pairwise distance (quiet state). Bins are as in (B). (D) Distance-dependent decay of subthreshold equal-time cross-correlation (left axis) and spike-rate cross correlograms for three lags (0 ms, 2.5 ms, and 12.7 ms; right axis). (E) Histograms of subthreshold voltage at (top) all times and (bottom) 10 ms interval prior to spikes. (F) Voltage-dependent conditional spike rate. Seven randomly selected cells are shown in color. (G) Probability distributions of subthreshold voltage of cell *j* conditioned on the subthreshold voltage of cell *i* for all (*i*,*j*) pairs < 200 μm apart. *n* = 505 cells, *N* = 2 mice. Red line has slope of 1. (H) Example spike triggered average waveforms for individual cell pairs. Each column shows a particular triggering cell and self- and cross-STA waveforms for individual triggered cells. (I) Mean STA waveforms, resolved by pairwise distance (quiet-state). Bins are as in (B). (J) Schematic illustrating how threshold nonlinearity combined with private positive-going voltage excursions blocks correlation transfer from subthreshold to spiking.

In the quiet state, the voltage dynamics were strongly correlated between nearby cells (correlation = 0.6, 60 μm < r < 120 μm) and decayed with an exponential length constant of 2.4 mm (Fig. 4B; Supplementary Fig. 20D); in the whisking state, the short-range correlation was 0.5 and the decay length constant was 4.0 mm (Supplementary Fig. 20A,D). Overall, the subthreshold dynamics were relatively low dimensional: principal components analysis (PCA) showed that the first 10 components accounted for 72% of the variance in the quiet state and 62% of the variance in the whisking state (Supplementary Fig. 21B).

In the quiet state – but not the whisking state – the subthreshold dynamics often showed global oscillations between 1 and 10 Hz (Supplementary Fig. 21A). These oscillations corresponded to large-scale wave propagation (Movie 4). We tracked the wavefronts (Methods) and observed a variety of propagation directions and speeds (Supplementary Fig. 21D-H), with a slight bias toward the medial-posterior direction and an average speed of 18.2 ± 8.3 mm/s (mean ± s.d., n = 3858 events, 2 mice). For a typical 5 Hz oscillation, the wavelength was ∼4 mm (Supplementary Fig. 21H). Similar state-dependent oscillations have been reported by *in vivo* patch clamp studies in cortex^58–62^, and similar traveling wave dynamics have also been reported via local field potential and meso-scale GEVI and Ca^2+^ recordings^63–65^.

The spiking showed a markedly different correlational structure (Fig. 4C). While the subthreshold cross-correlations decayed slowly over space (quiet-state length constant 2.4 mm) and time (quiet-state full-width at half-maximum 80 ms), the spike-rate cross-correlogram had a peak that was sharply localized in space (length constants of 277 and 270 μm for quiet and whisking state respectively) and time (decayed by 50% at lags of just 2.5 ms; Fig. 4D, Supplementary Fig. 20F,G). The length-scale of this decrease in spike synchrony was within a factor of two of the length-scale of the pairwise gap-junction connections (159 μm, Supplementary Fig. 19F).

### Threshold nonlinearities and private excursions limit the transfer of shared voltage fluctuations to spikes

Why were the cell-to-cell correlations in spike timing so different from the correlations in subthreshold dynamics? We sought to assess the relative contributions of private variations in input-output properties vs. private subthreshold fluctuations.

First, for each cell, we determined an empirical voltage-to-spike relationship from the spontaneous activity data. We calculated the distribution of all subthreshold voltages during spontaneous activity; and the distribution of subthreshold voltages in the 10 ms preceding spike times (Methods, Fig. 4E). The ratio of these two distributions yielded an estimate of the conditional spike rate as a function of subthreshold voltage, i.e. the voltage-to-spike relationship. To avoid arousal-related effects, we only used quiet state epochs for this analysis. These curves all had a characteristic sigmoidal shape, with substantial cell-to-cell variations in threshold and maximum firing rate (Fig. 4F).

Next, we searched for private subthreshold fluctuations related to spiking. We examined the joint distribution of the voltage of each cell (V_i)_ and the voltage of its neighbors (V_j;_ < 200 μm distance). The correlation between V_i_ and V_j_ was strongest near resting potential and for hyperpolarizing fluctuations, but was markedly reduced as the subthreshold voltage approached spike threshold (Fig. 4G). This analysis pointed to private positive-going fluctuations preceding spikes.

We then examined the population-level voltage dynamics specifically around spike times. We calculated the self- and cross-STAs of the membrane potential. For cells separated by < 200 μm, the subthreshold components of these two waveforms closely followed each other at lags > 30 ms (Fig. 4H). However, the spiking cell saw both a higher baseline and an upward divergence in the self-STA in the 30 ms preceding the spike (Fig. 4I), indicating shared slow subthreshold fluctuations and private fast spike-inducing fluctuations.

Together, these results showed that the rare positive-going fluctuations in subthreshold voltage were largely private to individual cells, whereas the predominant fluctuations around and below resting potential were more strongly correlated. That is, the correlated component of subthreshold voltage did not strongly drive spiking, and the voltages that drove spiking were only weakly correlated. The nonlinear and heterogeneous voltage-to-spike relationship preferentially weighted the depolarizing excursions, helping explain the different spatial and temporal structures of subthreshold correlations and spike synchrony (Fig. 4J).

### Intrinsic properties shape subthreshold-to-spiking transfer functions during spontaneous activity

Given the diverse single-cell intrinsic properties (Fig. 2), we asked how these intrinsic properties affected the spontaneous circuit dynamics. We first compared the subthreshold-to-spiking transformations under optogenetic drive vs. spontaneous activity. We divided cells into five bins of increasing optical rheobase (a measure of intrinsic excitability), measured during ramp optogenetic stimulation. For each bin we calculated the mean conditional firing rate as a function of subthreshold voltage, during either ramp optogenetic stimulation or quiet state spontaneous activity (Fig. 5A). This procedure produced similar curves for the evoked and spontaneous activity: in both datasets, cells with higher optical rheobase showed lower firing rate for a given subthreshold depolarization (Fig. 5A; Supplementary Fig. 22A,B). Thus, the optical rheobase measured a key aspect of the subthreshold-to-spiking transfer function, applicable to spontaneous activity without optogenetic drive.

**Figure 5.**
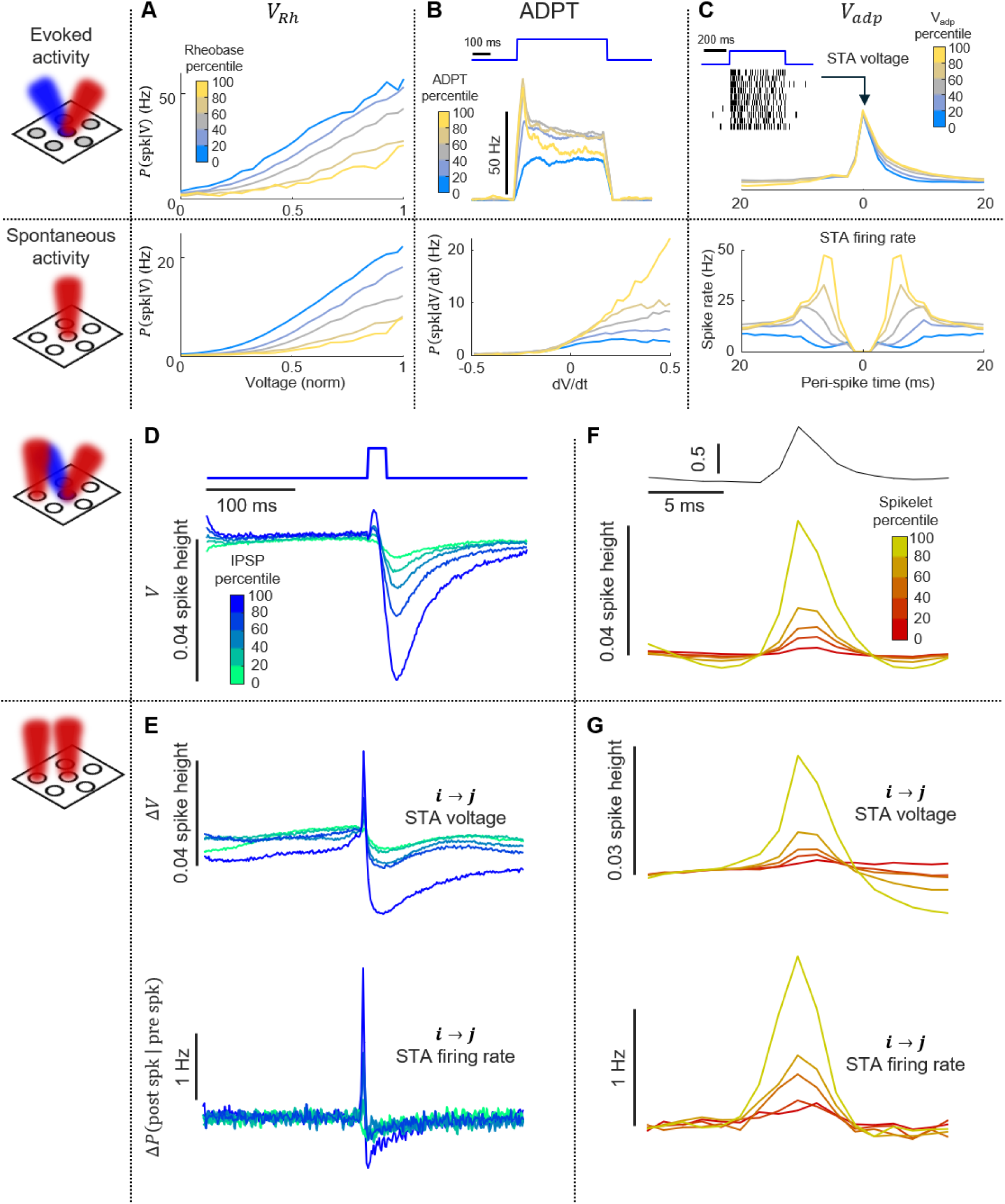
Intrinsic properties and local connectivity shape network dynamics. (A) Optical rheobase affects excitability during spontaneous activity. Top: spike rates conditioned on subthreshold voltage, *V*, during optogenetic ramp stimulus, sorted by quintiles of optical rheobase, *V_R_*_ℎ_. Bottom: spike rates conditioned on *V* during spontaneous activity, sorted by quintiles of *V_R_*_ℎ_. (B) Spike-rate adaptation affects sensitivity to voltage upstroke velocity during spontaneous activity. Top: mean spike rate during optogenetic step stimulation, sorted by quintiles of spike-rate adaptation (*ADPT*). Bottom: spike rates conditioned on *dV*/*dt* during spontaneous activity, sorted by quintiles of *ADPT*. (C) Spike after-depolarization affects bursting during spontaneous activity. Top: mean STA waveform, sorted by quintiles of spike afterdepolarization, *V_adp_*. Bottom: Spike autocorrelograms during spontaneous activity, sorted by quintiles of *V_adp_*. n = 280 cells in (A-C). (D) Optically evoked IPSP waveforms, sorted into quintiles of IPSP strength. n = 800 pairs for (D) and (E). (E) Synaptic inhibition affects network correlations during spontaneous activity. Top: Cross-STA voltage waveforms, sorted by quintiles of synaptic strength. Bottom: cross-STA firing rates, sorted by quintiles of synaptic strength. Waveforms and spike rates are measured relative to distance-matched unconnected controls (Methods). (F) Optically evoked spikelet waveforms, sorted into quintiles of spikelet amplitude. Top: Spike waveform. Bottom: evoked spikelets. n = 3553 pairs for (F) and (G). (G) Gap junction coupling induces network synchrony. Top: cross-STA voltage waveforms, sorted by quintiles of optogenetically evoked spikelet amplitude. Bottom: cross-STA firing rates, sorted by quintiles of spikelet amplitude. Waveforms and spike rates are measured relative to distance-matched unconnected controls (Methods).

Under step-wise optogenetic stimulation, cells showed variable amounts of spike-rate adaptation (Fig. 2A), a feature not captured by the simple excitability curves calculated in Fig. 5A. Spike-rate adaptation captures the degree to which the cell acts as a high-pass filter, converting a step input into a transient output. We thus hypothesized that spike-rate adaptation might correlate with response to the rate of upswing of the subthreshold voltage, dV/dt. We divided the cells into five bins of spike-rate adaptation under step-wise optogenetic stimulation, and for each bin we calculated the mean conditional firing rate as a function of dV/dt during quiet-state spontaneous activity. Cells with higher spike-rate adaptation preferentially spiked in response to rapidly rising inputs (Fig. 5B; Supplementary Fig. 22C). Thus, spike-rate adaptation adds a temporal high-pass filter to the subthreshold-to-spiking transfer function during spontaneous activity.

The spike afterpotential influences how quickly the membrane potential returns to baseline after a spike. We reasoned that spikes with greater after-depolarization would be closer to threshold and hence more likely to fire a second spike in rapid succession after a first. We sorted the cells into five bins by afterdepolarization of the optogenetically evoked spikes. For each bin we then calculated the mean spike autocorrelogram during spontaneous activity (quiet and whisking). Cells with greater after-depolarization were more likely to fire spikes in bursts (Fig. 5C; Supplementary Fig. 22D). Together, these results show how single-cell excitability properties can set the sensitivity of spiking to subthreshold voltage, rate of change of subthreshold voltage, and recent spike history, during spontaneous behavior.

### Local connectivity shapes network dynamics

We then asked how the synaptic and gap junctional connectivity affected spontaneous network dynamics. We had already observed that the individual pairwise cross-STA waveforms failed to reveal synaptic connections, due to dominant contributions from shared subthreshold fluctuations (Supplementary Fig. 11). We thus sorted neuron-pairs into quintiles of optogenetically determined synaptic strength, and for each quintile we constructed a control population of nearby pairs without a detected connection (Supplementary Fig. 23A; Methods). The influence of synaptic inhibition became apparent in the difference between sub-group averages of the connected vs. non-connected control cross-STA waveforms (Fig. 5D,E; Supplementary Fig. 23B).

For cell-pairs in the top quintile of synaptic strength, a presynaptic spike induced a median postsynaptic voltage decrease of -2.1% of a spike height in the 15 – 30 ms after a spike, relative to the distance-matched unconnected controls (p = 1.9E-33, n = 293 cell pairs, 2 mice). For the median synaptic strength (among connected pairs), this hyperpolarization was -1% of a spike height (p = 5.7E-39, n = 1,467 cell pairs, 2 mice, Wilcoxon rank sum test; Supplementary Fig. 23F). For the control population of pairs without a detected connection, we calculated the cross-STA between arbitrarily designated “pre-synaptic” and “post-synaptic” cells. The STA waveform showed an increase by 1% of a spike height in the interval 15 – 30 ms after the triggering spike vs. the -20 to - 5 ms before the spike, reflecting correlated subthreshold fluctuations (n = 14,433 pairs, 2 mice).

To see how pairwise inhibition affected network output, we then applied the same procedure to the spike-rate cross-correlograms. The raw pair-wise cross-correlograms did not show a clear signature of synaptic inhibition (Fig. 4C), but comparing the mean cross-correlograms for connected vs. nearby non-connected control subpopulations revealed clear spike-triggered inhibition in spike rate (Fig. 5E, Supplementary Fig. 23D,H). For cell-pairs in the top quintile of synaptic strength, a presynaptic spike induced a median postsynaptic decrease in spike rate of 0.18 Hz in the 5-13 ms after the triggering spike, measured relative to nearby non-connected controls. Integrating the decrease in spike rate over the entire IPSP duration gave a median loss of 7×10^-3^ spikes in the post-synaptic cell. For cell-pairs near median synaptic strength (among connected pairs), this decrease was 0.083 Hz, corresponding to a loss of 2.7×10^-3^ spikes.

Finally, we explored the influence of gap junctional connections on spontaneous dynamics, using the same analysis procedures as for synaptic connections. Sorting cells by quintiles of gap junction strength and comparing to nearby non-connected controls revealed gap-junction-mediated subthreshold spikelets (Fig. 5F,G, Supplementary Fig. 23C,E,G). These spikelets enhanced spike synchrony: for cell-pairs in the top quintile of gap junction coupling, the rate of synchronous firing (time lag ≤ 2.5 ms) was 0.86 Hz greater than near-synchronous firing (time lag 5 ms, p = 8.6E-53, n = 1151 pairs, 2 mice, Wilcoxon rank sum test against unconnected neighbors; Supplementary Fig. 23I). Integrating the change in spike rate across the entire spikelet duration gave a gain of 1.7×10^-3^ spikes. For the median gap-junctional coupling, the synchrony enhancement was 0.32 Hz (p = 1.9E-61, n = 5752 pairs, 2 mice, Wilcoxon rank sum test against unconnected neighbors), corresponding to a gain of 7×10^-4^ spikes. The overall rate of spike synchrony showed a distance-dependent decay, with a length constant of ∼270 μm (Fig. 4D, Supplementary Fig. 20F). Thus, stronger optically measured electrical coupling was associated with greater pairwise spike synchrony during spontaneous activity.

We noted that the number of spikes lost per connected pair due to inhibition (median 2.7×10^-3^ spikes) was approximately four-fold greater than the number of spikes gained due to gap junction coupling (7×10^-4^). To determine the overall contributions of inhibitory and gap junctional (excitatory) interactions to the circuit dynamics, these numbers should be scaled by the connection probabilities, which we estimated to be 11 ± 9% for synaptic inhibition and 41 ± 15% for gap junctions (Fig. 3B,D). Hence, oppositely acting synaptic and gap-junctional influences are approximately balanced – despite different timescales, length scales, and statistical connection structures. These estimates are necessarily approximate, on account of the incomplete sampling of the network and approximate measurements of pairwise connection strengths.

### Brain state differentially modulates functional and genetic L1 interneuron subtypes

L1 NDNF interneurons have two genetic subtypes, demarcated by expression of neuropeptide Y (NPY). We produced a double transgenic mouse line, NDNF-cre::NPY-GFP, which labeled the NPY^(+)^ subtype with GFP (Fig. 6A). We expressed virally delivered Optopatch broadly in the NDNF population and recorded neural activity during self-directed locomotion and whisking. We then imaged the GFP channel and determined the NPY status of each recorded neuron. Approximately 49% of recorded neurons were NPY^(+)^ (166/320 in mouse 1, 145/320 in mouse 2). The two subtypes were intermixed throughout the field of view, and had similar spike SNR and similar average spike-rate (Supplementary Fig. 24A-D).

**Figure 6.**
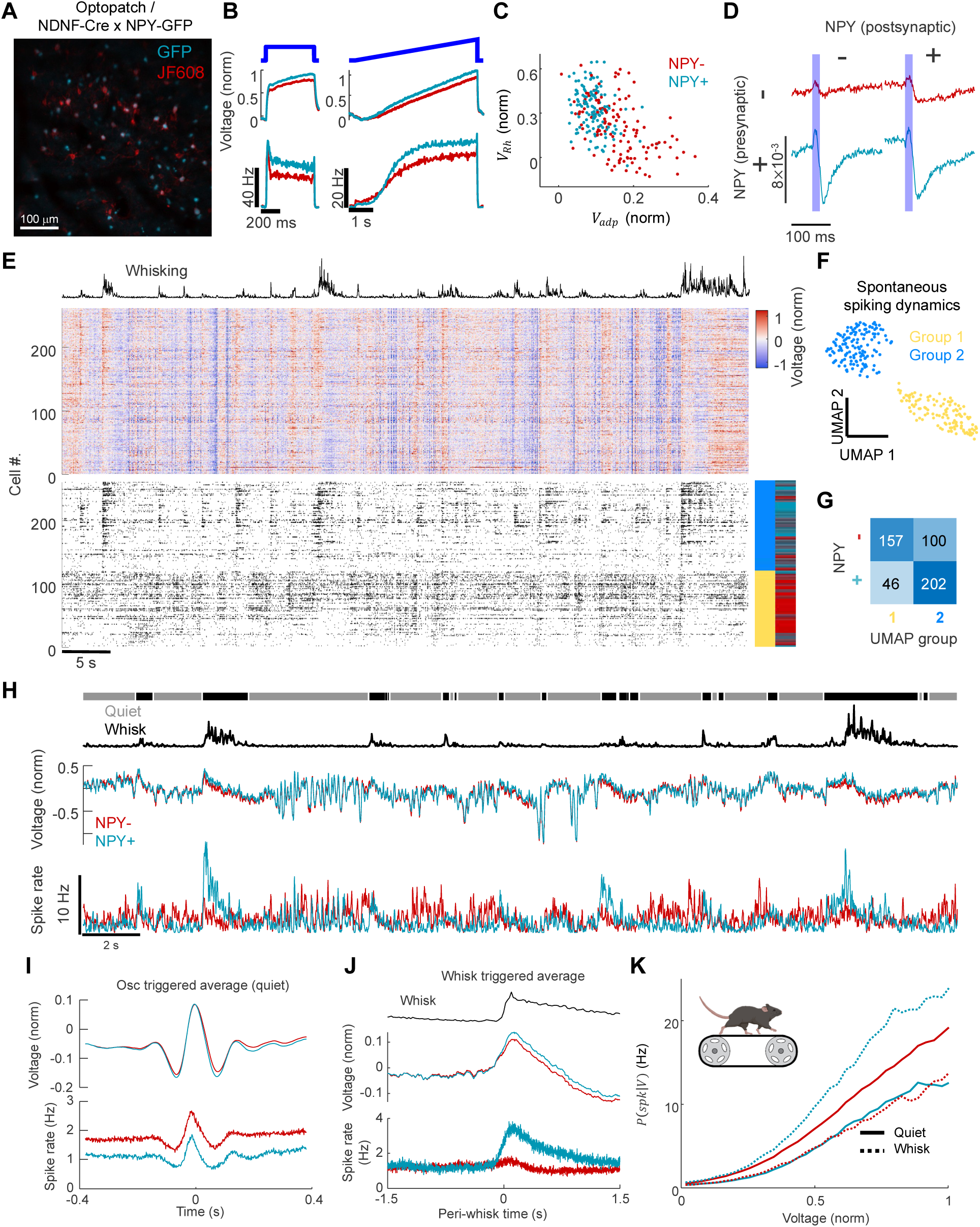
Brain state differentially modulates intrinsic excitability of genetic subtypes in L1. (A) Spinning disk confocal image of Optopatch construct expressed in the barrel cortex L1 of NDNF-Cre x NPY-GFP mouse. Red: JF608, cyan: GFP. (B) Genetic subtype-average responses to targeted optogenetic stimulation. Left: step stimulation. Right: ramp stimulation. Top: subthreshold voltage. Bottom: firing rate. Red: NPY^(-)^ (n = 125 cells), cyan: NPY^(+)^ (n = 132 cells). (C) NPY^(+)^ and NPY^(-)^ cells have significantly different intrinsic properties. V_Rh_ and V_ADP_ are shown here. Red: NPY^(-)^, cyan: NPY^(+)^. (D) Genetic subtype-dependent mean optically evoked IPSP waveforms. n = 1420 (−) → (−), 1818 (−) ↔ (+), 2224 (+) → (+) pairs. Blue shading marks the stimulation pulse. Red: presynaptic NPY^(-)^, cyan: presynaptic NPY^(+)^. (E) Subthreshold voltage (middle) and spiking dynamics (bottom) during spontaneous behavior. The cells were sorted by UMAP on the spiking alone, without input from NPY marker (Methods). The two functional subgroups (blue/yellow bar), co-assorted with the NPY indicator (red/cyan bar). The same sorting was applied to the subthreshold voltages. (F) 2D UMAP embedding of spontaneous spiking dynamics, clustered into two groups. Yellow: group 1, blue: group 2. (G) Confusion matrix comparing genetic identity and functional subgroup identity. N = 2 mice. (H) Genetic subtypes have similar population average subthreshold voltages but distinct behavior-dependent spiking patterns. Top: behavior-state classification. Middle: subtype-average subthreshold voltage. Bottom: subtype-average spike rate. n = 141 NPY^(-)^ cells, 117 NPY^(+)^ cells. Red: NPY^(-)^ average, cyan: NPY^(+)^ average. (I) Peak-triggered average of subthreshold oscillations (quiet state). Top: subthreshold voltages. Bottom: spike rates. n = 5888 events. Red: NPY^(-)^ average, cyan: NPY^(+)^ average. (J) Whisking onset-triggered average subthreshold voltage (top) and spike rate (bottom). n = 547 events. Red: NPY^(-)^, cyan: NPY^(+)^. (K) Brain state-dependent subtype-average excitability curves during spontaneous activity. subthreshold voltage-dependent conditional spike rate during quiet epochs (solid) and int the 640-ms window following the whisking onset (dashed). Red: NPY^(-)^, cyan: NPY^(+)^.

We then compared the intrinsic properties and connectivity of the two subtypes. The NPY^(–)^ neurons had greater after-depolarization, lower optical rheobase, more spike-rate adaptation, slightly lower maximum firing rate, and a shorter membrane time constant (Fig. 6B,C; Supplementary Fig. 24E). Using these five electrophysiological parameters, a logistic-regression classifier identified cell type with 78% (95% CI [74%, 81%]) cross-validated balanced accuracy, compared with a 50% (95% CI [46%, 55%]) chance level. The most impactful feature for cell-type classification was spike afterdepolarization.

The synaptic connectivity also differed among the populations (Fig. 6D): NPY^(+)^ cells were significantly more likely to synapse onto other L1 interneurons of either class (outgoing connection probability within 400 μm: 14 ± 15%, n = 307 NPY^(+)^ cells, 2 mice) compared to NPY^(–)^ cells (connection probability: 7 ± 11%, n = 269 NPY^(-)^ cells, 2 mice, *P* < 0.001, Wilcoxon rank sum test; Supplementary Fig. 25A). The IPSP amplitudes averaged across all detected synaptic connections were significantly larger for NPY^(+)^ presynaptic cells (−3.7 ± 2.7% of a spike height, n = 977 pairs, 2 mice) than for NPY^(-)^ presynaptic cells (−3.1 ± 2.3% of a spike height, n = 490 pairs, *P* < 0.001, Wilcoxon rank sum test; Supplementary Fig. 25B). For synaptic inputs, IPSP durations were significantly longer in NPY^(+)^ postsynaptic cells (58 ± 63 ms, n = 699 pairs, 2 mice) than in NPY^(-)^ cells (43 ± 54 ms, n = 646 pairs, 2 mice, *P* < 0.001. Wilcoxon rank sum test; Supplementary Fig. 25C), consistent with prior patch clamp studies.^21^ The gap junction out-going and incoming connectivity did not differ systematically between NPY^(+)^ and NPY^(-)^ sub-types (Supplementary Fig. 25D,E).

We next compared the spontaneous dynamics of the two genetic sub-types. The population-average subthreshold voltages were almost indistinguishable between NPY^(+)^ and NPY^(-)^ subpopulations (*R* = 0.97, slope = 0.96; Fig. 6E,H), indicating that these populations largely sampled the same synaptic input field. Remarkably, the spiking patterns were strikingly distinct. We applied UMAP^39^ to sort the single-cell firing patterns during spontaneous activity (Methods). The sorted spike raster clearly showed two subpopulations (Group 1 and 2, Fig. 6E,F), which co-assorted with the NPY^(-)^ and NPY^(+)^ subtypes respectively (157/203 Group 1 cells were NPY^(-)^ and 202/302 Group 2 cells were NPY^(+)^, n = 2 mice; Fig. 6G). Thus, the NPY^(+)^ and NPY^(-)^ populations convert similar input information into different spiking outputs. Specifically, spiking dynamics at the onset of whisking were dramatically different: In NPY^(+)^ cells, spike rate increased from 1.4 ± 1.7 Hz to 2.8 ± 4.3 Hz (n = 248 cells, N = 2 mice, *P* < 0.001, Wilcoxon rank sum test). In NPY^(-)^ cells, spike rate did not show a significant change (from 1.8 ± 2.1 Hz to 2.0 ± 2.6 Hz, n = 257 cells, N = 2 mice; Fig. 6H,J; Supplementary Fig. 26 A,B).

How could near-indistinguishable subthreshold voltage patterns give rise to such distinct subpopulation-specific spiking patterns? We calculated subtype-average conditional firing rates as a function of subthreshold voltage, separately for quiet and whisking epochs. Whisking induced a significant decrease in NPY^(-)^ excitability and a significant increase in NPY^(+)^ excitability (Fig. 6K; Supplementary Fig. 26C). Thus, behavioral state selectively enhanced the response of the NPY^(+)^ neurons and muted the response of the NPY^(-)^ neurons to the same input field. We speculate that these different responses to whisking might reflect differing effects of arousal-related neuromodulation, though our data do not identify the mechanism. This hypothesis is consistent with a recent report that L1 NDNF interneuron subpopulations show differential modulation during sleep.^66^

## Discussion

We used simultaneous perturbation and recording of membrane voltage at population scale to map the excitability and connectivity of an interneuron network *in vivo*, and to follow the transformation from subthreshold input to spiking output in the same cells. This process relied on technical advances in four areas: (1) Identification of spectrally orthogonal and high-performance voltage actuator-indicator pairs; (2) development of a wide-area, high-speed, and high-sensitivity LED-based dual-wavelength structured illumination microscope; (3) creation of experimental protocols that separately probed intrinsic excitability, synaptic inhibition, and gap junction couplings; and (4) creation of data analysis algorithms that related these parameters to the subthreshold-to-spiking transformation during spontaneous activity. These tools can be used individually or in combination to dissect optically accessible neural circuits.

Earlier all-optical studies established the feasibility of measuring excitability and functional coupling *in vivo*^11–14,67^, but only in small ensembles of cells, and without resolving individual pair-wise synaptic potentials. Here, the scale, duration and linked multi-modal nature of the measurements allowed us to ask how time-dependent single-cell excitability properties, the strengths and kinetics of pairwise interactions, and behavioral state jointly sculpt ongoing circuit activity.

One benefit of measuring evoked and spontaneous responses in the same cells is that electrophysiological diversity can be connected directly to features of ongoing activity. Neuronal excitability varies within and between genetically defined populations^37,68^, and also within individual cells over time. In L1, optical rheobase predicted the voltage dependence of spontaneous firing, spike-rate adaptation predicted sensitivity to the rate of depolarization, and spike after-depolarization predicted short-lag bursting. These relationships show how the measured electrophysiological diversity affects circuit function.

The connectivity measurements revealed previously inaccessible elements of network organization. By sampling multiple outgoing and incoming connections for each neuron, we observed distinct contributions of the presynaptic vs. postsynaptic cell to synaptic properties. The presynaptic cell more strongly influenced IPSP amplitude, whereas the postsynaptic cell more strongly set IPSP decay time. Both parameters are important in governing cell-to-cell influences in the circuit. These effects may be due to differences in release machinery, receptor composition, or electrotonic filtering, but the present measurements do not establish molecular mechanisms.

Mapping synaptic and gap junctional interactions together also revealed their distinct spatiotemporal organization. Optically evoked spikelets were brief and broadly shared in the circuit, whereas inhibition developed more slowly, persisted for tens of milliseconds, and was sparser. The corresponding signatures in spontaneous activity were rapid spike synchrony and delayed spike suppression, respectively. Earlier studies established synchronizing effects of electrical coupling in cortical interneuron networks^69^, and prior all-optical work implicated L1 lateral inhibition in temporal filtering^13^. Our measurements show how these mechanisms combine *in vivo*, leading to short-time excitatory coupling followed by slower inhibition over similar spatial ranges. This organization suggests a network-level preference for brief, coincident activity over sustained firing.

An independently measured connection map was essential for relating synaptic and gap junctional interactions to spontaneous dynamics. Shared voltage fluctuations were ∼10-fold larger than pairwise couplings, so individual connections were generally not evident from the raw pairwise spike cross-correlograms or spike-triggered average voltages. Grouping pairs by optically measured coupling strength and comparing with nearby control pairs without a detected connection exposed the expected electrical and inhibitory signatures in the spontaneous activity. Thus, a strong shared background conceals local interactions that nevertheless have measurable effects on population statistics. This observation reinforces the limitations of inferring connectivity solely from activity^1^, while showing how perturbation-based maps can overcome this limitation.

Both patch clamp and local field potential (LFP) recordings previously reported state-dependent low-frequency oscillations^62,70–73;^ and traveling waves of brain activity have been reported in multiple species^74^. These prior measurements probed population ensemble activities measured either extracellularly as field potentials^63,75^, or optically via ensemble-average voltage imaging^70,71^. By measuring the subthreshold voltages at single-cell resolution, we unambiguously showed that traveling waves are collective traveling oscillations of membrane potentials of individual neurons.

The different correlation structures of voltage and spikes reveal subtle features of the input-output transformation at the network level. Spike threshold and firing rate have been shown to strongly alter the transfer of input to output correlations^76,77^. In our recordings, depolarizing excursions near spike initiation were less shared than fluctuations around or below resting potential, and self- and cross-spike-triggered voltage averages diverged shortly before a spike. These observations support a contribution of private depolarizing fluctuations to spike decorrelation, in line with prior intracellular observations and modeling^78,79^. The source of these private fluctuations remains unresolved. Cell-to-cell variations in spike threshold and in time-dependent subthreshold-to-spiking transformations likely contributed to further decorrelate spiking.

We observed strong representation of active whisking in the NPY^(+)^ but not the NPY^(-)^ neurons, suggesting different computational roles for these subpopulations. Remarkably, the subgroup-average subthreshold voltages were near-indistinguishable for NPY^(+)^ vs. NPY^(-)^. The differential behavioral modulation arose from a difference in behavioral sensitivity of the subthreshold-to-spiking transformations. Prior work showed behavioral-state-dependent modulation of NDNF interneurons^23^ and heterogeneity among NDNF populations across sleep and wakefulness^66^. Our results relate such heterogeneity to specific genetic subtypes and subthreshold-to-spiking transformations. The same arousal signal pushes the two subtypes in opposite directions.

Together, these findings demonstrate the value of characterizing a circuit at several interacting levels within the same experiment. The rich and time-dependent subthreshold-to-spiking properties of individual cells play an often-underappreciated role in converting shared synaptic input fields into individualistic firing patterns. Gap junction and synaptic connections together promote synchronous firing under abrupt-onset inputs, and suppress sustained network activity. Despite the presence of inhibitory couplings, most cell pairs showed positively correlated variations in spike rate, indicating a dominant influence of correlated drive. Combining measurements of distinct physiological mechanisms allows circuit models to be constrained by independently measured cellular and connection properties, rather than by fitting spontaneous activity alone. L1 provides a favorable setting for this approach, but the experimental logic applies more broadly to circuits in which optical access and genetic targeting permit joint perturbation and recording.

### Limitations and extensions

Several experimental limitations constrain the present scope. The main dataset comprises two mice, and the recorded cells sampled approximately 12% of NDNF-positive neurons within the volume bounded by the FOV. The connectivity maps omit unrecorded neurons and connections outside the tested distances or below the detection threshold. Scattered-light and spatial-footprint controls support the interpretation of detected responses, but validation by paired patch clamp *in vivo* was not performed. Finally, voltages were normalized to spike height, and optogenetic stimulation did not deliver calibrated current steps. Hence, the resulting excitability measures, PSP amplitudes, and effective firing relationships should be interpreted as semi-quantitative.

The principal technical opportunities are to increase photon efficiency, sampling density, and imaging depth while retaining independent optical stimulation. More sensitive, bright, fast, and red-shifted voltage indicators and improved light sources could expand the field of view or improve sensitivity, subject to constraints from bleaching and tissue heating. Combined confocal detection and targeted illumination^80,81^ offers a route to deeper recording, and multiplane^7,82^ or structured illumination^83^ approaches could sample a larger fraction of the local circuit. Two-photon holographic approaches could improve optogenetic targeting specificity and access deeper populations^84^. Combining these capabilities for multi-millimeter fields of view remains a substantial engineering task, and implementation of dense recordings may require advanced signal-extraction algorithms^85^. The molecular constructs, illumination strategy, and mapping and analysis protocols developed here can also be adopted separately as components of other all-optical experiments.

The tools introduced here open a broader opportunity to measure how cellular excitability, connection strength, and interaction kinetics change together during learning, altered behavioral states, or targeted molecular perturbations. Large-scale all-optical electrophysiology provides a route for relating circuit activity to fine-grained single-cell, synaptic, and gap-junctional properties.

## Acknowledgements

We thank Shahinoor Begum, Andrew Preecha and Camila Bodden for technical assistance. We thank Brent Doiron, Christoph Miehl, He Tian, Xiuyuan Li, David Wong-Campos, Pojeong Park, and Eric Moult for helpful discussions. Figures contain elements from BioRender. This work was supported by a Vannevar Bush Faculty Fellowship, the Harvard Brain Initiative, NIH grants R01-NS126043, R01-NS133755, and R01-MH117042, and Chan Zuckerberg Initiative Dynamic Imaging Grant 2023-321177. XW acknowledges support from an American Heart Association (AHA) postdoctoral fellowship.

## Author contributions

YQ designed and built the microscope, designed and performed the experiments, analyzed the data, and wrote the manuscript. VJP designed and built an earlier version of the microscope. XW performed the spinning disk confocal imaging. SEP and LDL synthesized and provided JF608-HTL. AEC supervised the project and wrote the manuscript.

## Data availability

Data are posted at: https://dandiarchive.org/dandiset/001960.

## Code availability

Data analysis scripts are at: https://github.com/adamcohenlab/Qi2026WideAreaL1

## Methods

### DNA constructs and AAVs

To combine optogenetics with voltage imaging, we co-expressed a blue-shifted channelrhodopsin, CheRiff, and a chemigenetic voltage indicator, Voltron1 or Voltron2, linked by a P2A ribosomal skip sequence^86^. We appended to Voltron a K_ir_2.1 membrane trafficking sequence^87^ and a K_V_2.1 soma localization sequence^88^. CheRiff was soma-localized in the same way. The Optopatch construct was inserted into an AAV backbone under a hSyn promoter and flanked by LoxP sequences for Cre-dependent expression. Genetic constructs used in this work are available at Addgene IDs 263112, 263113.

Optopatch AAV2/1 viruses were packaged either by Janelia viral core or UNC NeuroTools.

### Animals

All animal procedures were approved by the Institutional Animal Care and Use Committee (IACUC) of Harvard University and performed in accordance with institutional and NIH guidelines. Experiments used mice of both sexes aged P60–180 at the time of imaging. The NDNF-Cre::NPY-GFP double-transgenic reporter line was generated by crossing NDNF-Cre mice (Jackson Lab, Stock #028536) with NPY-hrGFP mice (Jackson Lab, Stock #006417); double-heterozygous offspring were used for imaging. Mice were group-housed before surgery and singly housed after cranial-window implantation on a 12 h:12 h light:dark cycle with food and water available *ad libitum*.

### Cranial window surgery and virus injection

The cranial window implantation procedure follows a previous protocol^89^. In short, a heterozygous NPY-GFP::NDNF-Cre mouse (P60-180) was deeply anesthetized by 3.5% isoflurane, then maintained in anesthesia by 1-1.5% isoflurane. A 3 mm diameter craniotomy was made over the left hemisphere (1.5 mm caudal from bregma, 3.2-3.5 mm from midline). Injections were made with a micropipette (World Precision Instruments, No. 504949) pulled with a pipette puller (Sutter Instruments, P1000). The pipette tip was broken to a diameter of 10 – 20 µm and beveled with a home-built pipette beveller. Virus AAV2/1 hSyn-DIO-Voltron-ST-P2A-somCheRiff was diluted to ∼3 x 10^12^ GC/mL with phosphate-buffered saline (PBS). Injections of 60 nL (30–60 nL/min) at depth ∼150 µm below the dura were performed at 10-14 different injection sites covering the entire exposed cortex. Following the virus injection, a cranial window was placed onto the cortex and glued to the skull with Vetbond (Patterson Veterinary, No. 07-805-5031). The cranial window was made by bonding two layers of 3 mm #1 round coverslip (Harvard Apparatus) and one 5 mm #1 round coverslip (Harvard Apparatus) with UV curable adhesive (Norland Products, NOA 81). The titanium head plate was bonded to the skull and all wounds sealed by dental cement (C&B metabond, Parkell, No. 242-3200). Animals were housed separately to avoid damage to the implant.

### Retroorbital injection

One day prior to imaging, 100 nanomoles of JF608-HTL was injected into the retroorbital sinus. To prepare the injection mixture, lyophilized JF608-HTL was dissolved in 20 µL of dimethyl sulfoxide (DMSO), 20 µL of Pluronic F127 (20% w/v in DMSO) and 60-80 µL of PBS.

### Free-running treadmill

We used a low-profile treadmill design by Janelia^90^, and modified it to float on three springs (McMaster, 9657K76) at the bottom to reduce walking-induced head motions.

### Behavioral videography

During imaging, the animal’s face was recorded to monitor whisking and arousal. Video of the face was acquired with a FLIR Grasshopper 3, GS3-U3-41C6NIR-C at 40 fps under infrared illumination using an LED source(Thorlabs, M850L3) and synchronized to the fluorescence acquisition via National Instruments DAQ, USB-6343. Both the LED source and the camera aperture were filtered by bandpass filters (Thorlabs, FB850-40). Mouse running speed acquisition was synchronized by the same DAQ via an optical rotary encoder (Broadcom, AEDM-5810) attached to the treadmill axle.

### Head-fixation and leveling

To image a large FOV, the cranial window must be precisely parallel with the focal plane of the microscope objective. To achieve this, the head restraint was mounted on a two-axis goniometer (Misumi Inc, B54-60UN, B54-60U2N). Prior to imaging, the head-fixed animal was placed under a home-built Michelson-Morley interferometer consisting of a 5 mW 630 nm laser pointer, a mirror (Thorlabs, BB1-E02), and a glass slide (VWR, No. 16004-422) mounted at precisely 45° as the beam splitter. The cranial window acted as the second mirror. The two-axis goniometer was adjusted until the two reflected beam spots overlapped on a white target ∼4 m away.

### Microscope

The home-built wide-area microscope consists of a pattern projection system, and a large field of view (FOV) epifluorescence microscope.

Two LED light sources, blue (Luminus, SBT-70-B) and orange (OSRAM, GY CS8PM1.23 for JF585; or Luminus, PT-54-RAX-L35 for JF608) in custom housings were combined via a custom-sized dichroic mirror (Semrock, Di01-R488/543/635, 60x85 mm) and reimaged onto a digital micromirror device (DMD) (Vialux, V-7001; 1024 X 768 pixels, 13.7 μm pitch). The orange LED was cooled by a Peltier module (Sheetak, SKHC1-199-16-T100-SS-TF00-ALO) to achieve desired illumination power. A total-internal-reflection prism (Young Optics Inc, 72.CAC14G001) was used to direct the light onto the DMD chip. Patterned light was collected by a 100 mm tube lens (Zeiss, Makro-Planar 100 mm), directed into a 50 mm objective lens (Olympus, MVPLAPO 2XC, NA 0.5) via a custom-sized dichroic mirror (Alluxa, 504/605 ULTRA Dualband Dichroic Beamsplitter, 60x85 mm^2^, 5 mm substrate for JF585; or Chroma, zt488/607rpc-UF3, 60x85 mm^2^, 3 mm substrate for JF608). Fluorescence emission was filtered by a custom-sized emission filter (Semrock, FF02-641/75, 60 mm diameter for JF585; or FF01-709/167, 60 mm diameter for JF608), and reimaged onto the camera (Teledyne Photometrics, Kinetix, 3200 x 3200 pixels) through a 50 mm tube lens (VS Technology, VS-50085/M42). All microscope hardware was synchronized by a data acquisition device (National Instruments, USB-6343) and controlled by custom-written software in MATLAB.

For high magnification high NA imaging, we changed the objective lens to XLPLN25XSVMP2 (Olympus).

### Targeted illumination

We developed a semi-automated pipeline to acquire high-contrast Hadamard digital confocal images, segment the images to locate neuron cell bodies, and generate targeted illumination patterns on the DMD. All calculations were performed in MATLAB.

### Hadamard microscopy

Hadamard digital confocal images were acquired as previously described^33^. In brief, the pseudo-random illumination patterns were generated with the last 63 columns of a size 64 normalized Hadamard matrix, corresponding to 64 illumination patterns for DMD 63 pixels, and this pattern is tiled for each row of DMD pixels. The patterns for the adjacent rows are offset by 13 pixels. A calibration dataset was acquired on a thin fluorescent sample under orange illumination. The sample consisted of a layer of pink neon Sharpie (Newell Brands) sandwiched between a three-layer stack of #1 coverslips (VWR, No. 48393-059) bonded by UV curable adhesive (matching the *in vivo* condition) and a 1 mm glass slide (VWR, No. 16004-422). Then the same patterns were projected and recorded on the live sample with ten repeats. The repeats were then reconstructed and averaged, yielding the final confocal image.

### Image segmentation and pattern generation

The Hadamard confocal image was filtered by a 7 x 7 pixel mean-subtracted Gaussian filter with standard deviation of 7 pixels, then segmented using the watershed algorithm. Watershed regions fewer than 10 pixels were discarded. The putative neuron in each watershed region was located using the maximum intensity pixel. We then cross-correlated the original unfiltered Hadamard neuron images with the Gaussian filter, and selected the 320 putative neurons with the highest cross-correlation values. We generated a 40 μm wide disk-shaped illumination mask for each neuron.

### Voltage imaging

For all imaging experiments, targeted neurons were continuously imaged at 787 Hz under orange illumination at ∼16 mW/mm^2^.

### Temporal multiplexing for independent dual color illumination

To achieve independent patterning of two-color illumination by the same DMD, we flickered two DMD patterns within each camera exposure and synchronized each light source to one of the patterns. 80% of each exposure was allocated to the orange pattern/light pair, and 20% to the blue pattern/light pair.

### Characterizing blue light off-target stimulation

To test for off-target stimulation by scattered blue light, we stimulated the entire FOV for 1 s using a square lattice of 35-44 blue spots (20 μm wide disk, 2.5 mW/mm^2^) with 400 μm lattice spacing and a random lateral offset for each trial, for a total of 1000 trials.

### Mapping intrinsic excitability

In each trial, a random sparse subset of cells was selected for stimulation. No two stimulated cells were separated by less than 800 μm. Selected cells were stimulated by four 500-millisecond-long steps of increasing blue light intensity (1.5–6 mW/mm^2^, equal increment), with 500-millisecond rest between each stimulation, followed by a 5-second-long linear ramp (0–6 mW/mm^2^). Blue light was targeted at the soma (20 μm wide disk). 300 trials were conducted per session. We recorded the voltage of all cells throughout the session.

### Mapping synaptic connectivity

In each trial, a random sparse subset of cells was selected for stimulation (minimum pairwise separation 800 μm), the same way as above. Selected cells were stimulated by a 20 ms pulse (10 mW/mm^2^) targeted at the soma (20 μm wide disk) followed by 180 ms rest. 14400 trials were conducted per session so that each cell was stimulated 243-480 times. We recorded the voltage of all cells throughout the session.

### Mapping gap junction connectivity

Cells were stimulated with 50% duty cycle pseudo-random Hadamard-Walsh codes, to evoke uncorrelated spiking patterns across the population. In each trial, stimulated cells received a 750-millisecond-long step (4 mW/mm^2^) targeted at the soma (20 μm wide disk). To determine which subset received stimulation, we assigned 320 cells to a column of a 320 x 320 Hadamard matrix: +1 cells received stimulation; -1 cells did not. Thus a 320 x 320 Hadamard matrix generated 320 trials. The full set of trials was repeated 4 times per session. To avoid continuous stimulation of any cells, we elementwise multiplied the Hadamard matrix with a pseudorandom mask (50:50 ±1). To test for crosstalk from *en passant* neurites, we repeated the same measurement under the high-magnification objective (7.5x system magnification, NA 1.0, XLPLN25XSVMP2, Olympus). 16 cells were imaged per FOV, and each cell was stimulated with a unique code drawn from a 16x16 Hadamard matrix.

### Correcting mechanical drift

Each imaging session lasted 60 - 150 minutes. Without intervention, neurons sometimes drifted out of the illumination masks. To mitigate this drift, we took a Hadamard confocal image every 6 minutes. Lateral drifts were measured by the Farneback optical flow algorithm. We then added the lateral offset to the illumination mask. To measure axial drift, we first calculated the correlation between the fast Fourier transform (FFT) of the Hadamard confocal image and the FFT of a 60 μm deep confocal z-stack (3 μm resolution). The most correlated depth was found by fitting the correlation with a quadratic polynomial. We then moved the objective to correct the axial drift. The z-stack was taken prior to the connection mapping trial.

### Spinning disk confocal structural imaging

A 3D volume of all optically accessible L1 was acquired using a custom-built two-channel spinning disk confocal microscope (described in a separate publication^80^) at 8x magnification and 500 nm z-step size, while the mouse was under isoflurane anesthesia.

### Data analysis

All analysis was performed in MATLAB.

### Statistics

Unless otherwise stated, summary statistics are reported as mean ± s.d., and *n* denotes the number of cells, cell pairs, or mice, as specified in each figure legend. Correlations were quantified with Pearson’s correlation coefficient. Group comparisons (e.g., NPY^(+)^ vs. NPY^(-)^ cells) used the two-sided Wilcoxon rank-sum test unless otherwise specified. The statistical significance of individual synaptic and gap-junctional connections was assessed against per-pair null distributions at a threshold of p < 0.05, with Benjamini-Hochberg correction across all tested pairs (see below). No statistical methods were used to predetermine sample size; sample sizes were as large as practical within the constraints of the experimental setup. Cells and cell pairs, rather than animals, are the unit of analysis throughout. With N = 2 mice the design does not support inference at the level of a population of animals.

### Signal extraction

We adapted recent approaches to signal extraction^91,92^ to suit our low-magnification targeted-illumination imaging setup. We first corrected in-plane motion by applying the Farneback optical flow algorithm on temporally smoothed movies (2^nd^ order Savitzky-Golay filter with 9-frame window). We next detrended the movie using predetermined exponential functions via linear regression. For a small ROI (13-pixel wide disk) around each cell, we then modeled the fluorescence movie as

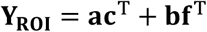

where **Y**_ROI_ is the vectorized movie (d × T matrix), **a** ∈ R^d^ is the spatial footprint of the neuron, **c** ∈ R^T^ is the fluorescence signal of the neuron, **b** = [b_1_ b_2_ … b_n_], b_i_ ∈ R^d^ is the spatial footprint of the background fluorescence (other than the cell of interest, i.e. fluorescence of nearby neurons, neuropil, etc.), and **f** = [f_1_ f_2_ … f_n_]^T^, f_i_ ∈ R^T^ is the time-dependent fluorescence signal of the background.

We first located each neuron by segmenting the digital confocal image. We initialized **a** as the first principal component of the background-removed **Y**_ROI_. We calculated the background by first dividing the full movie into a 2x3 grid of sub-regions then spatially averaging the sub-regions into 6 fluorescence traces. The background was removed by linear regression. The cell footprint **a** was then further refined by two iterations of spike-time regression^91^.

Nonnegative matrix factorization (NMF) can effectively separate spatially overlapping non-negative signals such as Ca++ dynamics^93^. However, voltage signals have fast spikes and slow fluctuations that can be both positive and negative. To overcome this, we sought to apply NMF separately on the fast and slow components, and positive and negative halves. We decomposed **Y**_ROI_ into four components: high-frequency positive, high-frequency negative, low-frequency positive, and low-frequency negative,

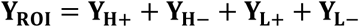

where the low-frequency component is produced by a moving median filter with a 60-frame moving window, and the high-frequency component is

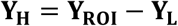

Each component was then modeled as

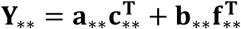

We reasoned that 1) all footprints should be the same for both positive and negative components, 2) **a** should be the same for all frequency components, 3) **b**_H∗_ contains only nearby neurons, and 4) **b**_L∗_ has one additional component to capture the diffuse background (i.e. out-of-focus fluorescence and neuropil). We first ran non-negative matrix factorization (NMF) on **Y**_H+_ to solve for **a**, **c**_H+_, **b**_H∗_, **f**_H+_. We then solved for C_H−_, **f**_H−_ by running NMF on −**Y**_H−_ while fixing **a** and **b**_H∗_. To solve for **c**_L∗_, **b**_L∗_ and **f**_L∗_, we appended a constant term to **b**_H∗_ as the initialization for **b**_L∗_ , then ran NMF on **Y**_L+_ and −**Y**_L−_ . Finally, we reconstituted neuronal fluorescence signal c as

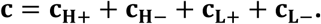

### Correcting motion artifacts

Due to the targeted illumination, lateral motion led to changes in neuron fluorescence. For a point emitter, lateral motion samples the illumination profile. Approximating the illumination profile near its center as a two-dimensional quadratic function, the motion-induced fluorescence change is approximately:

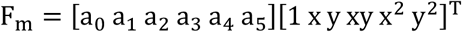

where x, y ∈ R^T^ describe the motion trajectory in 2D and *a*_i_ are coefficients that describe the shape of the paraboloid. However, the motion of each cell is never tracked perfectly. To account for this, we used PCA to generate a motion basis set from the motion of all cells. To avoid overfitting, we truncated PCA to only top 10 PCs, and then kept only the PCs that explained 80 percent of the total variance. The resulting basis set has the form:

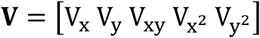

where V_x_ ∈ R^n×T^ is the table of *n* PCs calculated from the x-motion of all cells, and similarly for V_y._ We then decomposed **V** and **c** into high- and low-frequency components and regressed **V** from **c** in the two frequency bands separately. The regression weights were calculated on the running epoch of the dataset, when the motion artifacts were largest.

### Detecting spikes, estimating false positive/negative rates, and calculating SNR

Spike detection was adapted from Volpy^91^. In brief, we first generated a template spike waveform for each cell by averaging its 500 largest spikes. This template was then used as a matched filter and applied to the fluorescence traces. Spikes were then detected by thresholding the filtered trace. We estimated the noise distribution using the negative portion of the filtered trace, and chose a threshold that set the false-positive rate to ∼500 spikes per hour. To estimate the false positive and negative rates, we modeled the supra-threshold peaks as a mixture of two truncated (by the detection threshold) Gaussian distributions for supra-threshold noise (false positive) and detected spikes (real positive) respectively. We estimated the parameters of these truncated Gaussians via maximum likelihood estimation. We defined spike height as the mean of the spike Gaussian distribution, and the fluorescence trace was normalized by this spike height. We defined SNR as the ratio between the spike amplitude and shot noise. We estimated the shot noise as standard deviation of the 1-frame difference of the fluorescence trace. Standard deviation was approximated as 1.4826 times the median absolute deviation to avoid contamination from spikes.

### Face movie analysis

Whisking motion was extracted from the facial videography by first averaging pixels in a manually defined region around the whisker pad. We then defined motion as the absolute 1-frame difference of the pixel-average trace. The trace was then smoothed by an 8-frame moving median filter.

We observed that the log-transformed whisking motion has a bimodal gaussian distribution, corresponding to quiet and whisking epochs. We fitted a gaussian mixture model to the log-transformed whisking motion, and set the whisking threshold at the cross-over point of the two gaussian distributions. We defined whisking epochs to be whenever the whisking motion was greater than this threshold, and quiet epochs to be the exclusion of the whisking epochs extended by 100 frames before and after each whisking epoch.

### Characterization of blue light off-target effects

After stimulating the entire FOV with square lattice of blue spots with 400 μm lattice spacing and a random lateral offset for 1000 trials, for each neuron, we assigned stimulation spots to 20 distance bins (15 μm increment) and calculated 20 stimulation triggered average (STA) voltage and firing rate. We then extracted voltage and firing increase from the STAs. We define voltage increase to be the average voltage 60 ms to 190 ms after stimulation onset minus the average voltage -130 ms to 0 ms before the onset. We defined the firing increase to be the average firing rate 0 ms to 130 ms after stimulation onset minus the average firing rate -130 ms to 0 ms before the onset.

### Removing blue light fluorescence crosstalk

Fluorescence excited by blue light can bleed into the JF608 channel, but this fluorescence is simply a scalar multiple of the blue light waveform. We determined the unknown multiplicative factor by calculating the regression coefficient between the 1-frame difference of the fluorescence signal and the 1-frame difference of the blue light waveform during the step stimulus, then subtracted the scaled blue light waveform from the fluorescence signal. Across experiments mapping intrinsic excitability properties and connectivity, the blue light crosstalk effect was 0.3 ± 0.2 (mean ± s.d.) of a spike height.

### Calculating neural intrinsic excitability properties

We measured adaptation by taking the difference between the onset firing rate (average of the initial 50 ms) and the steady state firing rate (average of the last 250 ms) normalized by the steady state firing rate. We found adaptation increased with light dose and plateaued at the third of the four steps. We thus used the adaptation calculated on the third step stimulus.

During the slow ramp, we defined rheobase as the voltage at which the 10^th^ spike occurred. Although this is slightly above the typical *in vitro* rheobase, we found this definition more robust against spontaneous activity fluctuations. We defined the maximum firing rate as the average firing rate in the last 600 ms of the ramp.

We extracted the membrane time constant by first calculating the off-step triggered average voltage, then fitting an exponential function to the average decay.

For after-depolarization, we calculated the average spike waveform using isolated spikes only (no other spike within ±20-frame window), then defined ADP as the difference between the 10-frame averages centered at frames -11 and +7 relative to the spike peak. To divide cells into three intrinsic property bins, we first z-score normalized each of the five properties then performed hierarchical clustering on z-scored properties using Ward’s method.

### Analysis for mapping synaptic connections

We found that blue illumination flashes evoked stereotyped depolarizations waves that propagated across the FOV in ∼100 ms, possibly due to visual detection of the blue flash. We calculated the effect of this artifact for each cell *j* by averaging the response of cell *j* to trials targeted to faraway cells (> 800 µm). We then subtracted this stimulus artifact from the trial-average IPSP of cell *j* (independent of presynaptic cell *i*). For each presynaptic cell *i*, the trial-average IPSPs of its postsynaptic cells had a correlated component similar to spike triggered averages. We estimated this as the median IPSP of postsynaptic cells between 300 and 400 μm away, then subtracted this signal from the IPSPs of all postsynaptic cells. To both detect synaptic connections and extract IPSP parameters (amplitude and decay time constant), we modeled the empirical IPSP waveform (*y_ij_*) for connected pairs as a sum of a double exponential (pure IPSP), a quadratic baseline term, and a blue light excitation term (*H*_1_), while for unconnected pairs, we did not include the double exponential term (*H*_0_). For connection detection, we calculated a connection statistic: *T_ij_* = *max*{0, *RSS*_0_ − *RSS*_1_}, where *RSS*_0_ = ǁ*y_ij_* − *H*_0_ǁ^2^ , and *RSS*_1_ = ǁ*y_ij_* − *H*_1_ǁ^2^ . We then generated a 10000-element null distribution of IPSP waveform for each cell pair (i,j), by randomly sampling *N* pulses targeted to cells > 800 μm from cell *j*, where *N* equals the number of trials which probed the (i,j) connection. With the null IPSP waveform, we calculated the null distribution of the connection statistic. The statistical significance threshold was set to 0.05 with Benjamini-Hochberg correction for connection detection.

The 20-ms stimulus pulses typically evoked stereotyped bursts of spikes, but the spikes in these bursts were sometimes difficult to detect on top of the large baseline depolarization. We estimated the average number of evoked spikes per trial by first calculating the trial average frequency spectrum during the 20 ms stimulus, then multiplying the average frequency by 20 ms. For cells that did not show consistent optically evoked bursts, we used our standard spike detection algorithm, above.

We explicitly included the possibility that optical stimulation evoked spikes in non-targeted cells via scattered blue light, and that these spikes then evoked IPSPs in other cells. To incorporate non-target spiking, we modeled our IPSP matrix as a matrix product of the firing rate matrix (containing optically evoked spikes in targeted and non-targeted cells) and an unknown connectivity matrix. We solved the unknown matrix via lasso regression (Supplementary Fig. 16). Fewer than 20% of corrected IPSPs differ more than 20% from the raw IPSPs, so for simplicity we used the connectivity matrix calculated without correction for scattered light.

### Analysis for mapping gap junction connections

We calculated spike-triggered cross-average voltage or spike rate as

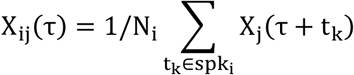

where N_i_ is the number of spikes of cell *i*, spk_i_ is the collection of spike times of cell *i*, X_j_ is either the voltage or binary spike train of cell *j*, and −200 ms < τ < 200 ms.

For each cell *i*, we found slow correlated dynamics in the cross STA voltages of all cells *j*. These dynamics had a distance-dependent and a global component. We estimated the distance-dependent component by first averaging cross STA voltage of cells within 200 μm thick annulus at distance *r* from cell *i*, then fitting a 3^rd^ order polynomial as a function of *r*. We removed the distance dependent component by subtracting the fit from the individual cross STA voltage of cell *j* at distance *r* from cell *i*. We then estimated the global component as the average cross STA voltage of cells > 800 μm from cell *i*, then regressed it out from the cross STA voltage of each cell *j*. We excluded frames {-2, -1, 0, 1, 2} when calculating the regression coefficient to preserve the spikelet.

We defined spikelet amplitude as the difference between the average of frames {0, 1} and the average of frames {-4, -3, -2, 4, 5, 6} of the cross STA voltage trace, where the frame indices were counted relative to the spike peak. We generated a 5000-element null distribution for the spikelet for each cell pair (*i*, *j*) by randomly sampling *N* spikes from cells > 800 μm from cell *j*, where *N* equals the number of spikes in cell *i*, then extracting the spikelet from the null STA voltage trace. The statistical significance threshold was set to 0.05 with Benjamini-Hochberg correction.

To quantify the effect of fluorescence scatter on the spikelet measurement, we calculated the cross STA fluorescence of interstitial pixels within 400 μm of each triggering cell. The STA fluorescence was then normalized by the spike fluorescence of the triggering cell. The spike fluorescence was defined as the difference between frame 0 and -2 (relative to spike peak) of the STA fluorescence. We extracted the scatter spikelet amplitude the same way as we did for gap junction-mediated spikelets.

### High magnification spikelet footprint analysis

To test whether the putative gap junctional spikelets truly represented membrane voltage changes in the post-junctional cells, we analyzed the footprints of these events. For genuine gap-junctional connections, the post-junctional spikelet and full spike footprints should match. For scattered fluorescence or *en passant* neurites, these footprints would differ.

To extract the spikelet footprints, we calculated the cross STA fluorescence for each pixel containing the post-junction cell (cell *j*) using the spikes of pre-junction cell (cell *i*) as trigger. We normalized this footprint by the spike height of cell *j* at each corresponding pixel, then extracted the spikelet amplitude from the STA trace for every pixel. We modeled this spikelet footprint as a weighted sum of the cell *j* spike footprint and a flat background (Supplementary Fig. 18), and compared the relative amplitudes of these two contributions.

### High resolution confocal z-stack segmentation and registration to 1x-magnification functional data

We first fine-tuned the Cellpose 3.0 model^94^ on a manually labeled training set, then verified the final segmentation results by visual inspection.

To register the low-magnification functional data onto the high-resolution confocal z-stack, we first calculated a 2d affine transformation using blood vessels as fiducials, then manually selected and registered ∼500 fiducial cells, and finally calculated a 4^th^ degree polynomial transformation and applied it to the low-magnification cell coordinates. We then registered the transformed low-magnification 2d cell coordinates to the 3d z-stack cell coordinates using the iterative closest point algorithm. We identified each low-magnification cell coordinate with the closest z-stack coordinate, with a 15 μm distance cutoff. We determined co-localization of GFP and Voltron-JF608 by visual inspection.

### Linear mixed effects model for synaptic properties and pre-/post-synaptic identity variance

We observed that both synaptic amplitude, *w_ij_*, and IPSP decay time constant, *τ_ij_*, followed log-normal distributions. We therefore modeled *w_ij_* as:

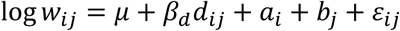

where

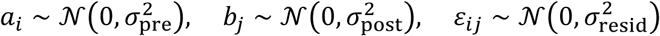

and μ is a mean, *d*_ij_ is the distance between cells *i* and *j*, *a*_i_ is the presynaptic random effect for cell *i*, and *b*_j_ is the postsynaptic random effect for cell *j*, and *ε*_ij_ is the model residual, i.e. the synapse-specific contribution to the weight. We modeled synaptic time constants *τ_ij_* in the same way.

IPSP decay time constant had much larger uncertainty than amplitude from IPSP waveform fitting described above. When fitting the linear mixed effects model for the IPSP decay time constant, we weighted each log *τ_ij_* by

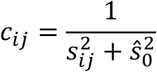

where *s_ij_* is the standard error of log *τ_ij_* estimated using the delta method, and 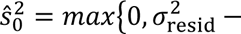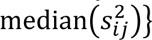, where 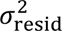 was estimated first using an unweighted linear mixed effects model fit. Uncertainty of variance parameters (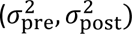) was estimated by simulating log *y_ij_* (or log *τ_ij_*) then refit the model on simulated data (parametric bootstrap). We repeated the bootstrap simulation 2000 times.

### Subthreshold voltage extraction

To extract subthreshold signals from the fluorescence trace of each cell, *F*, we first replaced the frames {-2, -1, 0, 1, 2, 3, 4} around each spike by those of *F*_f_, where *F*_f_ is *F* filtered by a 16-frame moving median filter, then applied a 17-frame 2^nd^ order Savitzky-Golay filter.

### Wave propagation analysis (speed and direction)

A wave propagation event results in a sharp rise in the population-average voltage, which we detected by finding peaks in the first derivative of the average voltage (*d̄V*/*dt*). Since these events only happen during the quiet epoch, we set the peak threshold to be 3.5x standard deviation of *d̄V*/*dt* during the whisking epoch. We restricted the analysis of each event to the time points between the minimum and maximum of voltage flanking the peak derivative. To find the average propagation speed for each event, we first calculated the subthreshold cross-correlation between all cell pairs. We then found the lag-time with the maximum correlation for each pair and calculated the slope of the line relating lag-time to separation for all cell pairs.

To find the flow field of each wave propagation event, we first calculated the Hilbert transform of the subthreshold voltage of each cell, then spatially smoothed this signal with a 325 μm wide Gaussian kernel and down-sampled onto a rectangular grid at 160 μm resolution. We then calculated the time average of this signal. The Hilbert-transformed signal has both an amplitude and a phase, and the flow field is the spatial gradient of the phase values.

### Oscillation peak detection

We first band-pass filtered (butterworth) the population average voltage between 1.5 and 10 Hz, then detected peaks with values greater than 0 from the quiet epoch.

### Frequency spectrum

The frequency spectrum of the subthreshold was estimated by the Welch method. The population-average spectrum was the average power spectrum across all cells.

### Dimensionality of subthreshold fluctuations

We first computed the covariance matrix of subthreshold voltage as

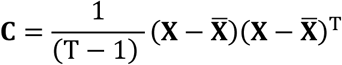

where **X** ∈ R^N×T^, N is the number of cells, T is the number of time points, **X** is the subthreshold voltage, and ̄X is the time average of **X**.

When the voltage is not smoothed, we estimated its shot-noise-free variance using the lag-1 variance:

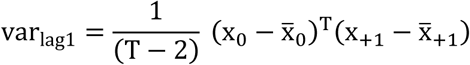

where x_0_, x_+1_ ∈ R^T−1^, x_0_ is the voltage without the last time point, and x_+1_ is the voltage without the first time point. We then replaced the diagonal of *C* with var_lag1_. Finally, we computed the eigenvalues of C and normalized them by the total variance.

### P(spk|V) for spontaneous and optically evoked firing

We calculated the empirical spike probability conditioned on subthreshold voltage as

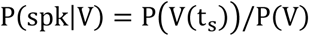

where V(t_s_) is the subthreshold voltage at spike times and P(V) is the distribution of all subthreshold voltages. We then converted this probability to rate by dividing it with the exposure time, 1.27 ms. For the linear ramp stimulus epoch, we only used the first half of the ramp to avoid the firing rate saturation in the latter half.

### P(spk|dV/dt) for spontaneous epochs

We calculated the empirical spike probability conditioned on first derivative of the subthreshold voltage as

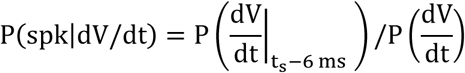

where dV/dt is shifted 6 ms relative to spike times, and we restricted V to between 0.2 and 0.8 spike height. We then converted this probability to rate by dividing it with the exposure time, 1.27 ms.

### Average firing rate and average burst rate

We defined average firing rate as the total number of spikes divided by the recording duration. To measure the influence of ADP on spontaneous spiking, we defined burst rate as the average value of the autocorrelogram from lag 2.5 ms to 15 ms (same time window as ADP).

### Estimating cross STAs of voltage or firing rate

To assess the contribution of synaptic or gap junctional connectivity to the cross STA voltage or firing rate, we first binned postsynaptic cells into quintiles by the evoked voltage amplitude (either IPSP or spikelet). We then calculated the average cross STA of cell pairs in each quintile. We found that these average STA were dominated by correlated inputs to the two cells. We then estimated these shared dynamics of each quintile by calculating the average cross STA of unconnected cell pairs, which we defined for each connected cell pair (i,j) as the pair (i,k), where cell k was less than 400 μm from both cell i and j, and unconnected to cell i. We then regressed the shared dynamics from the quintile average cross STA:

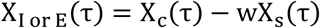

where

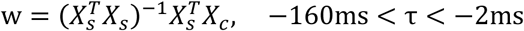

X_I or E_ is the contribution of either synaptic or gap junction connections to the average cross STA voltage or firing rate, X_c_ is the average cross STA voltage or firing rate of connected cell pairs, and X_s_ is the average cross STA voltage or firing rate of unconnected cell pairs.

### Effect of connections on voltage and spike rate

To quantify the contribution of synaptic or gap junctional connectivity in the cross STA voltage or firing rate, we defined:

1. Voltage suppression (due to an IPSP) as the difference between the average of frames 12 to 25 and the average of frames -17 to -4 of the cross STA voltage (all times measured relative to the triggering spike peak);
2. Rate suppression (due to an IPSP) as the difference between the average of frames 4 to 10 and the average of frames -10 to -4 of the cross STA firing rate. Relative rate suppression was normalized by the average of frames -10 to -4 of the cross STA firing rate;
3. Instantaneous voltage increase (due to a spikelet) as the difference between the average of frames 0 to 1 and the average of frames {-4, -3, -2, 4, 5, 6} of the cross STA voltage;
4. Synchrony (due to spikelets) as difference between the maximum of frames -1 to 1 and the average of 3 frames either before frame -1 or after frame 1, whichever is larger, of the cross STA firing rate. Relative synchrony was normalized by the average of 3 frames either before frame -1 or after frame 1, whichever is larger, of the cross STA firing rate.

### Lasso regression for whisk event detection

To find the onset of each whisking event, we modeled the whisking motion as a convolution between a motion kernel and an event variable, and solved for the event variable via sparse deconvolution:

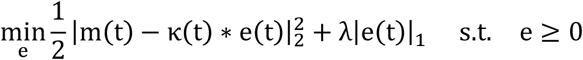

where *m* is the whisking motion, *e* is the event variable, κ is a 200-ms-long step function. We found the event onset by detecting the zero-crossings of e(t). A series of λ ranging from 1 to 15 were tested. We picked the λ that minimized the variance of whisking motion during the 2 s prior to the onset.

### Cell type classification using optogenetically measured intrinsic properties

To assess whether measured intrinsic properties can distinguish NPY(+) and NPY(-) subtypes, we trained a logistic regression classifier on 80% of the cells, then tested the classification on the remaining 20%. We measured the classification performance via balanced accuracy defined as

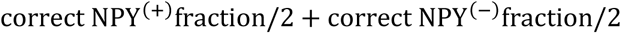

To estimate a 95% confidence interval on the balanced accuracy, we resampled the cross-validation fold 10,000 times, then used the 2.5^th^ and 97.5^th^ percentiles as the 95% interval. For control, we performed 5-fold cross-validated logistic regression using shuffled subtype labels.

### Clustering of cells by spontaneous firing patterns

We calculated the time-dependent firing rate for each cell by applying a 500-ms moving average to the binary spike train. We then z-scored the firing rate along the time dimension. To divide the population into two functional groups, we embedded the z-scored firing rate onto two UMAP dimensions, then performed k-means clustering with k = 2. To sort the raster plot, we projected the 2d UMAP onto the linear discriminant axis then sorted the 1d projection.

## Declaration of generative AI and AI-assisted technologies in the manuscript preparation process

During the preparation of this work, the author(s) used ChatGPT and Claude to help structure the Introduction and Discussion and to check the manuscript for errors or omissions. The author(s) reviewed and edited the output as needed and take full responsibility for the content of the published article.

