## Supplementary Figures for "Large-scale all-optical electrophysiology links excitability and connectivity to circuit dynamics in vivo"

**Supplementary Movie 1:** Spontaneous voltage dynamics in 320 Layer 1 interneurons and simultaneously recorded mouse face motion. Movie slowed down 25x from real time.

**Supplementary Movie 2:** Targeted optogenetic stimulation of a randomly selected subset of 76 out of 320 simultaneously recorded L1 interneurons. Plots show the voltage dynamics of non-stimulated neurons (top) and stimulated neurons (bottom). Movie slowed down 13x from real time.

**Supplementary Movie 3:** Population-averaged spatiotemporal interaction kernel, comprising rapid excitatory gap-junction coupling and delayed synaptic inhibition. Movie slowed down 125x from real time.

**Supplementary Movie 4:** Subthreshold wave propagation during quiet-state spontaneous activity. Movie slowed down 25x from real time.

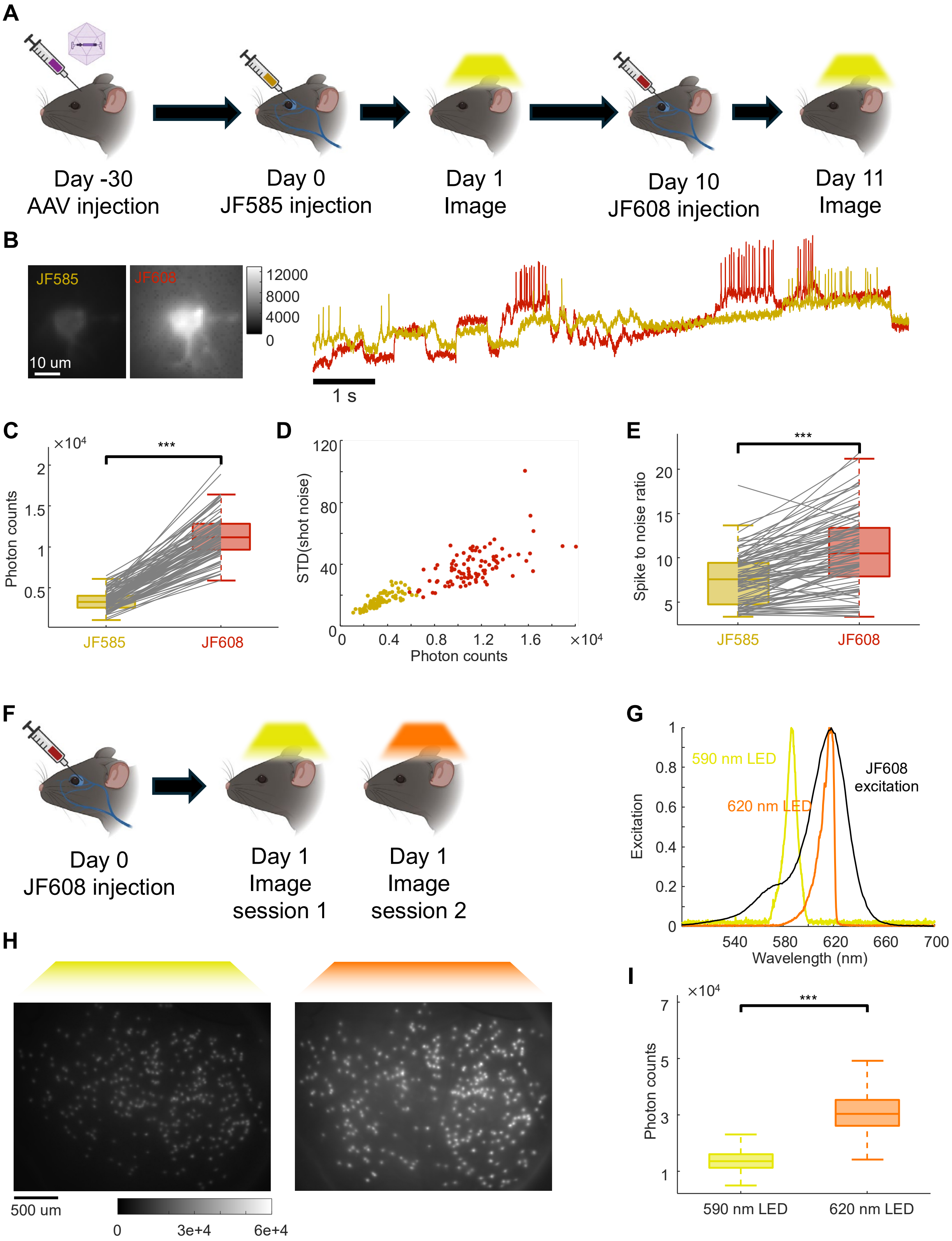


**Supplementary Figure 1. Optimizing dye and LED for *in vivo* Optopatch.**

(A) Experimental timeline for *in vivo* comparison between JF585-HTL and JF608-HTL.

(B) *In vivo* fluorescence of a cell expressing Optopatch construct stained by either JF585-HTL or JF608-HTL under the same illumination intensity by a 590 nm LED.

(C) *In vivo* photon counts from JF585-HTL vs. JF608-HTL under the same illumination condition, n = 92 cells.

(D) Shot noise vs. photon counts for JF585 and JF608.

(E) Comparison of spike-to-noise ratio of the same cell stained sequentially with JF585-HTL, then JF608-HTL, n = 91 cells. Single-sided Wilcoxon rank sum test. *, **, ***, indicate $0.01 < p \leq 0.05$, $0.001 < p \leq 0.01$, and $p \leq 0.001$.

(F) Experimental timeline for *in vivo* comparison between 590 nm and 620 nm LED.

(G) Spectra of 590 nm and 620 nm LEDs and the absorption spectrum of JF608-HTL.

(H) *In vivo* fluorescence of the same FOV illuminated with matched illumination intensities by either 590 nm or 620 nm LEDs.

(I) Quantification of the single-cell intensities in (H). Single-sided Wilcoxon rank sum test for statistical significance. Box plots show median; 25$^{th}$ and 75$^{th}$ percentiles; whiskers show 1.5x inter-quartile range.

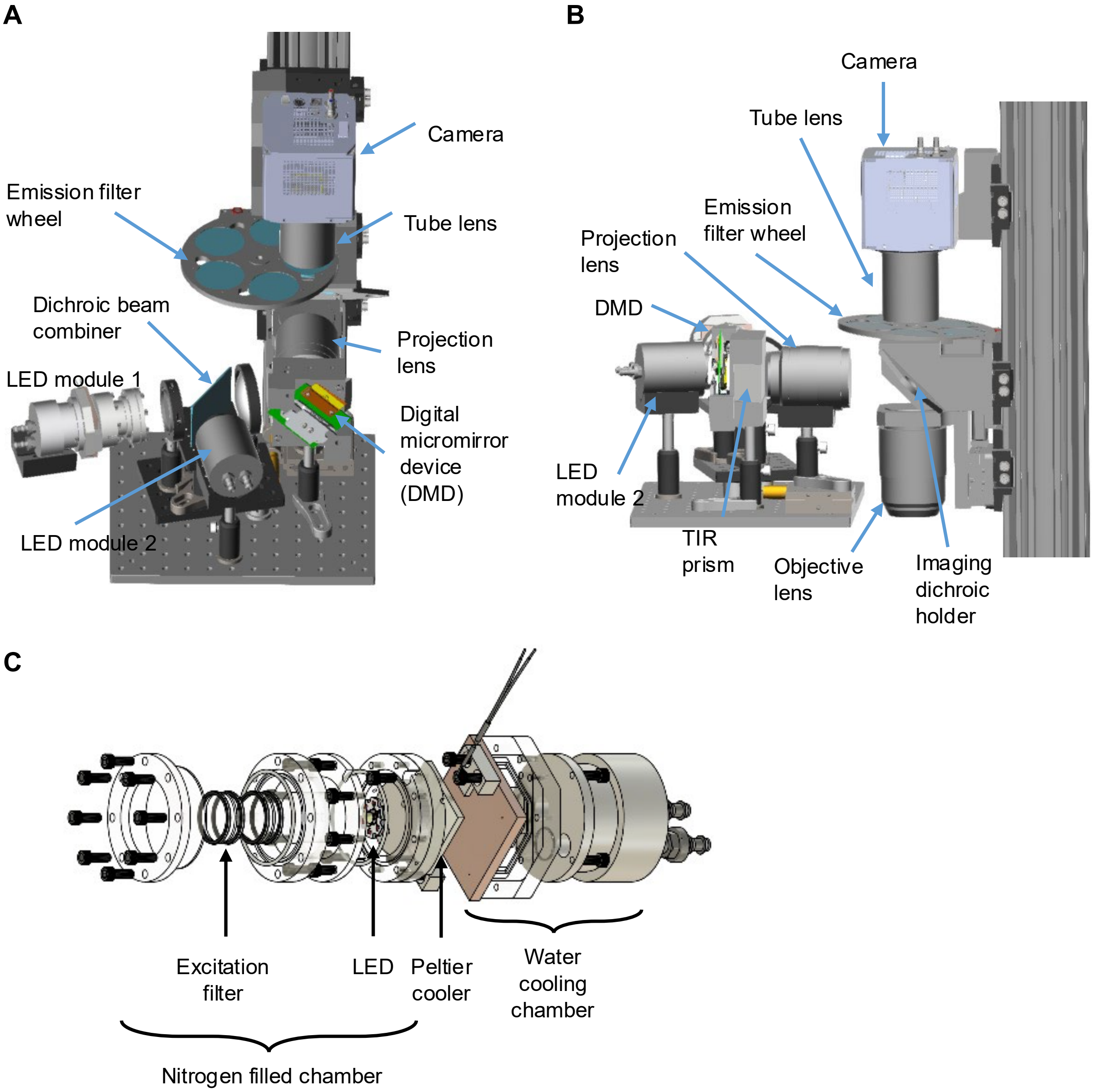


**Supplementary Figure 2. Microscope design.**

(A, B) Front and side views of the microscope.

(C) Custom LED chamber for active cooling to -20 °C.

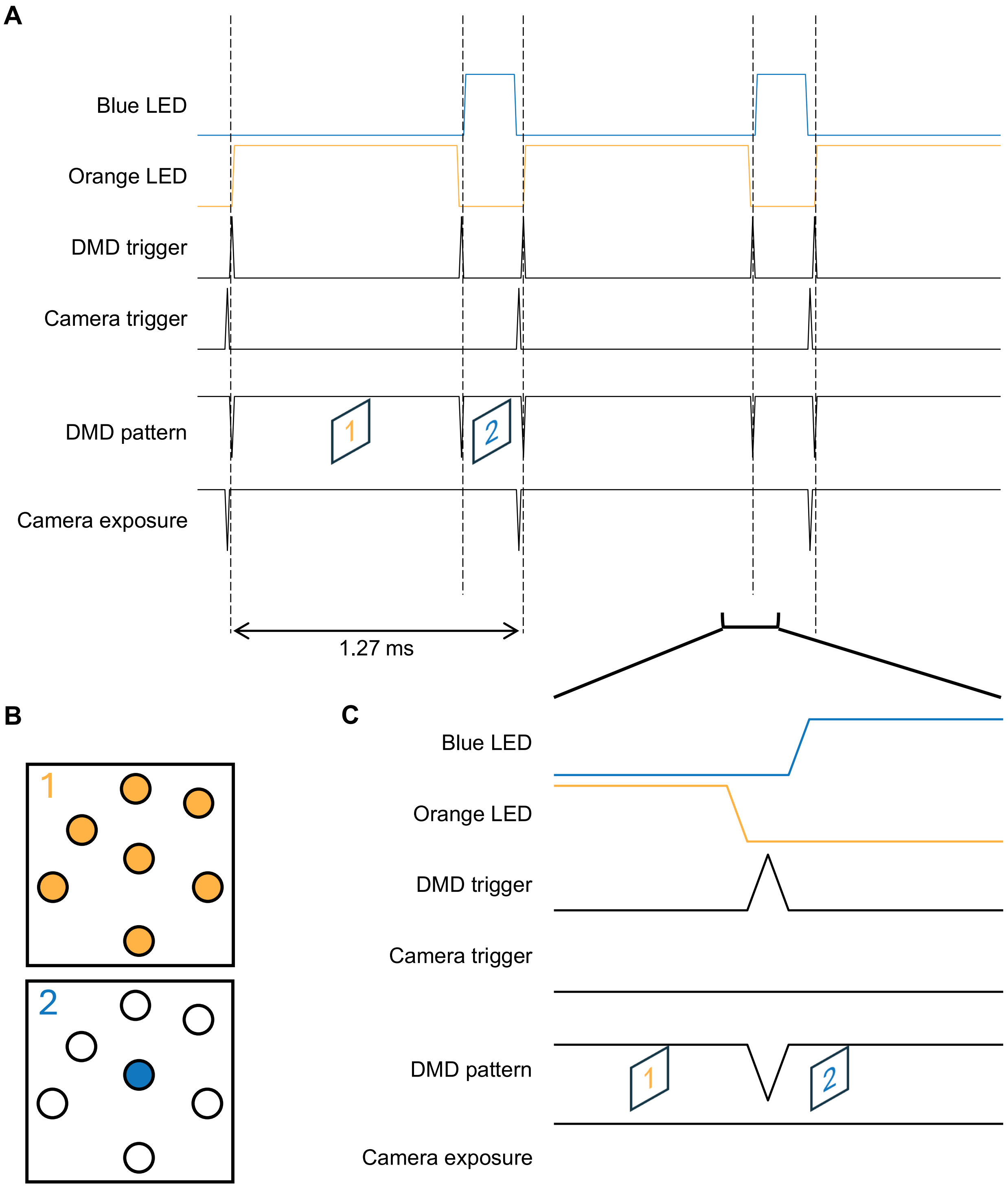


**Supplementary Figure 3. Simultaneous patterned illumination voltage imaging and targeted optogenetic stimulation using a single DMD.**

(A) Timing diagram for synchronous modulation of camera exposure, DMD patterns, and LED drive currents.

(B) DMD pattern schematics.

(C) Zoomed in view showing the transition from structured illumination for voltage imaging to targeted illumination for optogenetic stimulation.

Targeted illumination workflow:

1. Wide field fluorescence image
2. Hadamard digital confocal image
3. Image segmentation and DMD mask generation
4. DMD mask projection

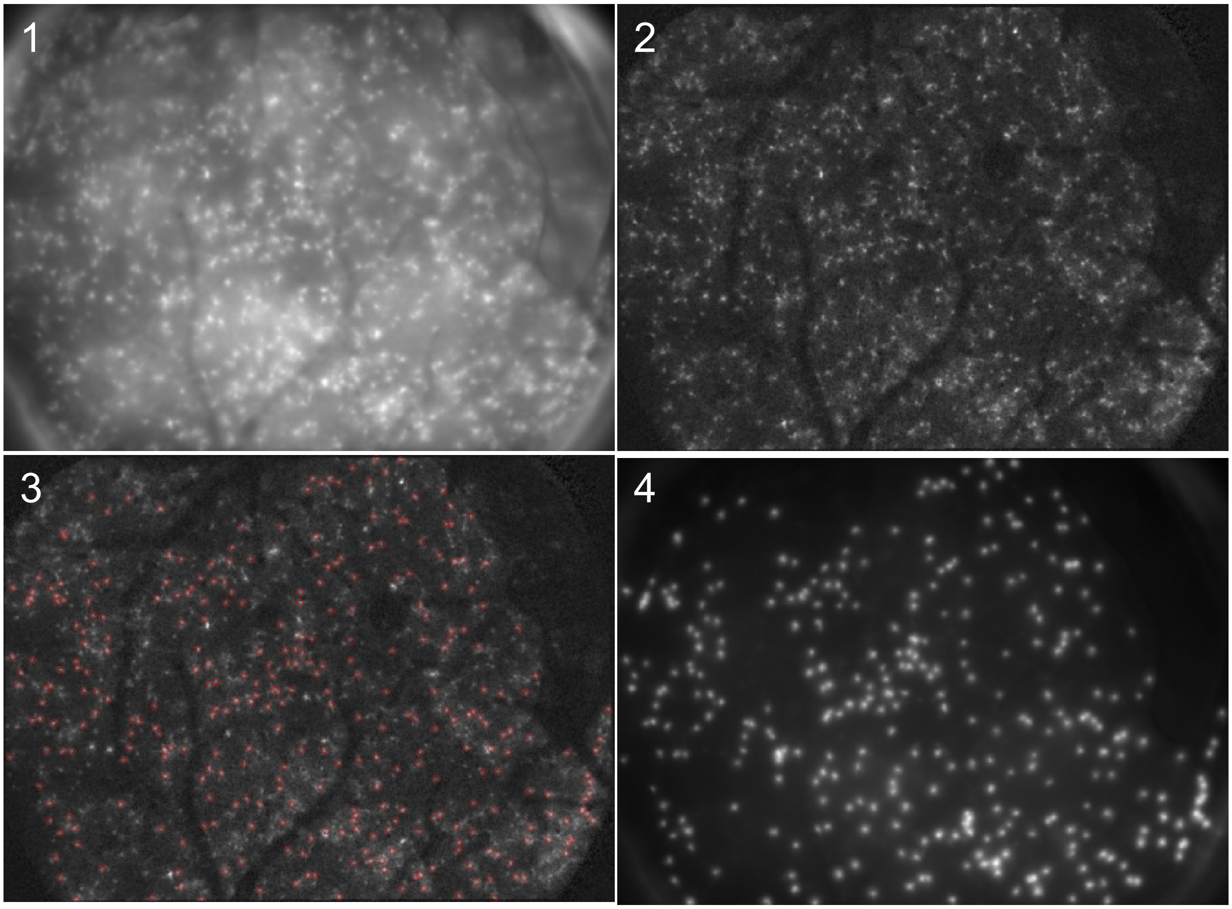


**Supplementary Figure 4. Targeted illumination workflow**

To generate soma-targeted illumination patterns for voltage imaging, we 1) found the FOV of interest under wide field illumination, 2) acquired a Hadamard digital confocal image, 3) segmented the confocal image and generate an illumination mask targeting neuronal somas, and 4) projected DMD mask to illuminate only the somas.

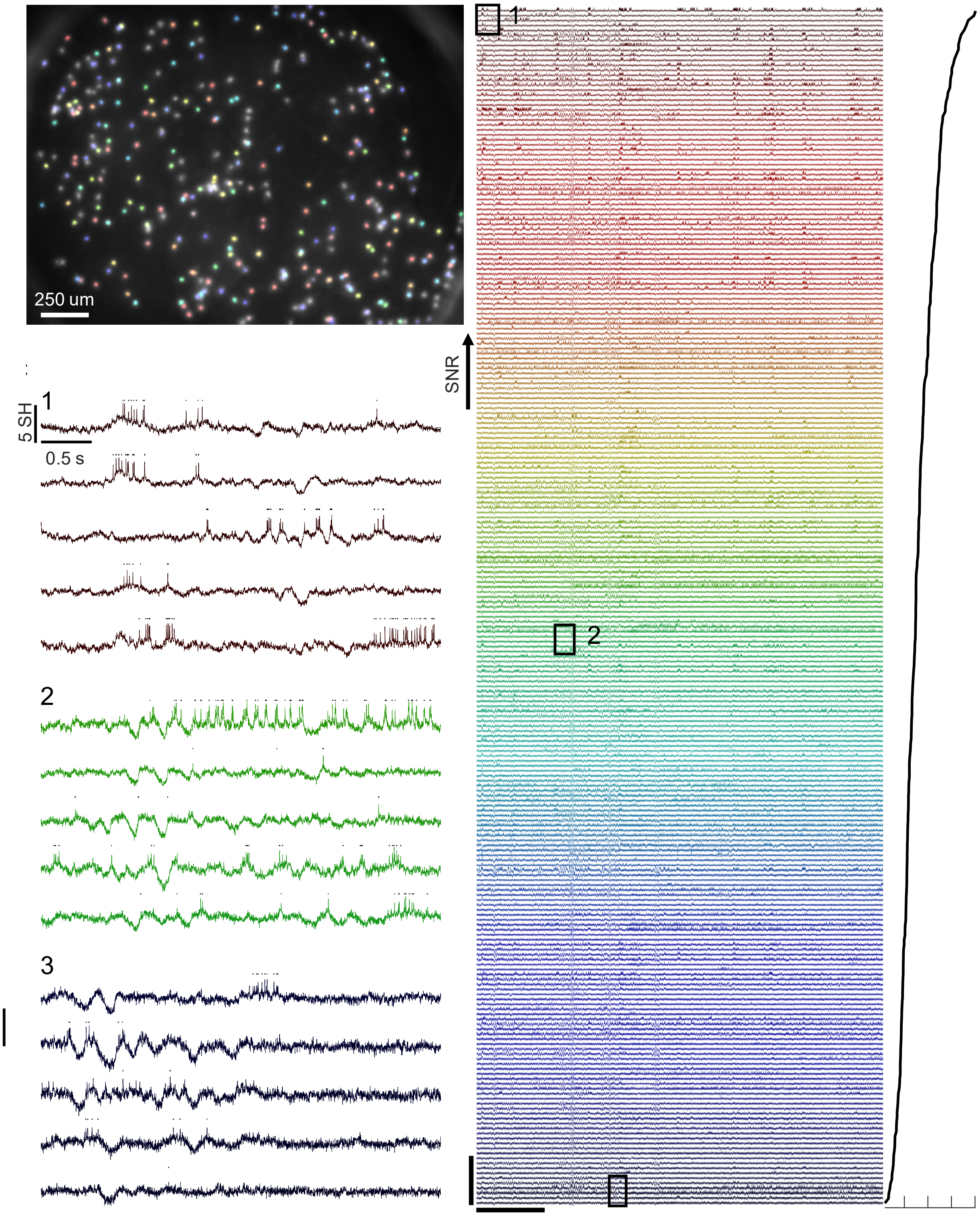


**Supplementary Figure 5. Example simultaneously recorded fluorescence traces.**

(A) Example FOV. The fluorescence traces of highlighted cells are shown in (B) with the same color scheme.

(B) Fluorescence traces sorted by SNR. Left: fluorescence traces, right: SNR.

(C) Zoomed-in view of regions highlighted by numbered boxes in (B). Black dots mark detected spikes, including some false positives.

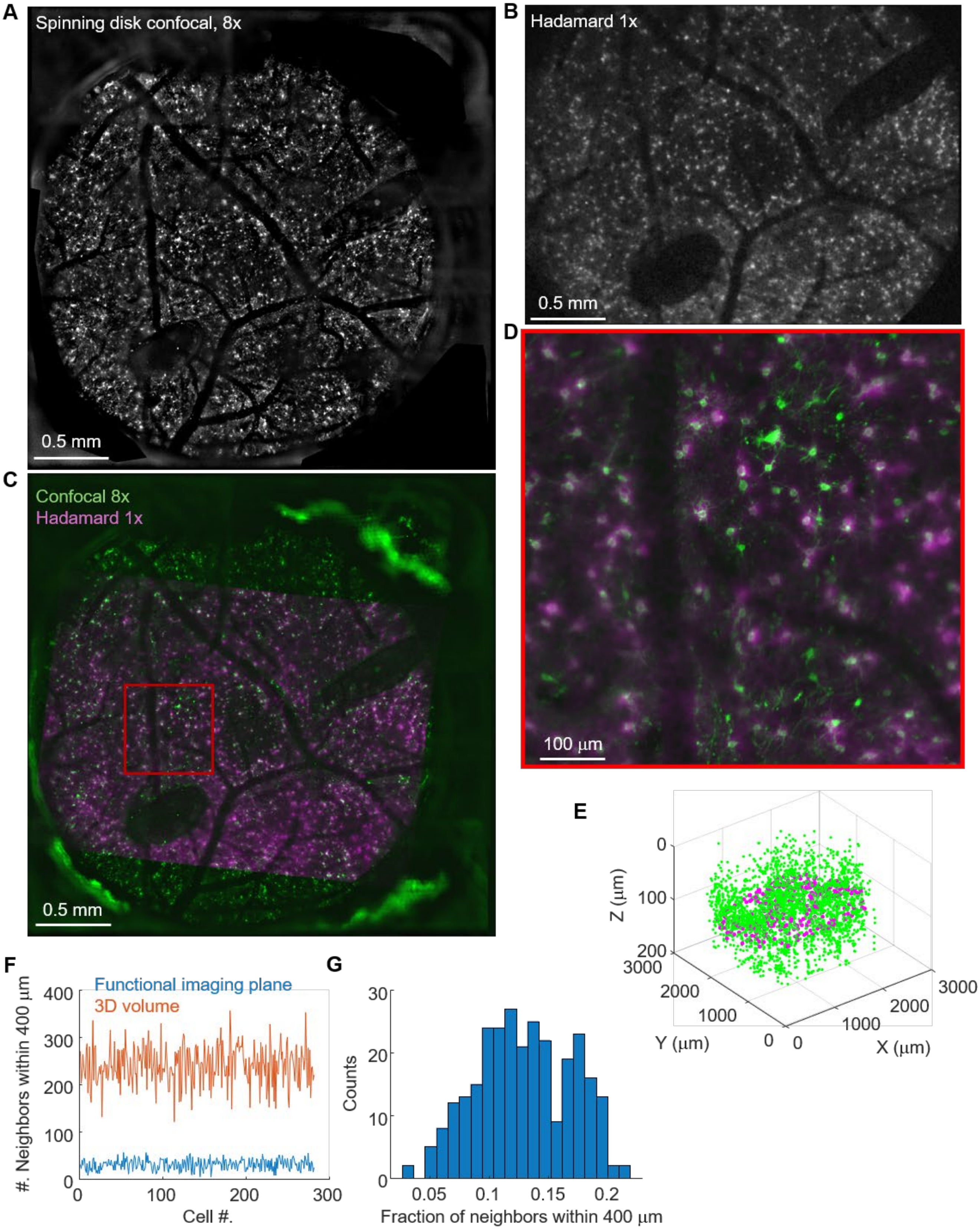
A
Spinning disk confocal, 8x
0.5 mm
B
Hadamard 1x
0.5 mm
C
Confocal 8x
Hadamard 1x
0.5 mm
D
100 μm
E
Z (μm)
0
100
200
3000
2000
1000
0
Y (μm)
0
1000
2000
3000
X (μm)
F
Functional imaging plane
3D volume
#. Neighbors within 400 μm
400
300
200
100
0
0
100
200
300
Cell #.
G
Counts
30
20
10
0
0.05
0.1
0.15
0.2
Fraction of neighbors within 400 μm

**Supplementary Figure 6. High-resolution volumetric reconstruction of L1**

(A) Maximum intensity projection of volumetric image of L1 acquired using spinning disk confocal microscope in the barrel cortex L1 of NDNF-Cre mouse.

(B) Hadamard digital confocal image of L1, same mouse as in A.

(C) Registration of the Hadamard digital confocal image onto the volumetric image. Green: Maximum projection of volumetric image, magenta: registered Hadamard digital confocal image.

(D) Zoomed-in view of the red box in C.

(E) Registration of the voltage imaging cell coordinates onto the volumetric reconstruction. The volumetric image was segmented using Cellpose 3.0. Green: volumetric reconstruction coordinates, magenta: voltage imaging cell coordinates.

(F) Neighbor density comparison between the voltage imaging plane and entire 3D volume of L1 for one mouse.

(G) Fraction of neighbors sampled by the voltage imaging plane for one mouse.

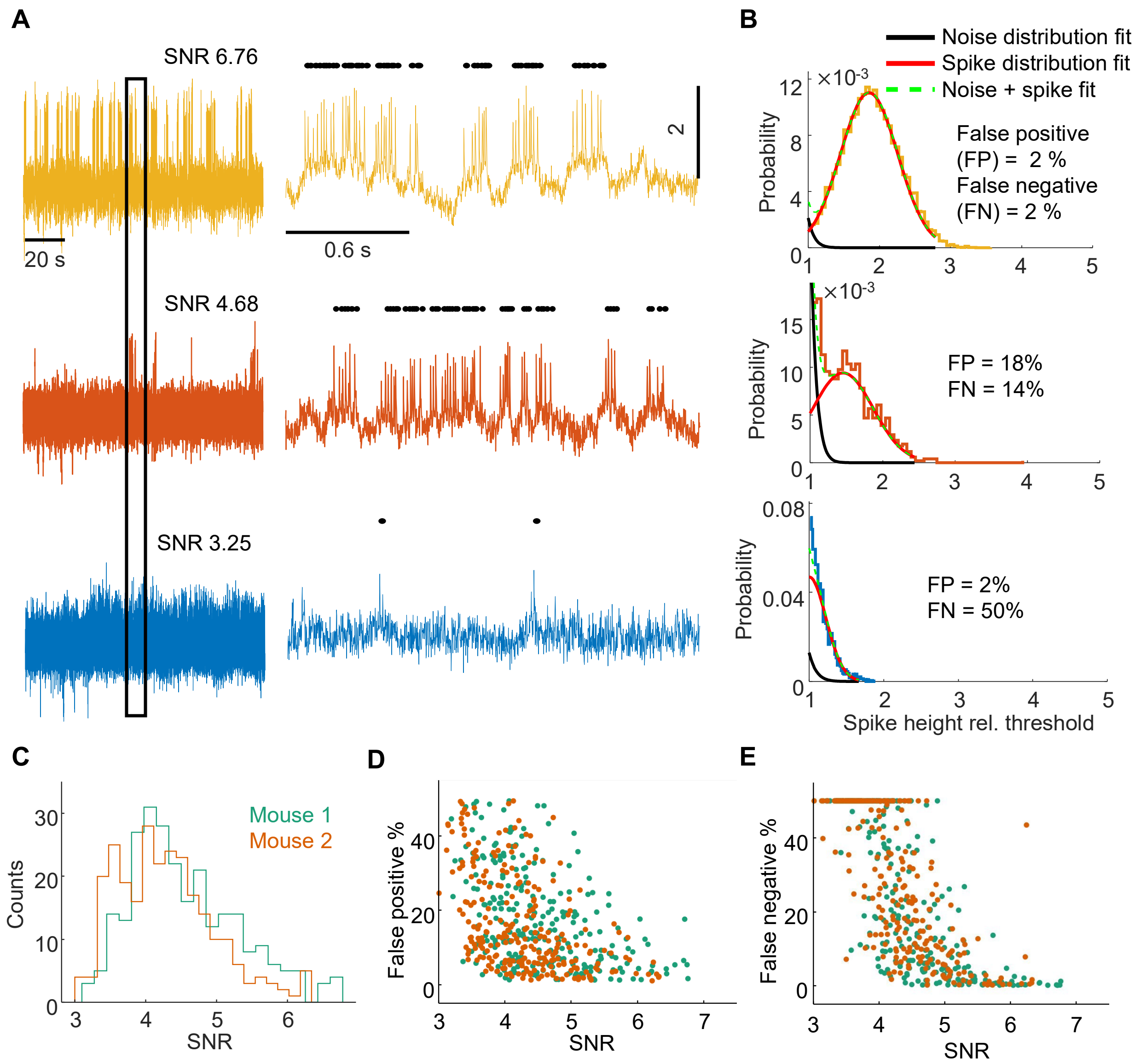


**Supplementary Figure 7. Relationship between SNR and spike detection accuracy.**

(A) Example fluorescence traces with high (top), medium (middle) and low (bottom) SNR. Left: full trace, right: zoomed-in trace.

(B) Gaussian mixture model fitting results for the three cells shown in (A). Black line: truncated Gaussian fit for supra-threshold noise. Red line: truncated Gaussian fit for spike height. Green dashed line: mixture of these two Gaussian distributions.

(C) SNR distributions of typical recordings from two mice, n = 520 cells. Green: mouse 1, orange: mouse 2.

(D) Comparison between SNR and false positive percentages, n = 520 cells, N = 2 mice. Green dots: mouse 1, orange dots: mouse 2.

(E) Comparison between SNR and false negative percentages, n = 520 cells, N = 2 mice. Green dots: mouse 1, orange dots: mouse 2.

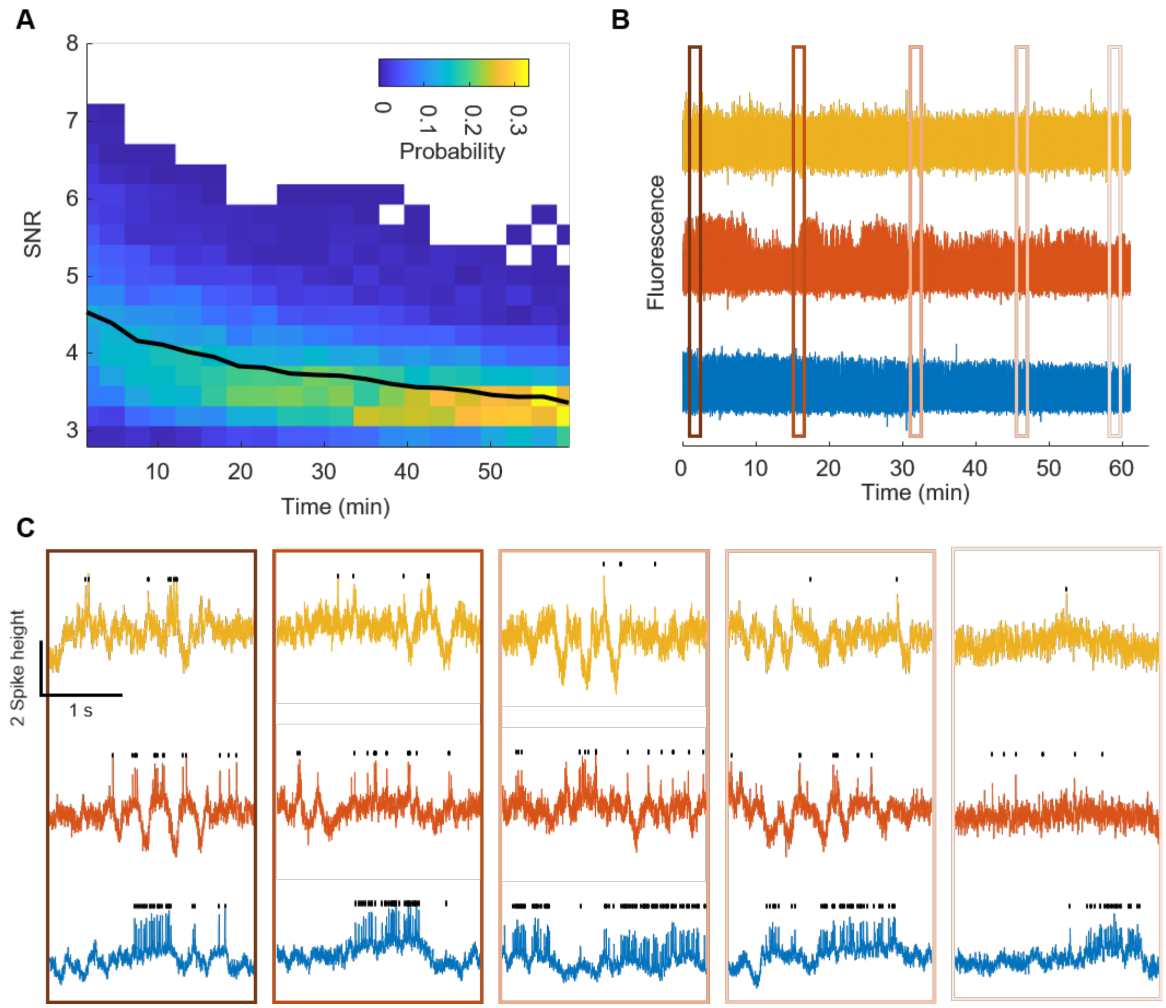


**Supplementary Figure 8. SNR decay during 1 hour of continuous imaging *in vivo*.**

(A) Distribution of single-cell SNR values (spike height:shot noise) during 1 hour of continuous illumination, n = 520 cells, N = 2 mice. Recording was divided into 3 min chunks, on which SNR was calculated. Black line: mean SNR across all cells.

(B) Three example fluorescence traces with low (yellow), medium (orange) and high (blue) SNR.

(C) Zoomed-in views of (B)

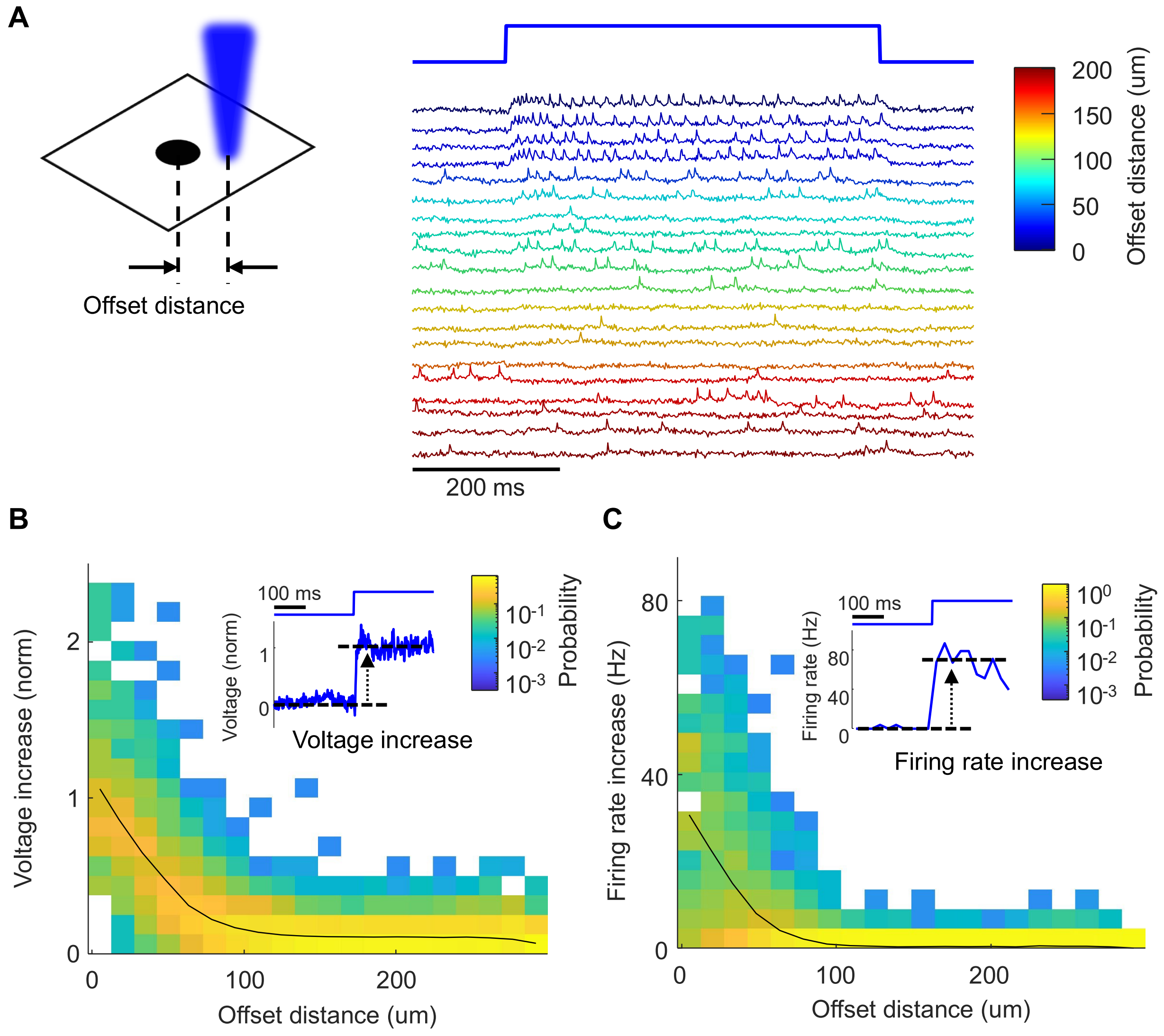


**Supplementary Figure 9. Characterizing off-target effects of optogenetic stimulation.**

(A) Example of a cell responding to off-target stimulation with increasing offset distance.

(B) Histogram of off-target stimulus-evoked subthreshold voltage increase as a function of offset distance. Each column was normalized by the number of cells in the column. Black line: average voltage increase. Inset: example trial-average voltage response of a cell to stimuli less than 10 $\mu$m away.

(C) Histogram of off-target stimulus-evoked firing rate increase as a function of offset distance. Each column was normalized by the number of cells in the column. Black line: average firing rate increase. Inset: example trial-average firing rate response of a cell to stimuli less than 10 $\mu$m away.

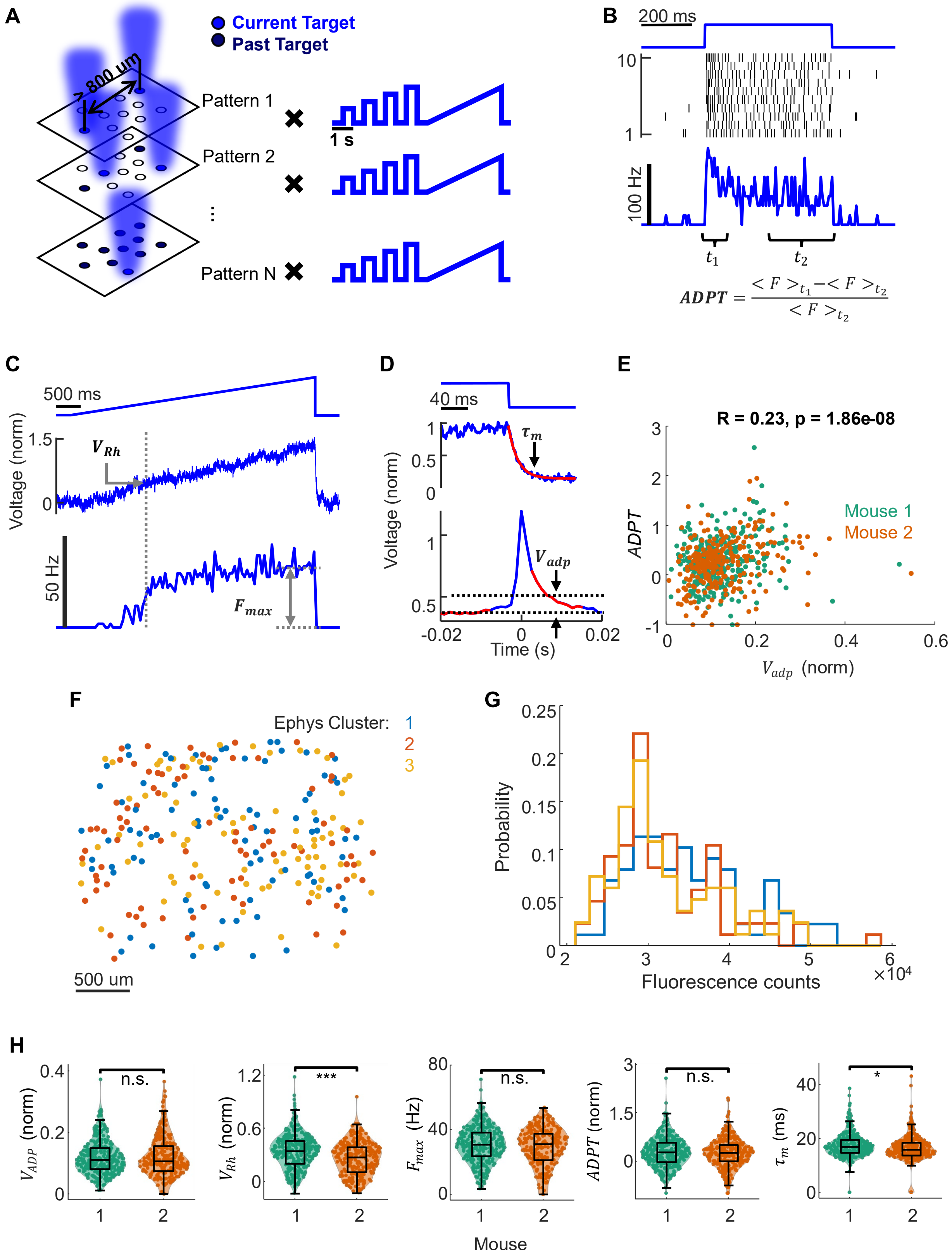


**Supplementary Figure 10. Mapping intrinsic excitability.**

(A) Intrinsic excitability mapping protocol. Sparse subsets of cells were stimulated with a step and ramp pattern. No two simultaneously stimulated cells were closer than 800 $\mu$m.

(B) Example response to 500-ms step stimulation, showing spike raster, trial-average firing rate, and definition of firing rate adaptation.

(C) Example response to 5-s ramp stimulation, showing trial-average subthreshold voltage and trial-average firing rate.

(D) Top: example trial-average voltage response to illumination turn-off, and exponential fit (red). Bottom: example average waveform of spikes evoked by blue light. The segments used to calculate $V_{adp}$ are highlighted in red.

(E) $ADPT$ and $V_{adp}$ shows correlation consistent across two mice (mouse 1 in green, mouse 2 in orange). Pearson correlation R = 0.23, p = 1.86E-8, n = 564 cells, 2 mice.

(F) Cells in each of the three intrinsic property bins (see text) are distributed across the FOV.

(G) Cells in each of the three intrinsic property bins have similar brightness distributions.

(H) Comparison of intrinsic properties of L1 interneurons between two mice, n = 320 for each mouse. Double sided Wilcoxon rank sum test was used. *, **, ***, indicate $0.01 < p \leq 0.05$, $0.001 < p \leq 0.01$, and $p \leq 0.001$.

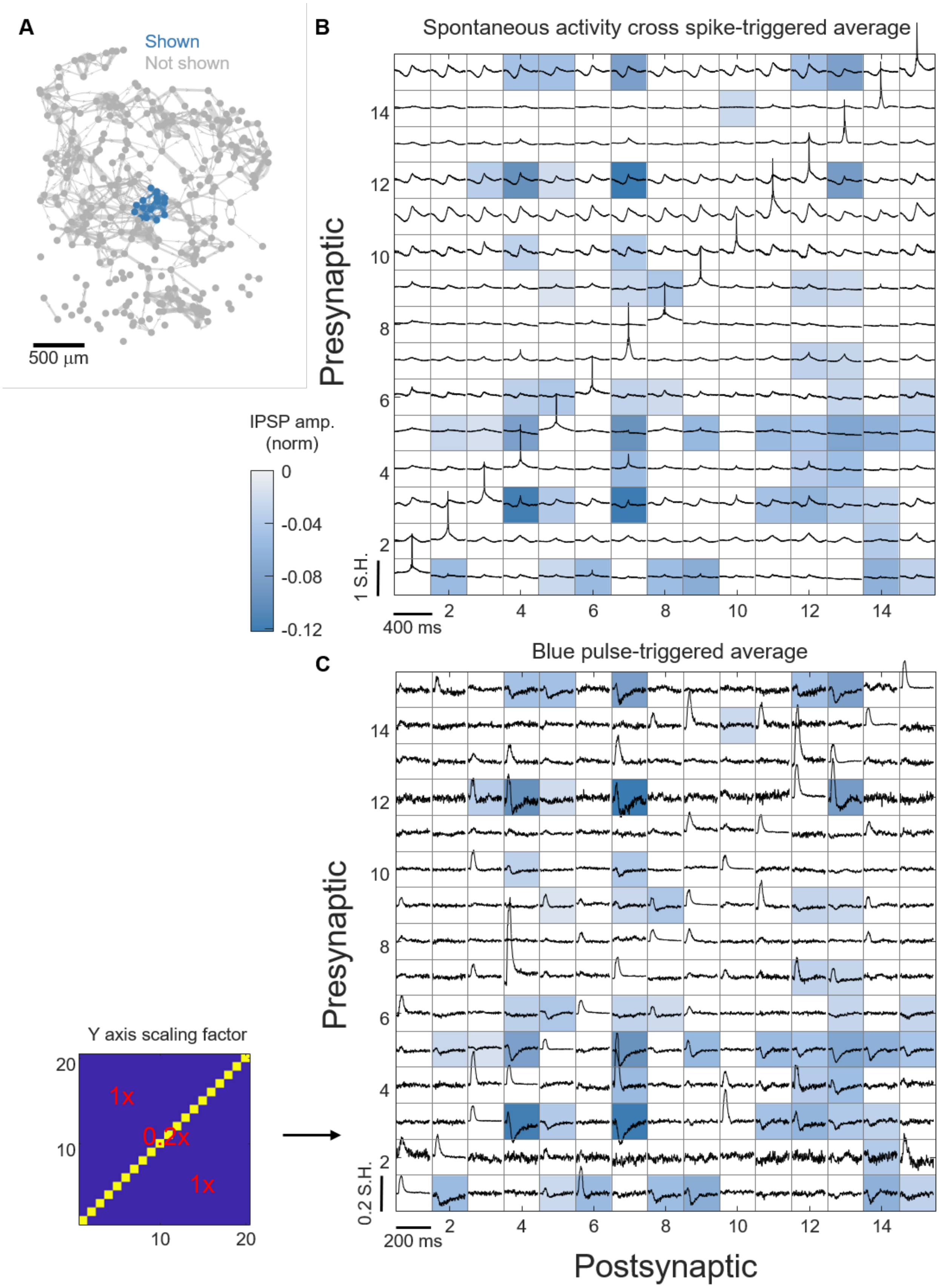
A
Shown
Not shown
500 µm
B
Spontaneous activity cross spike-triggered average
Presynaptic
IPSP amp.
(norm)
0
-0.04
-0.08
-0.12
1 S.H.
400 ms
C
Blue pulse-triggered average
Presynaptic
Y axis scaling factor
1x
0.2x
1x
0.2 S.H.
200 ms
Postsynaptic

**Supplementary Figure 11. Comparison between spontaneous activity cross STA voltage and optogenetically measured synaptic connectivity**

(A) Synaptic connectivity matrix as directed graph. Dots mark cell positions. Lines mark statistically significant synaptic connections. Line thickness indicates connection strength. The sub-matrix shown in B and C is highlighted in blue.

(B) Cross STA voltage waveforms of the 225 pairs (15 cells x 15 cells) indicated in (A), calculated during periods of spontaneous activity. Elements along the diagonal indicate self-STA. Blue shading indicates the strength of the optogenetically measured IPSP for the same cell pairs. Connectivity (i.e. IPSP strength) is not apparent from the cross correlation between presynaptic spike train and postsynaptic voltage.

(C) Optogenetically evoked IPSP waveforms for the 225 pairs in (A) and (B). Blue shading indicates the strength of the optogenetically measured IPSPs, determined from this dataset. The diagonal traces are scaled by 0.2 as indicated by the inset on the left.

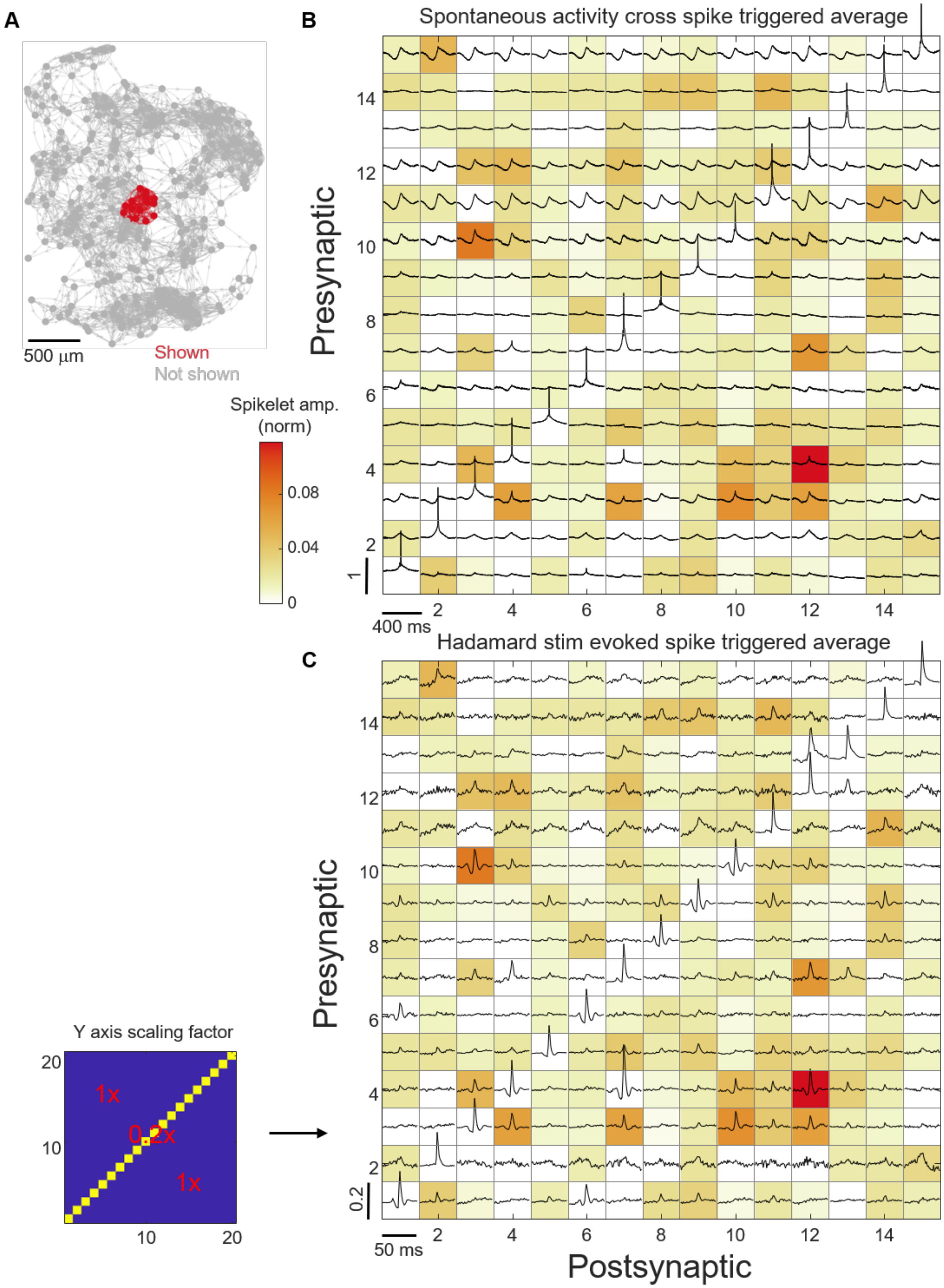

A
500 µm
Shown
Not shown
B
Spontaneous activity cross spike triggered average
Presynaptic
Spikelet amp. (norm)
0.08
0.04
0
400 ms
C
Hadamard stim evoked spike triggered average
Postsynaptic
50 ms
Y axis scaling factor
1x
0.2x
1x

**Supplementary Figure 12. Comparison between spontaneous activity cross STA voltage and optogenetically measured gap junction connectivity**

(A) Gap junction connectivity matrix as a directed graph. Dots mark cell positions. Lines mark statistically significant gap junction connections. Line thickness indicates connection strength. The sub-matrix shown in B and C is highlighted in red.

(B) Cross STA voltage of 225 pairs (15 cells x 15 cells) overlaid on the gap junction connectivity matrix. These waveforms are the same as in Supplementary Figure 11 B. Spikelets are visible in some gap junction-connected pairs.

(C) Optogenetically evoked spikelet waveforms for the 225 pairs in (A) and (B). Shading indicates the strength of the optogenetically determined spikelets, determined from this dataset. The diagonal traces are scaled by 0.2 as indicated by the inset on the left.

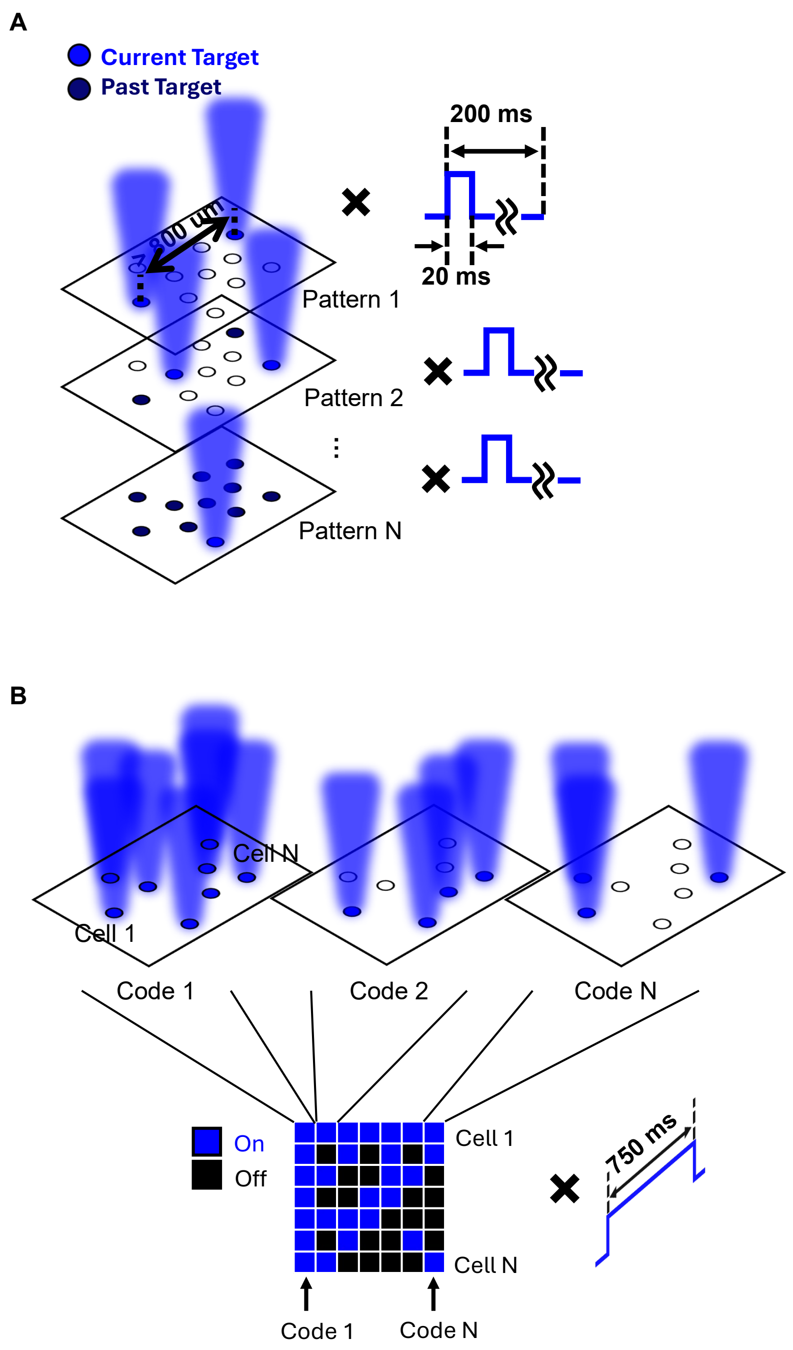


**Supplementary Figure 13. Synaptic and gap junction connectivity mapping protocols.**

(A) Synaptic connectivity mapping protocol. Simultaneously stimulated cells were never separated by < 800 µm.

(B) Gap junctional connectivity mapping protocol. Cells were stimulated with 750 ms pulses in a pseudo-random pattern, 50% duty cycle.

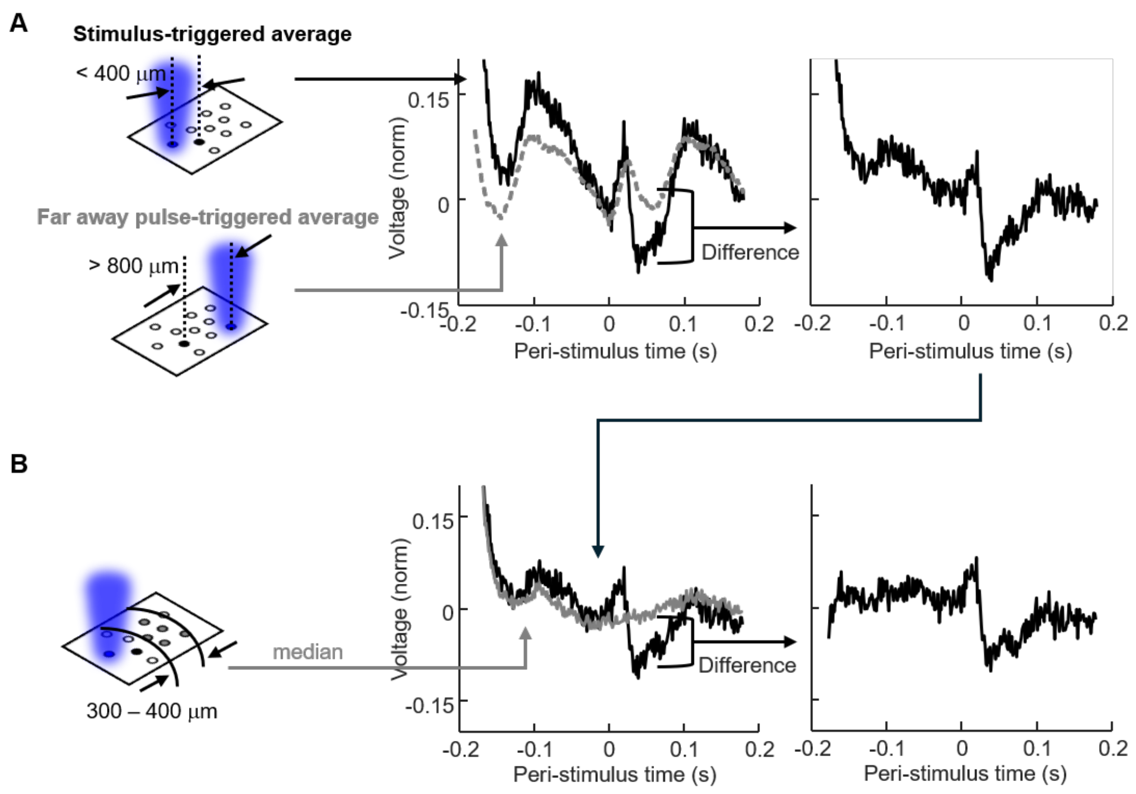


**Supplementary Figure 14. Computational corrections for stimulus artifacts during synaptic connectivity mapping.**

(A) Correction for blue light induced stimulus artifacts. Blue light induced stereotypical depolarization that propagated in the posterior-to-anterior direction, presumably originating from the visual cortex. We estimated this artifact for each postsynaptic cell *j* by sampling trials with blue pulses more than 800 μm from cell *j*, then calculating the trial average response (gray trace in the middle panel). We then removed this artifact by subtracting it from the stimulus triggered average for stimuli targeted to cell *i* < 400 μm from cell *j* (right panel).

(B) Correction for correlated spontaneous dynamics. For each presynaptic cell *i*, the IPSPs of its postsynaptic cells share correlated dynamics due to other shared synaptic inputs. We estimated these dynamics as the median stimulus-triggered average waveform of cells between 300 and 400 μm from cell *i* (gray trace in the middle panel). Since most cells at this distance were not connected to cell *i*, the median captured the effects of correlated dynamics at the stimulus times of cell *i*. We then subtract this control waveform from the IPSPs of all cells postsynaptic to cell *i*.

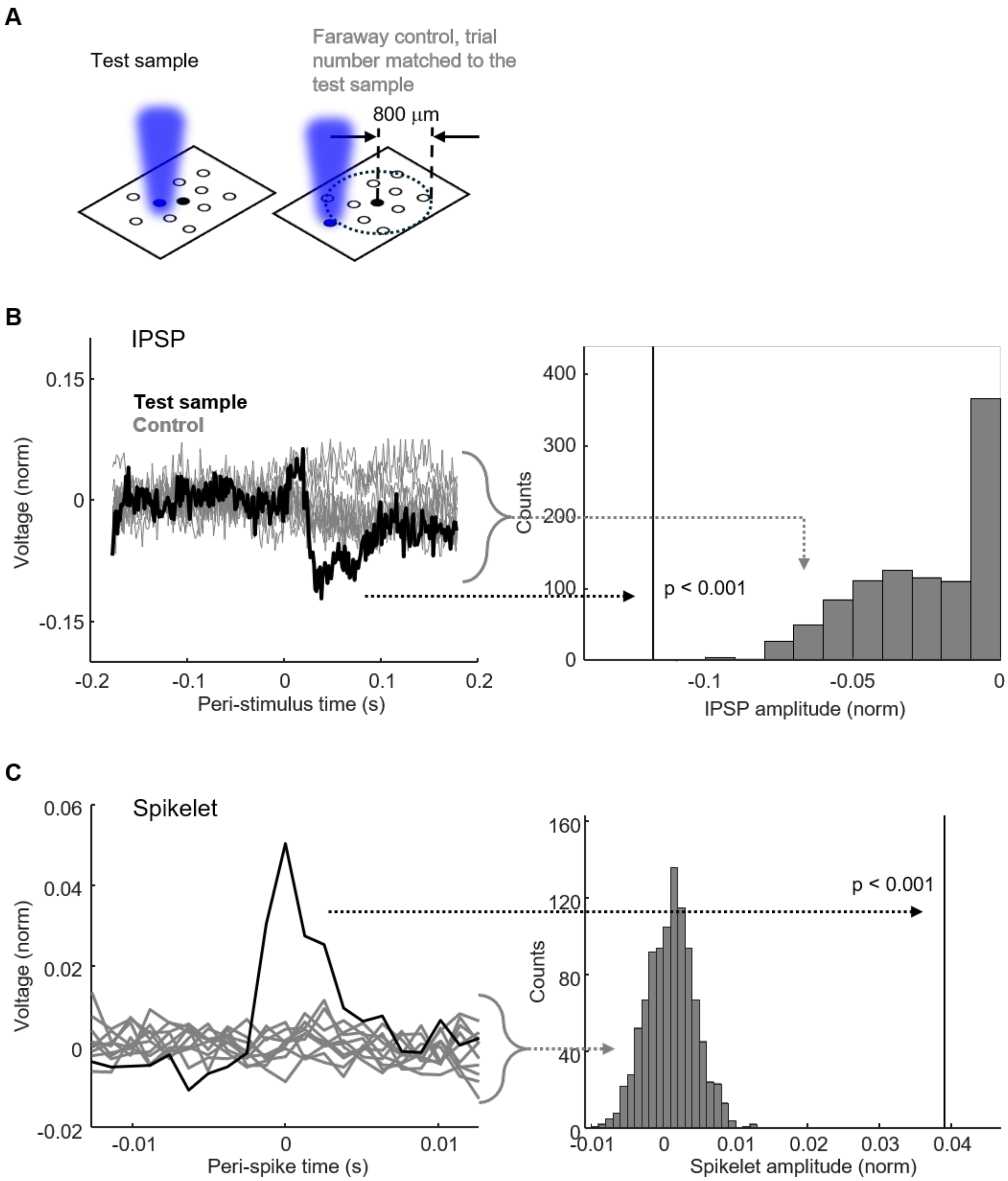


**Supplementary Figure 15. Null distributions for IPSP and spikelet amplitudes.**

(A) Sampling procedure to generate null distributions for IPSP and spikelet amplitudes.

(B) Example of a statistically significant IPSP (black trace) and the null IPSP traces (gray traces).

(C) Example of a statistically significant spikelet (black trace) and the null spikelet traces (gray traces).

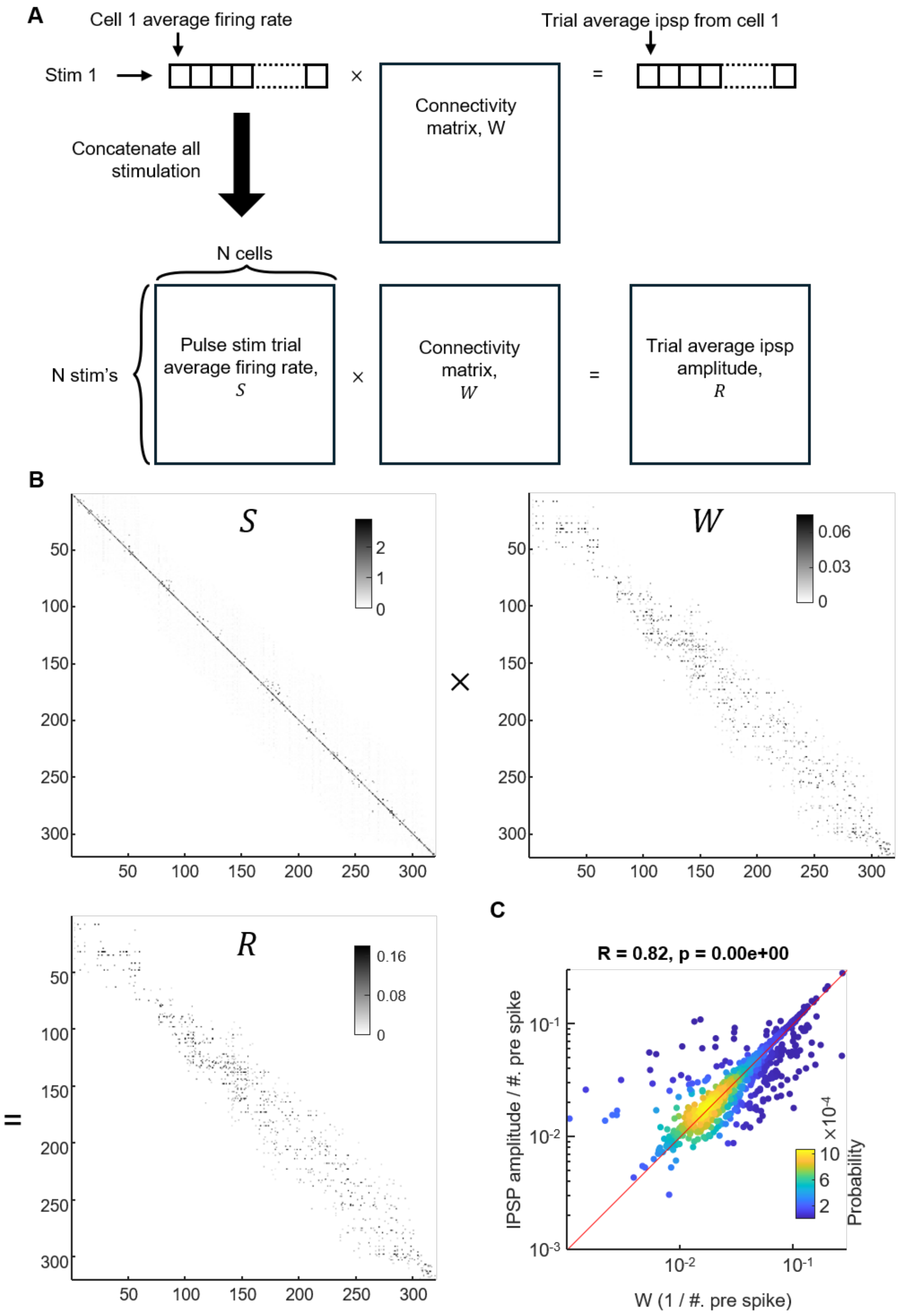


**Supplementary Figure 16. Effect of blue light scatter on synaptic connectivity mapping.**

(A) We included the possibility that a stimulus targeted to cell *i* could evoke spikes in cell *k* which would contribute to the IPSP in cell *j*. Since we measured the voltage of all cells during the optical stimuli, these off-target effects could be incorporated directly. In the presence of off-target stimulation, the evoked spiking matrix *S* (stimulus to cell *i* evoking spikes in cell *k*) had off-diagonal elements. The IPSP matrix, *R*, was a matrix product of the firing rate matrix *S* and an unknown connectivity matrix, *W*.

(B) Example solution of the unknown connectivity matrix *W* by lasso regression.

(C) Comparison between the connectivity matrix calculated including off-target effects and the IPSP matrix calculated assuming only on-target stimulation. The two matrices are highly correlated, with a Pearson correlation coefficient of 0.82, $p < 2E\text{-}308$, $n = 1755$ pairs from 2 mice, and less than 20% of statistically significant IPSPs differ from the connectivity matrix by more than 20%. Red line has slope of 1.

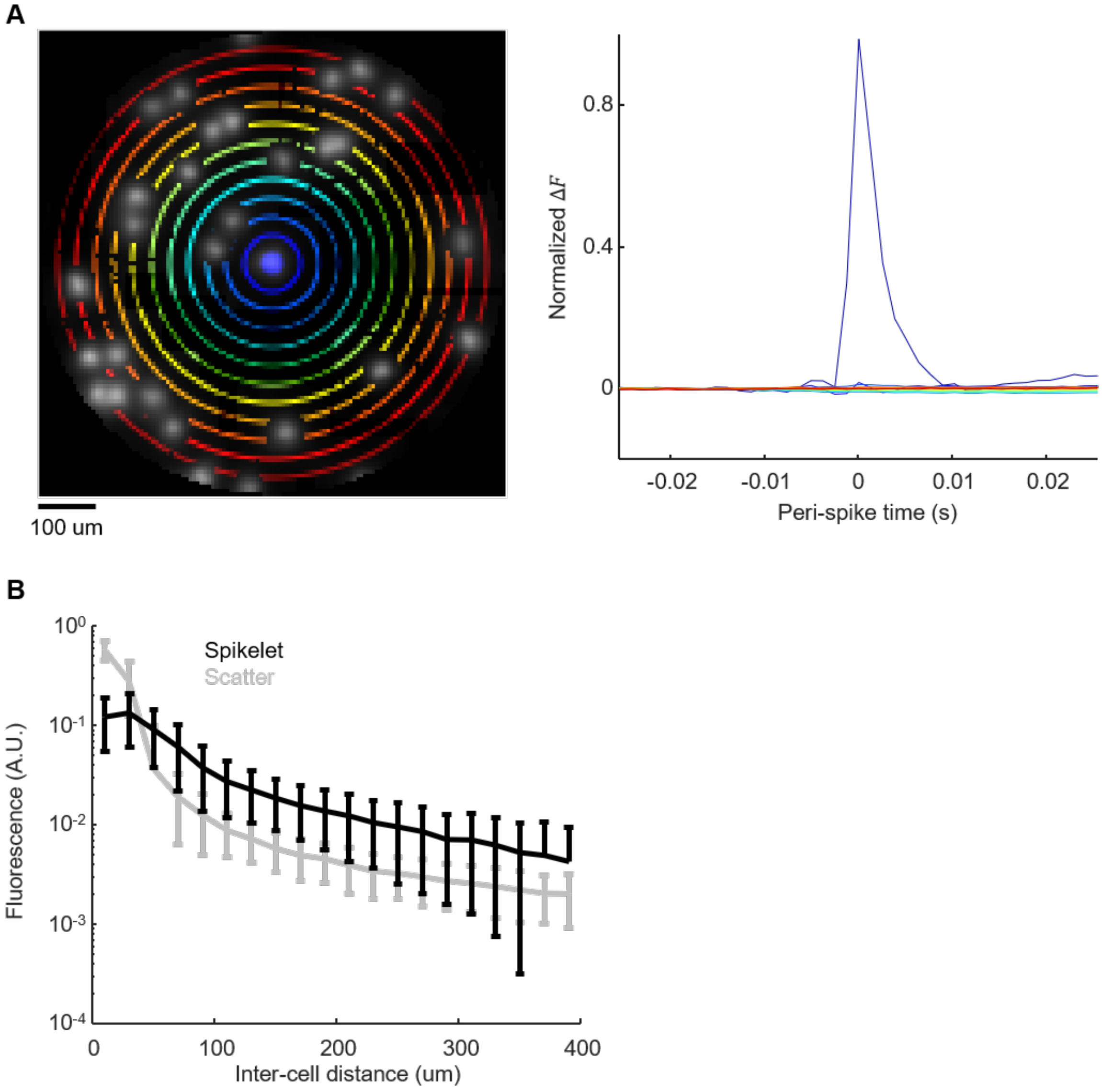


**Supplementary Figure 17. Quantifying the effect of scattered fluorescence.**

(A) Example of apparent spikelet due to fluorescence scatter. Left: FOV used to calculate the spikelet due to fluorescence scatter from the central cell. Regions containing other cells were excluded from the analysis. The cross STA fluorescence, triggered by spikes of the central cell, was calculated on colored pixels. Right: cross STA fluorescence for the colored pixels shown on the left.

(B) Comparison between gap junction-mediated spikelet amplitude and scattered fluorescence as a function of distance from the triggering cell. Cell pairs (real spikelet) or cell-pixel pairs (scatter) were divided into 20 distance bins (bin size = 40 μm). For each bin, we calculated the average (line) and std (error bar). Black: real spikelets, gray: light scatter.

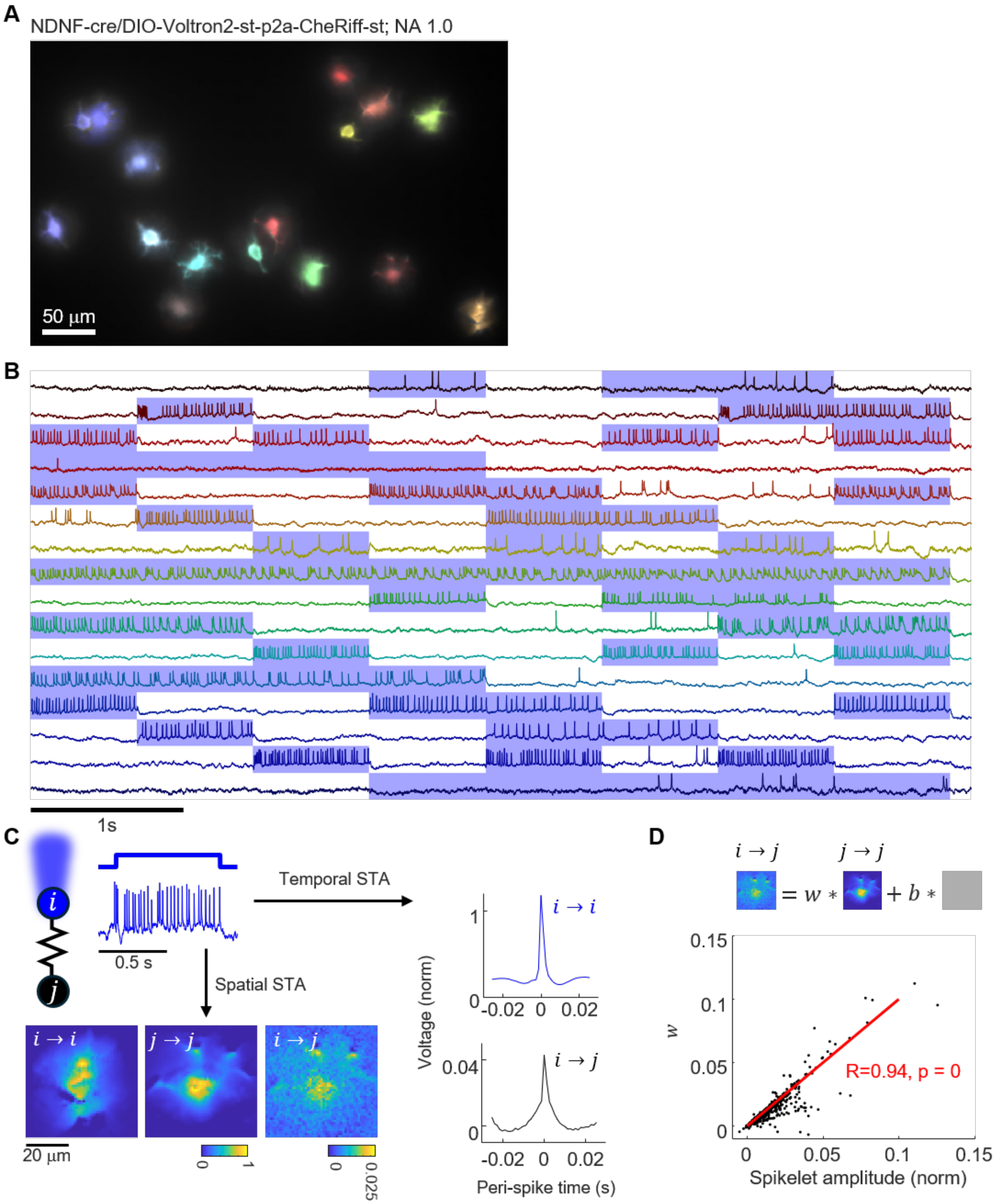


**Supplementary Figure 18. Validation of gap junction mapping protocol under high magnification imaging.** We used the spatial footprints of putative spikelets to confirm that these events followed the membrane of the post-junctional cell.

(A) FOV of a high-magnification gap junctional connectivity mapping experiment. Here cells expressed the Voltron2 version of the Optopatch construct. Cells have been pseudo-colored to indicate corresponding fluorescence traces shown in B.

(B) Fluorescence traces of cells highlighted in A. Blue shading indicates cell-targeted stimulation epochs, which followed a pseudo-random pattern of 750 ms pulses, 50% duty cycle.

(C) For a pair of cells, we calculated the self STA of each to map its membrane footprint. We then calculated the pixel-wise cross STA of voltages in cell *j* triggered off spikes in cell *i*. We compared the footprint of the *i* → *j* cross STA to the footprint of the *j* → *j* self STA to determine if the cross event corresponded to changes in membrane potential of cell *j*.

(D) Correlation between spikelet amplitude and spikelet footprint weight. The footprint of spikelet for cell pair (i,j) is modeled as a weighted sum of the spike footprint of cell j and a flat background. The Pearson correlation coefficient was 0.94, $p < 2E\text{-}308$, $n = 720$ pairs, from 48 cells, indicating that the cross-STA primarily corresponded to voltage changes in downstream cells.

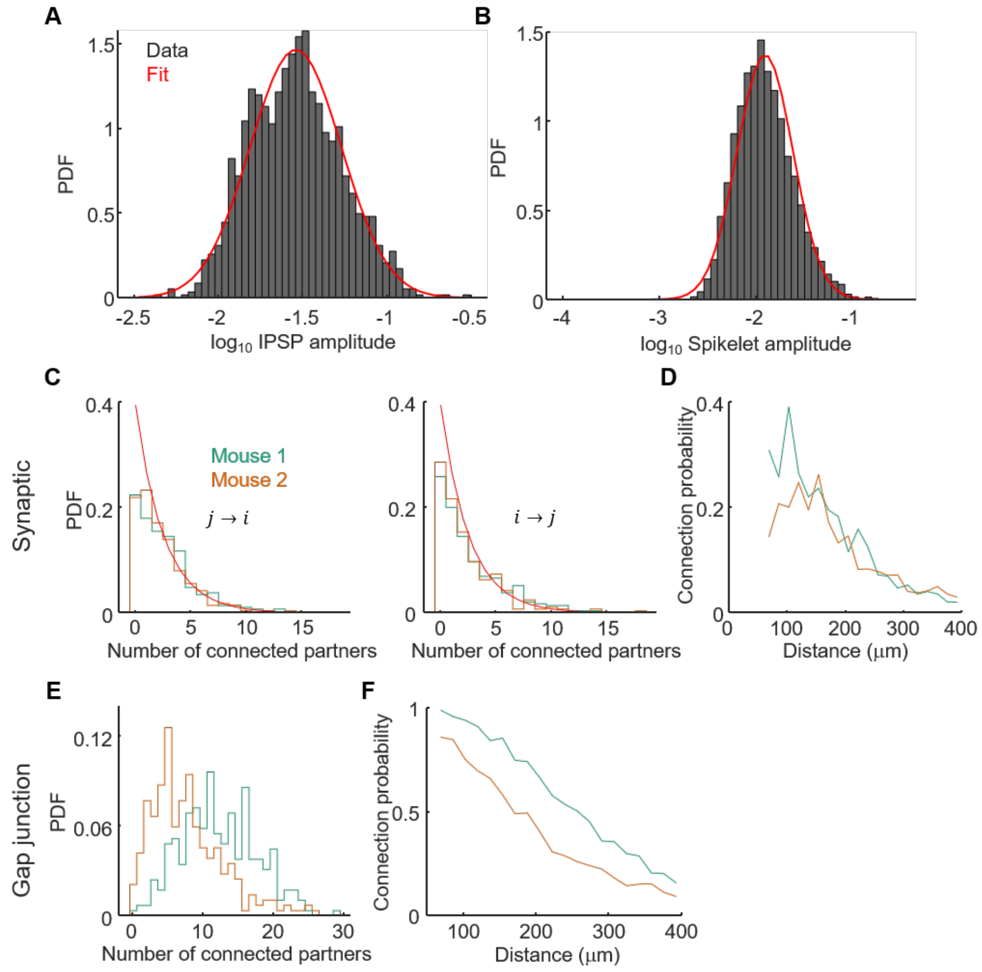


**Supplementary Figure 19. Distributions of synaptic and gap junctional connection strengths.**

(A) Log-normal distribution of IPSP amplitudes. Red line: Gaussian fit.

(B) Log-normal distribution of spikelet amplitudes. Red line: Gaussian fit.

(C) Degree distribution of synaptic connectivity. Left: degree distribution for $j \rightarrow i$ (incoming) connections, right: degree distribution for $i \rightarrow j$ (outgoing) connections. Red line: exponential fit. Green: mouse 1, orange: mouse 2.

(D) Synaptic connection probability as a function of distance. Green: mouse 1, orange: mouse 2.

(E) Degree distribution of gap junction connectivity. Green: mouse 1, orange: mouse 2.

(F) Gap junctional connection probability as a function of distance. Green: mouse 1, orange: mouse 2.

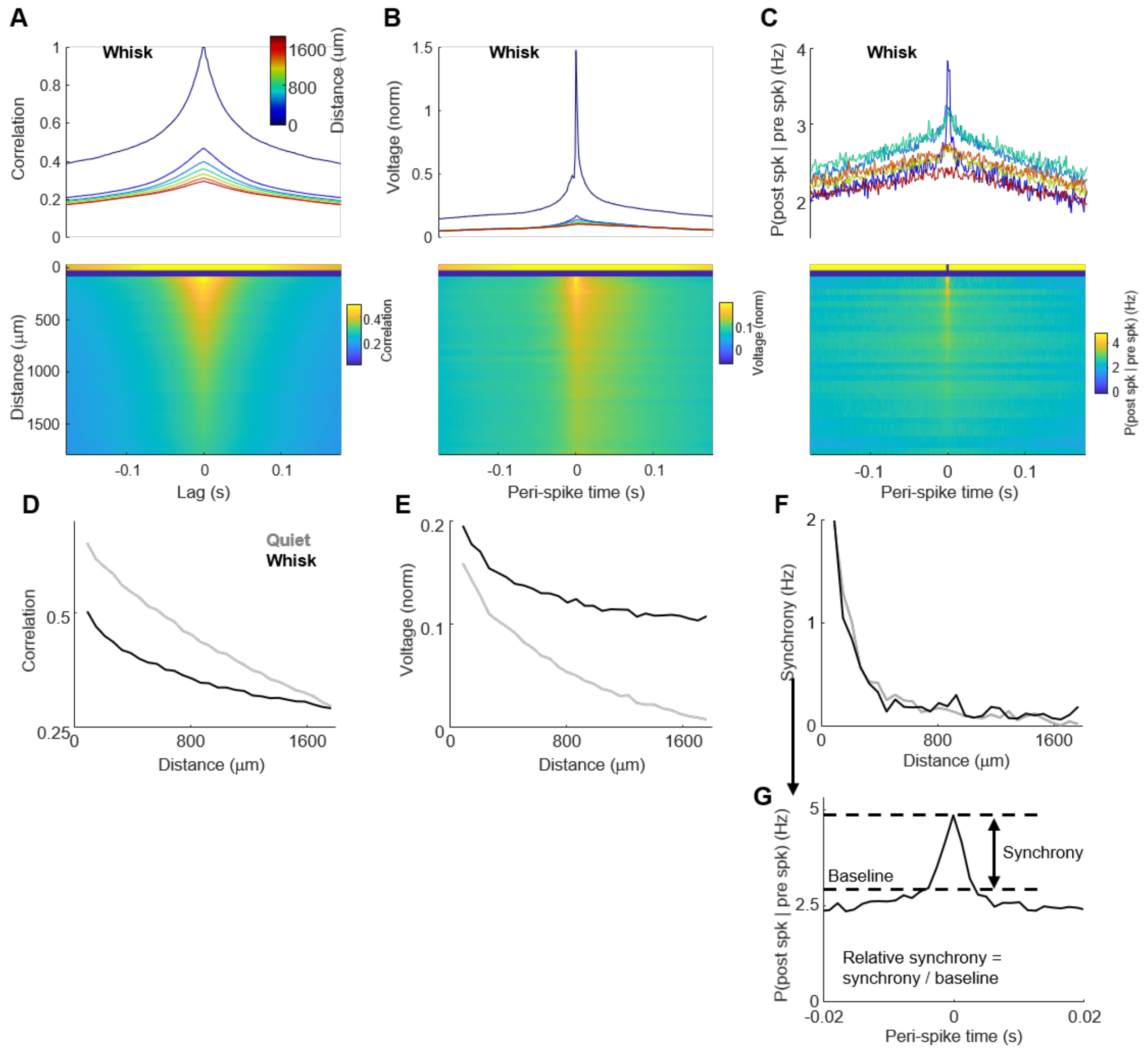


**Supplementary Figure 20. State-dependent correlational structure of subthreshold voltages and spike patterns.**

(A) Subthreshold voltage cross-correlation functions in the whisking state, resolved by cell-cell distance (see Main Text for quiet-state cross-correlations). We divided cell pairs into 30 distance bins (bin size = 60 µm) then computed the average cross-correlation function for cell pairs inside each bin. Cell pairs in the 0 – 60 µm bin were omitted due to fluorescence scatter. Top: seven examples of population average cross-correlation functions, bottom: all of the population average cross-correlation functions. We repeated the same distance binning and averaging in B and C.

(B) Spike-triggered average voltage waveform in the whisking state, resolved by cell-cell distance as in (A).

(C) Spike-rate cross correlogram in the whisking state, resolved by cell-cell distance as in (A).

(D) State-dependent equal-time ($\tau = 0$) cross-correlation vs. distance. Gray: quiet state, black: whisk-state.

(E) State-dependent equal-time ($\tau = 0$) cross-STA voltage vs. distance. Gray: quiet state, black: whisk-state.

(F) State-dependent spike synchrony vs. distance. Spike synchrony is defined in (G). Gray: quiet state, black: whisk-state.

(G) Definition of spike synchrony. For baseline, we calculated the average value of the spike cross-correlogram between $4\ \mathrm{ms} \le \tau \le 6\ \mathrm{ms}$ both before and after $\tau = 0$, then took the higher value.

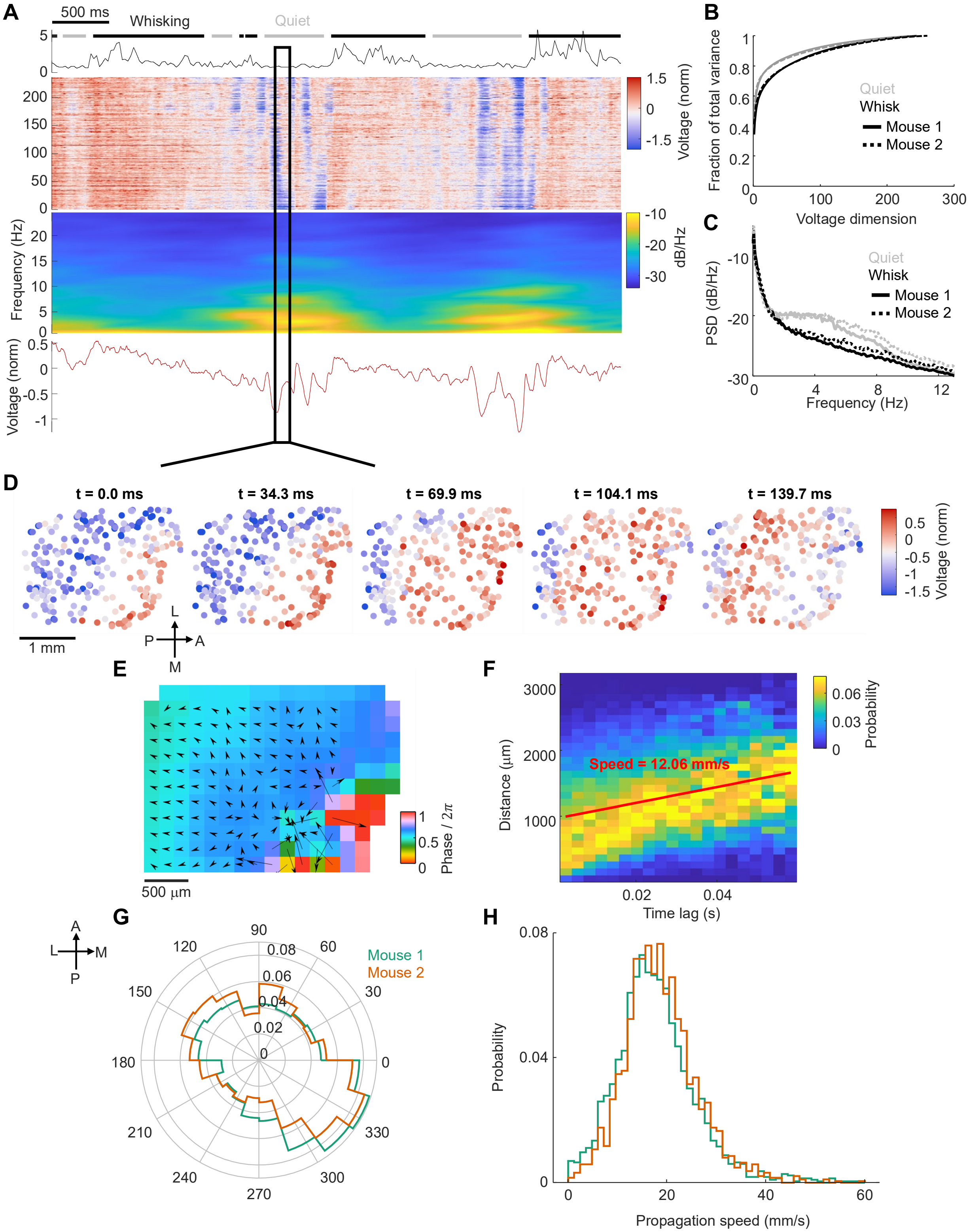


**Supplementary Figure 21. Subthreshold wave propagation during the quiet state.**

(A) Spontaneous subthreshold dynamics and its spectrogram. Top: whisking motion, gray and black bars mark state assignments. Middle: subthreshold traces of 247 cells sorted by the x-coordinate of the cell, and spectrogram computed by averaging the sliding power spectral density (PSD) across all cells. Bottom: average subthreshold voltage across all cells.

(B) Cumulative sum of the eigenvalues of the subthreshold voltage covariance matrix. Gray: quiet state, black: whisking state. Solid: mouse 1, dashed: mouse 2.

(C) State-dependent population-average subthreshold PSD. Gray: quiet state, black: whisking state. Solid: mouse 1, dashed: mouse 2.

(D) Snapshots of a subthreshold wave propagation event highlighted in A.

(E) Phase gradient of the propagation event in D.

(F) Distance-lag plot of the propagation event in D. For each event, we took snippets of subthreshold voltages of all cells then calculated the cross-correlation for all pairs. We then extracted the lag time of the peak of the cross-correlation. Red line: linear fit. Slope is the propagation speed, 12.06 mm/s.

(G) Histogram of the principal propagation direction for all wave propagation events during spontaneous activity, n = 3858 events, N = 2 mice. Green: mouse 1, orange: mouse 2.

(H) Histogram of propagation speeds for all wave propagation events during spontaneous activity, n = 3858 events, N = 2 mice. Green: mouse 1, orange: mouse 2.

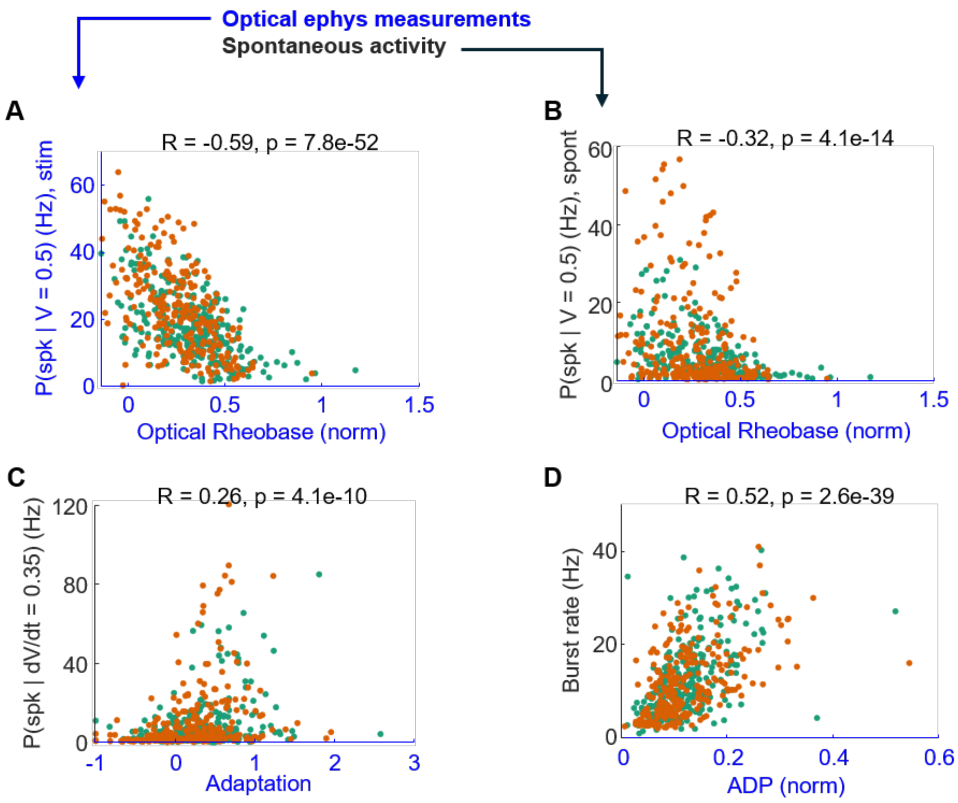


**Supplementary Figure 22. Correlation between optogenetically measured intrinsic properties and the subthreshold-to-spiking transforms during spontaneous activity.**

(A) Correlation between optogenetically evoked conditional spike rate at $V = 0.5$ and optical rheobase. R = -0.59 (Pearson correlation), p = 7.8e-52, n = 543, N = 2 mice. Green: mouse 1, orange: mouse 2.

(B) Correlation between spontaneous activity conditional spike rate at $V = 0.5$ and optogenetically measured rheobase. R = -0.32 (Pearson correlation), p = 4.1e-14, n = 543, N = 2 mice. Green: mouse 1, orange: mouse 2. This plot shows that the optogenetically measured rheobase is related to the intrinsic excitability during spontaneous activity.

(C) Correlation between spontaneous activity conditional spontaneous spike rate at $dV/dt = 0.35$ and spike rate adaptation. R = 0.26 (Pearson correlation), p = 4.1e-10, n = 543, N = 2 mice. Green: mouse 1, orange: mouse 2. This plot shows that the spike-rate adaptation, which acts as a high-pass filter on inputs, correlates with the sensitivity to the rate of upstroke in the subthreshold voltage during spontaneous activity.

(D) Correlation between burst rate and after-depolarization. We defined the burst rate as the spike probability averaged from 2.5 - 15 ms lag in the spike autocorrelogram. R = 0.52 (Pearson correlation), p = 2.6e-39, n = 543, N = 2 mice. Green: mouse 1, orange: mouse 2.

This plot shows that the after-depolarization favors initiation of a second spike shortly after a first.

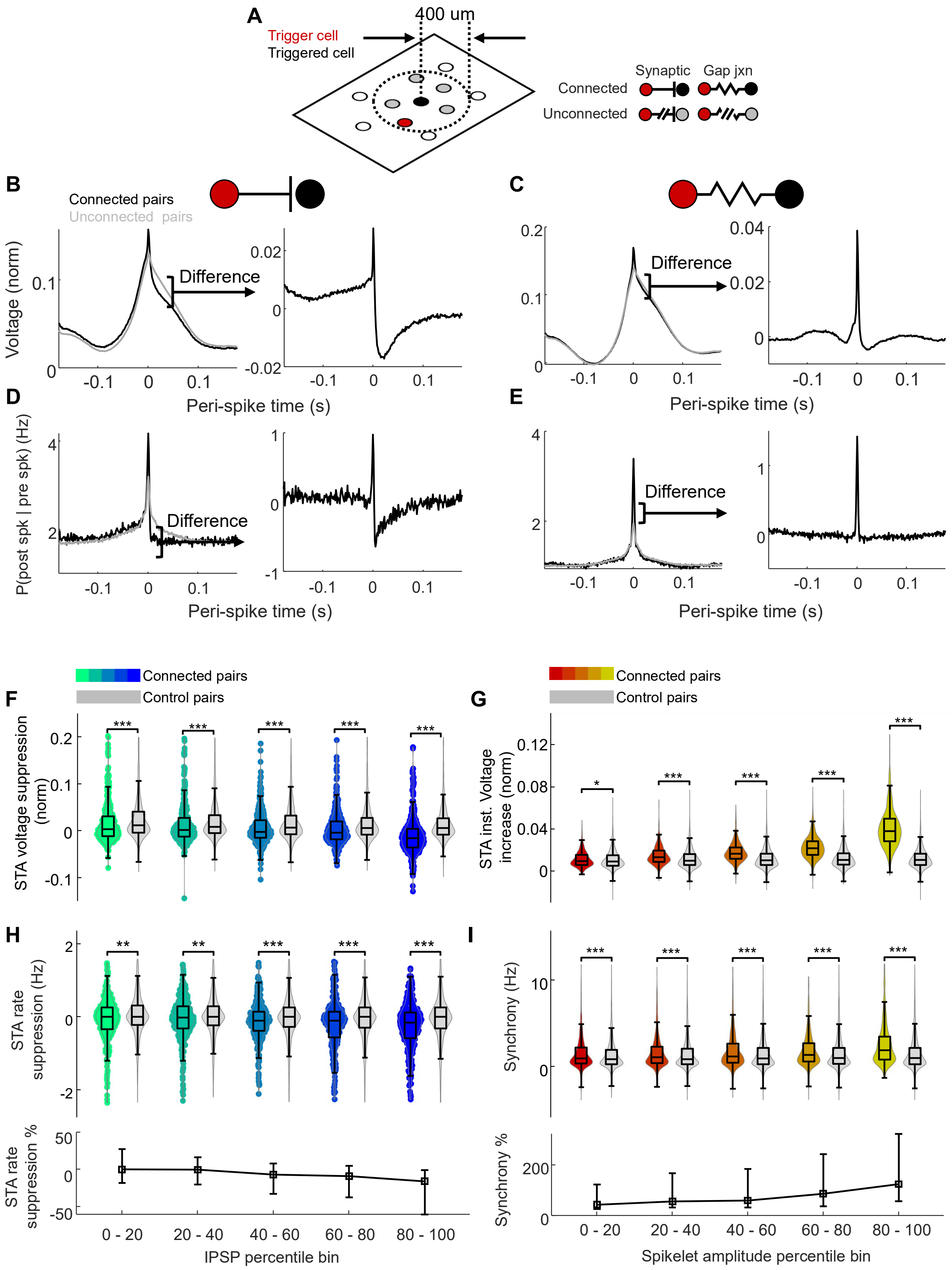
A
400 um
Trigger cell
Triggered cell
Synaptic
Gap jxn
Connected
Unconnected
B
Connected pairs
Unconnected pairs
Voltage (norm)
Difference
Peri-spike time (s)
C
Difference
Peri-spike time (s)
D
P(post spk | pre spk) (Hz)
Difference
Peri-spike time (s)
E
Difference
Peri-spike time (s)
Connected pairs
Control pairs
F
STA voltage suppression (norm)
Connected pairs
Control pairs
G
STA inst. Voltage increase (norm)
H
STA rate suppression (Hz)
STA rate suppression %
0 - 20
20 - 40
40 - 60
60 - 80
80 - 100
IPSP percentile bin
I
Synchrony (Hz)
Synchrony %
Spikelet amplitude percentile bin

**Supplementary Figure 23. Local connectivity shapes spontaneous voltage and spiking dynamics.**

(A) Procedure for selecting un-connected control pairs for cross STA analyses.

(B) Procedure for removing shared subthreshold dynamics from population average cross-STA voltages. Left: black, population average cross STA voltage of connected cell pairs; gray, average cross STA voltage of unconnected control pairs. Right panels: connected minus scaled un-connected waveforms (Methods).

(C) Same as (B) for gap junctionally connected pairs.

(D) Procedure for removing shared dynamics from population average spike cross-correlogram for synaptically connected pairs.

(E) Same as (D) but for gap junctionally connected pairs.

(F) Cross STA hyperpolarization due to synaptic inhibition. Comparison of hyperpolarization between connected vs. unconnected pairs in five quintiles of IPSP amplitudes. Single-sided Wilcoxon rank sum test for statistical significance. *, **, ***, indicate $0.01 < p \leq 0.05$, $0.001 < p \leq 0.01$, and $p \leq 0.001$ for F-I.

(G) Cross STA instantaneous voltage increase due to gap junction. Comparison of instantaneous voltage increase between connected and unconnected pairs in five quintiles of spikelet amplitudes. Single-sided Wilcoxon rank sum test for statistical significance.

(H) Cross STA firing rate suppression due to synaptic inhibition. Top: comparison of firing rate suppression between connected and unconnected pairs in five quintiles of IPSP amplitudes. Single-sided Wilcoxon rank sum test for statistical significance. Bottom: relative firing rate suppression (one minus the ratio of cross STA firing rate before and after a spike) of connected pairs in five quintiles of IPSP amplitudes. Center box marks median, top and bottom bars indicate $75^{th}$ and $25^{th}$ percentile.

(I) Spike synchrony due to gap junctions. Top: comparison of spike synchrony between connected and unconnected pairs in five quintiles of spikelet amplitudes. Single-sided Wilcoxon rank sum test for statistical significance. Bottom: relative synchrony (synchrony divided by baseline spike rate; see Supplementary Fig. 20 for definition) of connected pairs in five quintiles of spikelet amplitudes. Center box marks median, top and bottom bars indicate $75^{th}$ and $25^{th}$ percentile.

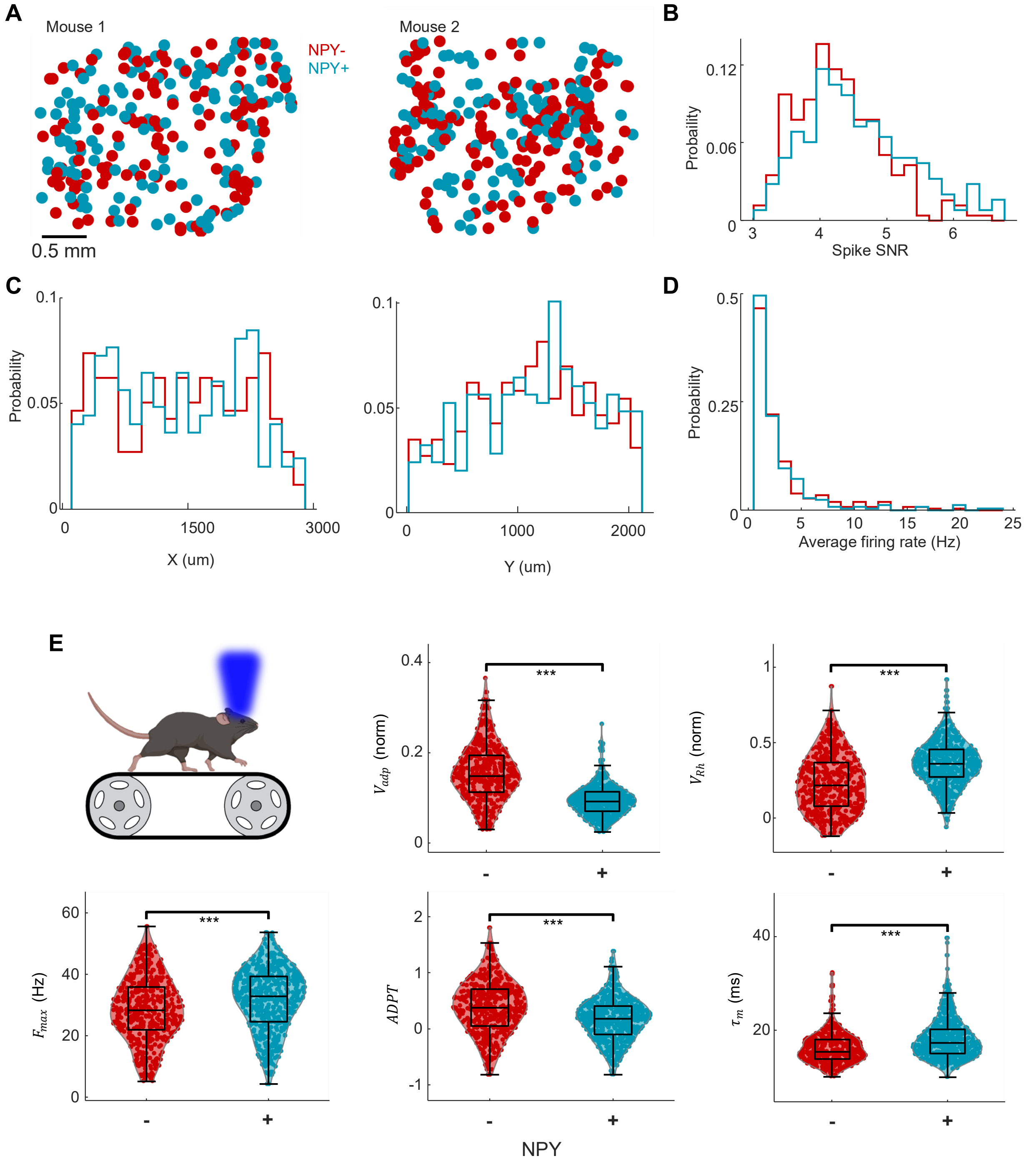


**Supplementary Figure 24. Comparison of intrinsic excitability of NPY$^{(+)}$ vs. NPY$^{(-)}$ subtypes.**

(A) Coordinates of NPY$^{(+)}$ and NPY$^{(-)}$ cells for two mice. For all panels: Red: NPY$^{(-)}$, cyan: NPY$^{(+)}$. n = 257 NPY$^{(-)}$ cells, n = 248 NPY$^{(+)}$ cells.

(B) Spike SNR distributions.

(C) Cell coordinate distributions.

(D) Average spontaneous firing rate distributions.

(E) Cross-subtype comparison of five intrinsic properties listed in Figure 2. Double sided Wilcoxon rank sum test for statistical significance. *, **, ***, indicate $0.01 < p \leq 0.05$, $0.001 < p \leq 0.01$, and $p \leq 0.001$.

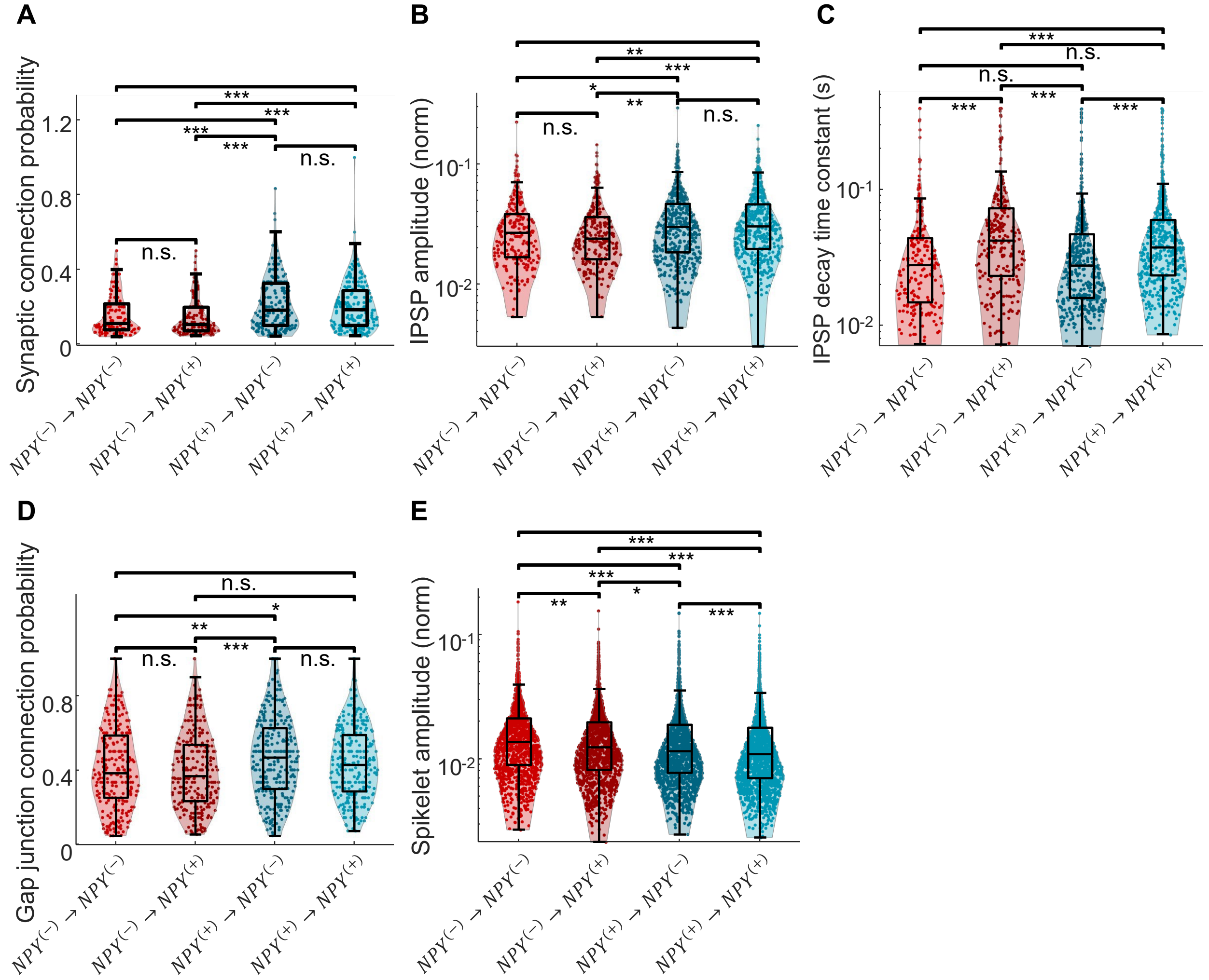


**Supplementary Figure 25. Comparison of network connectivity between NPY$^{(+)}$ vs. NPY$^{(-)}$ subtypes.**

(A) Comparison of uni-directional synaptic connection probability across subtypes. n = 126 $(-) \rightarrow (-)$, 134 $(-) \rightarrow (+)$, 193 $(+) \rightarrow (-)$, 202 $(+) \rightarrow (+)$, N = 2 mice. Double sided Wilcoxon rank sum test for statistical significance. *, **, ***, indicate $0.01 < p \leq 0.05$, $0.001 < p \leq 0.01$, and $p \leq 0.001$ for A – E.

(B) Comparison of uni-directional synaptic connection strength across subtypes. n = 229 $(-) \rightarrow (-)$, 261 $(-) \rightarrow (+)$, 479 $(+) \rightarrow (-)$, 498 $(+) \rightarrow (+)$ pairs, N = 2 mice. Double sided Wilcoxon rank sum test for statistical significance.

(C) Comparison of uni-directional IPSP decay time constant across subtypes. n = 212 (−) → (−), 237 (−) → (+), 434 (+) → (−), 462 (+) → (+) pairs, N = 2 mice. Double sided Wilcoxon rank sum test for statistical significance.

(D) Comparison of uni-directional gap junction connection probability across subtypes. n = 253 (−) → (−), 255 (−) → (+), 297 (+) → (−), 291 (+) → (+) pre-junction cells, N = 2 mice. Double sided Wilcoxon rank sum test for statistical significance.

(E) Comparison of uni-directional gap junction connection strength across subtypes. n = 1222 (−) → (−), 1327 (−) → (+), 1571 (+) → (−), 1632 (+) → (+) pairs, N = 2 mice. Double sided Wilcoxon rank sum test for statistical significance.

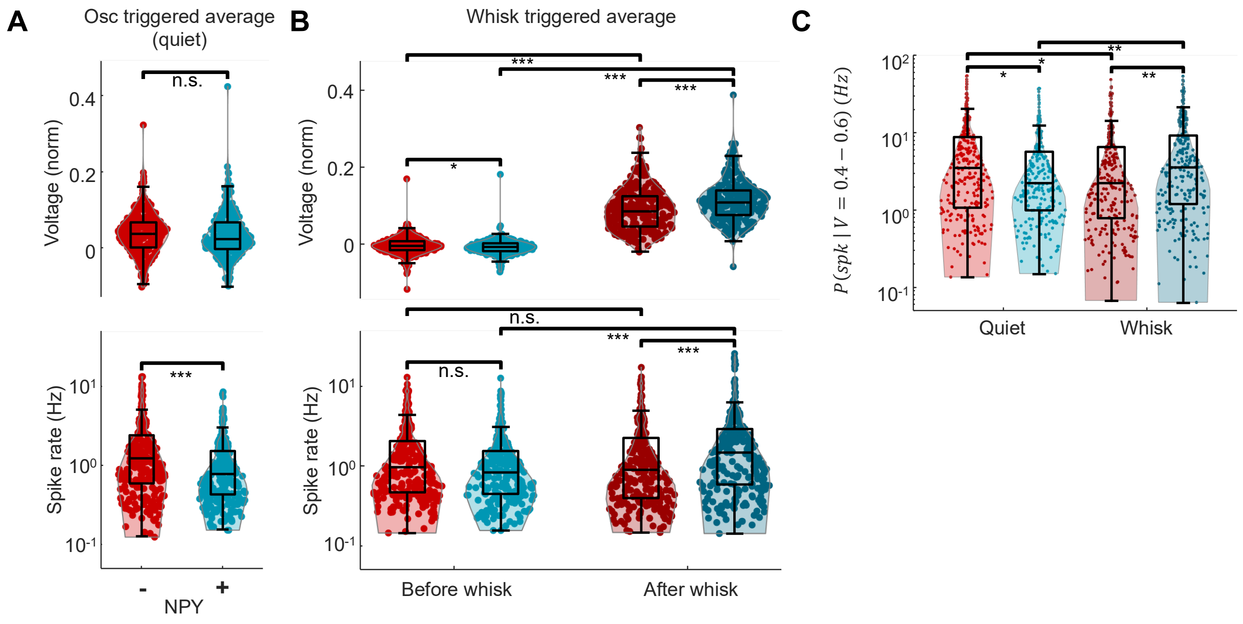


**Supplementary Figure 26. Differential modulation by brain state of excitability of NPY$^{(+)}$ vs. NPY$^{(-)}$ subtypes.**

(A) We calculated the mean subthreshold voltage (top) and spike rate (bottom), triggered off peaks in the quiet-state subthreshold oscillations. Each point represents one cell. n = 257 NPY$^{(-)}$, and 248 NPY$^{(+)}$ cells. Double-sided Wilcoxon rank sum test for statistical significance. *, **, ***, indicate $0.01 < p \leq 0.05$, $0.001 < p \leq 0.01$, and $p \leq 0.001$ for A – C. NPY$^{(-)}$ cells spiked at a higher rate during quiet-state subthreshold oscillations.

(B) We calculated the mean subthreshold voltage (top) and spike rate (bottom) over a 640-ms window before and after the onset of whisking bouts. Each point represents one cell. n = 257 NPY$^{(-)}$ (red), and 248 NPY$^{(+)}$ (cyan) cells. Double-sided Wilcoxon rank sum test for statistical significance.

(C) Conditional spike rate for subthreshold voltages between 0.4 and 0.6, during quiet and whisking epochs. Each point represents one cell. n = 257 NPY$^{(-)}$ (red), and 248 NPY$^{(+)}$ (cyan) cells. Double-sided Wilcoxon rank sum test for statistical significance. In the quiet state the NPY$^{(-)}$ cells spiked faster and in the whisking state the NPY$^{(+)}$ cells spiked faster.
